# Whole-genome 3D architectural screen reveals modulators of brain DNA structure

**DOI:** 10.64898/2026.04.15.718501

**Authors:** Bibudha Parasar, Achuthan Raja Venkatesh, Jonathan Perera, Lucas Sosnick, Siavash Moghadami, Yunji Seo, Jenny Shi, Lynette Chan, Satoshi Takenawa, Tetsuya Akiyama, Odilia Sianto, Takeshi Uenaka, Angela Hadjipanayis, Marius Wernig, Aaron D. Gitler, Longzhi Tan

## Abstract

Three-dimensional (3D) genome architecture is the foundation of gene regulation, and plays a critical role in normal physiology and disease. However, our understanding of its biochemical determinants has long been limited by technology: imaging-based screens only profile a small number of loci, while sequencing-based studies rarely exceed 100 samples or conditions. Here we present “in-plate chromosome conformation capture” (Plate-C), a high-throughput, cost-effective platform that profiles thousands of whole-genome architectures in a day. Plate-C enabled the first chemical screen for whole-genome structural changes—profiling 2,956 samples from 834 conditions across 5 neuronal and glial types, accompanied by 6,081 single cells using “easy diploid chromosome conformation capture” (Easy Dip-C) and 200,893 single-cell transcriptomes. We discovered that diverse, dose/time-dependent, and cell type/species-specific modes of DNA structural changes can be rapidly induced by manipulating epigenetic (HDAC, BET), metabolic (mTOR), proteostatic (UPR), developmental (GSK3/Wnt, Hedgehog), immune (cGAS/STING), and neurotransmission pathways. To validate our finding in vivo, we demonstrated in newborn mice that HDAC inhibition drives brain-wide genome rewiring within hours, highly correlated with changes in vitro and inducing a latent structural and transcriptional state orthogonal to normal differentiation. By enabling massively parallel profiling of whole-genome structures, Plate-C paves the way for systematic discovery of DNA folding principles to better understand and engineer the human genome in 3D.

## Introduction

The three-dimensional (3D) organization of the genome plays a critical role in both normal physiology such as smell^1–4^, learning^5,6^, immunity^7^, morphogenesis^8^, and aging^9,10^, and diseases such as autism^11^, schizophrenia^12^, Alzheimer’s^13,14^, and cancers^15^. Genome architecture controls gene expression by spatially organizing DNA across multiple length scales—from chromosome territories^16,17^ and A/B compartments^18^ to domains^19^ and loops^20,21^, thereby orchestrating transcriptional programs and shaping cellular identity. Despite the importance of genome architecture, its fundamental biochemical principles remain poorly understood across biological conditions, in part due to challenges in systematically interrogating how DNA is restructured in response to diverse perturbations.

This technological and knowledge gap is especially profound in complex systems such as the brain, where a large variety of postmitotic and cycling cell types constantly respond to myriad biochemical signals through epigenetic, metabolic, proteostatic, developmental, immune, and neurotransmission pathways. Recently, we and others showed that the brain genome architecture undergoes extensive, cell type–specific remodeling across development, aging, and other cellular transitions^9,10,14,22^. However, how these 3D genome structural programs respond to external perturbations, such as pharmacological interventions targeting epigenetic, metabolic, and other signaling pathways, and how this response differs between doses, times, drug combinations, cell types, and species remain largely unknown. To map the perturbation landscape of DNA structure across neuronal and glial types, a scalable, rapid, sensitive, and cost-effective screening platform is needed to resolve whole-genome architectural changes across a broad range of physiologically and clinically relevant treatments.

To date, no methods can profile whole-genome 3D architecture at a scale compatible with high-throughput screening (**Supplementary Table 1**). Imaging-based methods have high throughput, but can only probe a small fraction of the genome. For example, HiDRO^23^ and HIPMap^24,25^ screened hundreds to thousands of conditions but only measured 2–3 loci or gross nuclear morphology. Perturb-tracing^26^ analyzed ~100 conditions but only measured ~30 loci along a single chromosome. Sequencing-based methods, such as various versions of Hi-C^18,20,27,28^ and Micro-C^29^, offer genome-wide coverage, but are not scalable due to prohibitive cost and labor associated with biotin pull-down and separate library preparation. For example, method-development papers rarely profiled more than 10 biological conditions. Even among data-generation papers that applied existing methods, only a few^30–32^ reached ~80.

To overcome this technological barrier, we developed “in-plate chromosome conformation capture” (Plate-C), a highly multiplex Hi-C platform that rapidly profiled over an order of magnitude more conditions and samples than existing sequencing studies, without compromising data quality. Applying Plate-C to neurons and glia, we discovered that diverse biochemical pathways can rewire DNA structure within hours or days, in a locus-, dose-, time-, cell type–, and species-dependent manner (**Supplementary Tables 2–3**). We validated our finding in vivo with single-cell profiling, revealing chemically induced genome architectural states orthogonal to normal differentiation, and providing a structural mechanism for our companion paper’s^33^ discovery that chemical reprogramming enhances neuronal resilience.

### Plate-C measures whole-genome architecture at scale

In Plate-C, cells are cultured in 96- or 384-well plates and receive diverse biological or pharmacological perturbations. Following treatment, cells are processed directly in-plate through an optimized workflow that integrates Hi-C (fixation, digestion, and ligation) and library preparation (lysis, transposition, and barcoded amplification) with minimal washes or transfers and without biotin pull-down (**Fig. 1a**). This design minimizes sample loss, batch variability, reagent cost, and handling time, enabling massively parallel generation of thousands of whole-genome contact maps in 24 h, at a low cost of $4 per sample (reagents) and $2 per million contacts (sequencing) (**Extended Data Fig. 1a**). After sequencing all samples for large-scale screening, selected conditions (hits) can be extracted and sequenced deeper to enhance map resolution. This optimized workflow also improved downstream single-cell analysis^34^, which we termed “easy diploid chromosome conformation capture” (Easy Dip-C) (**Fig. 1a**).

**Fig. 1.**
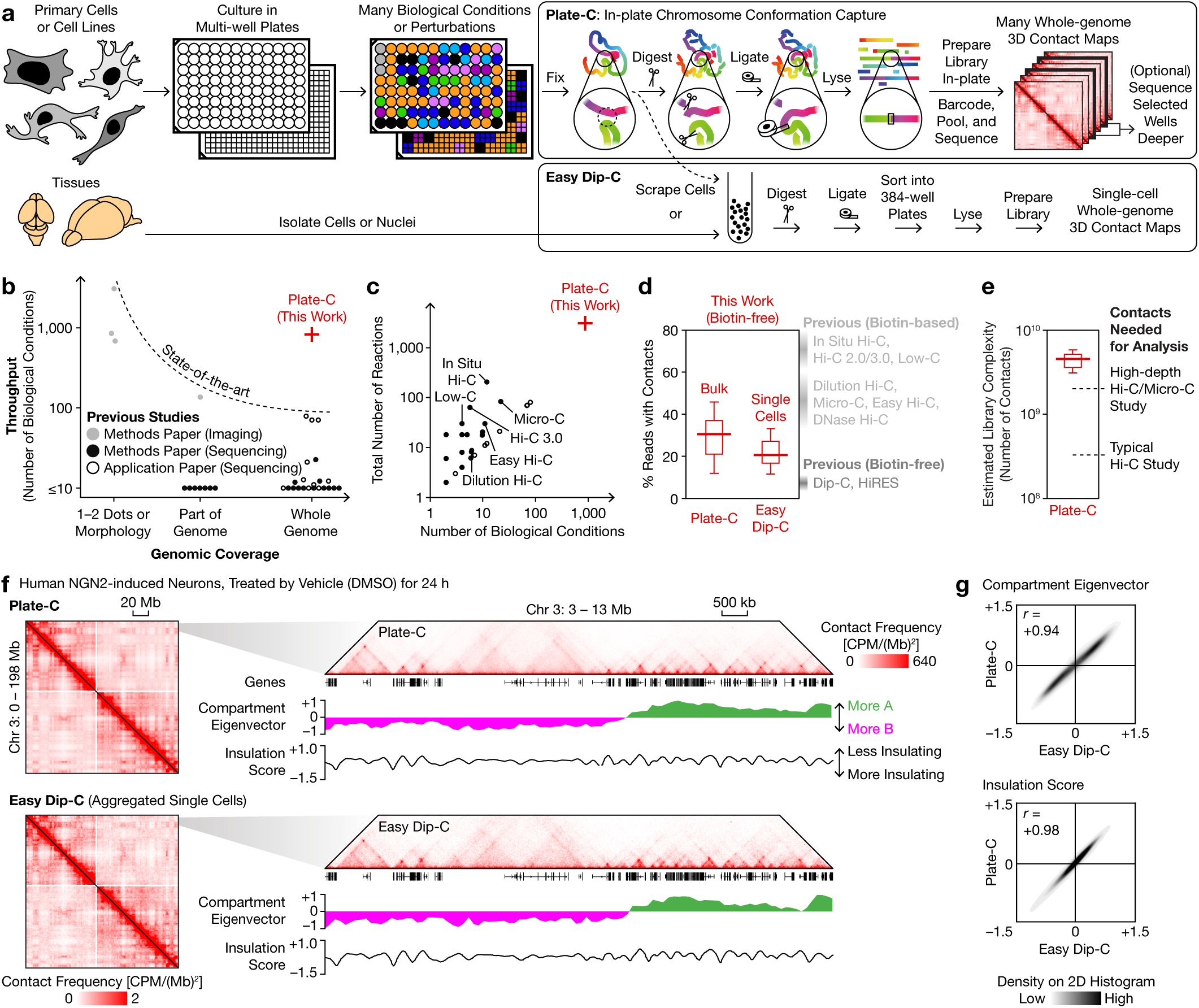
Plate-C measures whole-genome architecture at scale. **a**, Schematic of Plate-C for high-throughput in-plate chromosome conformation capture across diverse biological conditions and of the companion method, Easy Dip-C, for single-cell 3D genome profiling. **b**, Comparison of genomic coverage and throughput between Plate-C and existing 3D genome studies. **c**, Comparison of numbers of biological conditions and samples between Plate-C and existing sequencing-based studies. **d**, Distribution of percentages of Plate-C and Easy Dip-C reads that contained contacts. **e**, Distribution of estimated library complexity of deeply sequenced Plate-C samples. **f**, Representative whole-chromosome (1-Mb bins) and local (25-kb) contact maps of human NGN2 neurons, measured by Plate-C and by Easy Dip-C, with compartment eigenvector (100-kb) and insulation score (25-kb) profiles. **g**, Comparison of compartment eigenvector and insulation score profiles showed high correlation between Plate-C and Easy Dip-C. Boxes in **d, e** show quartiles; whiskers show 1.5× interquartile range (IQR). CPM: contacts per million. *r*: Pearson correlation coefficient.

We systematically benchmarked Plate-C against existing methods for genomic coverage, throughput, and data quality. While existing technologies face a trade-off between coverage (sequencing-based) and throughput (imaging-based), Plate-C uniquely combines genome-wide coverage with scalability to nearly a thousand biological conditions (**Fig. 1b**), filling a longstanding technological gap and achieving over an order of magnitude higher throughput than previous sequencing studies including in situ Hi-C^20,30^, Micro-C^29^, Hi-C 3.0^28^, and Easy Hi-C^35^ (**Fig. 1c**). Despite omitting biotin pull-down, a substantial fraction of Plate-C and Easy Dip-C reads (30% and 20%, respectively) contained contacts, far more efficient than previous biotin-free methods such as Dip-C^2,22,34^ and HiRES^36^ (~5%) (**Fig. 1d, Extended Data Fig. 1b**). While this fraction is lower than that of biotin-based methods (>40%), the additional sequencing cost is offset by eliminating costly biotin-associated reagents and labor, especially for large-scales screens and as sequencing cost continues to decline.

Importantly, performing Hi-C in-plate did not compromise data quality. Despite having only 1–100 k cells, each Plate-C sample yielded a high library complexity of billions of contacts (**Fig. 1e**). Direct comparison between Plate-C (in plate) and aggregated Easy Dip-C (which performs Hi-C steps in suspension) data demonstrated faithful capture of genome organization at all levels including global and local contact maps (**Fig. 1f**), A/B compartment eigenvectors, and insulation scores (**Fig. 1g**). Together, these data showed that Plate-C accurately measures whole-genome architecture at scale, providing a high-throughput platform with rich phenotypic readouts for systematic and quantitative interrogation of DNA organization across diverse perturbations.

### Diverse pathways drive multi-scale genome remodeling

To quantify how each perturbation remodels each level of genome organization, we devised a statistical framework that identifies significant changes across 7 architectural attributes, from coarse to fine scales: the extent of chromosome intermingling, the distribution of contact distances, the overall strength of A/B compartmentalization, the difference in strengths between compartments A and B (A–B difference), the locus-level spatial CpG–based A/B compartment (scA/B; originally defined for single cells) profile^2,9,22,34^, the overall strength of domain boundaries, and the overall strength of loops (**Fig. 2a**). Special care was taken in defining each attribute to ensure robustness against sequence depths (**Methods, Extended Data Fig. 1c**).

**Fig. 2.**
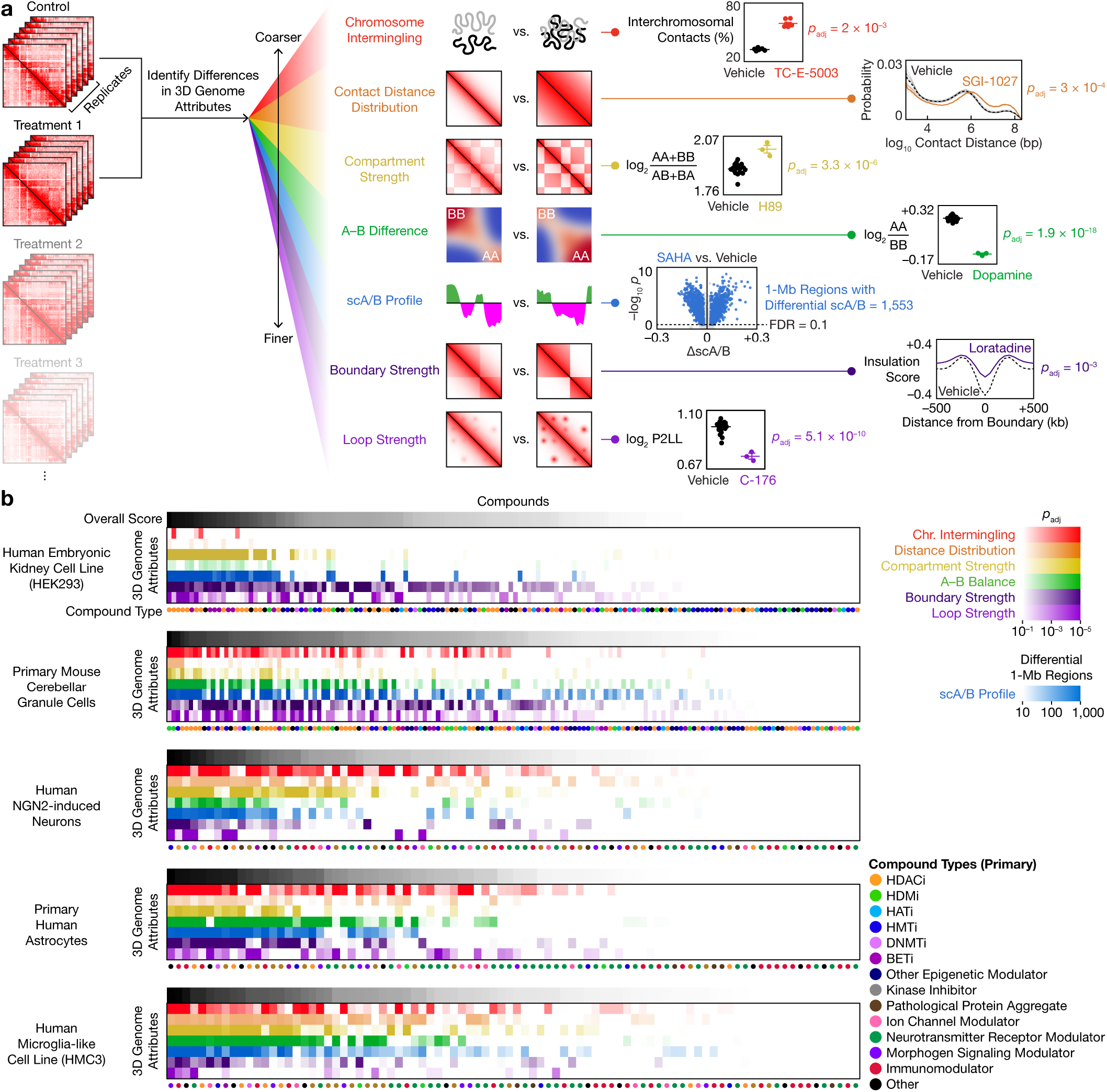
Systematic quantification of multi-scale genome restructuring across diverse perturbations and cell types. **a**, Schematic of the statistical framework to analyze changes across 7 levels of genome organization, with representative contact maps and quantifications. **b**, Summary of genome architectural responses across compounds and cell types. Compounds are sorted by descending overall changes in each cell type. *p* values are from two-sided, equal-variance *t*-tests; *p*_adj_ and FDR values are from BH correction. AA, BB, AB: strengths of A–A, B–B, A–B interactions. P2LL: peak to lower left ratio. HDACi: histone deacetylase inhibitor. HDMi: histone demethylase inhibitor. HATi: histone acetyltransferase inhibitor. HMTi: histone methyltransferase inhibitor. DNMTi: DNA methyltransferase inhibitor. BETi: BET inhibitor.

We applied this framework to examine how subsets of 229 compounds or proteins targeting diverse cellular pathways (**Supplementary Table 4**), including many FDA-approved and preclinical drugs, may rewire genome architecture in 5 neuronal and glial cell types: the human embryonic kidney neuronal cell line HEK293, primary mouse cerebellar granule cells^37^, human NGN2-induced neurons^38^, primary human astrocytes, and the human microglial cell line HMC3 (**Fig. 2b**). This dataset, which encompasses 2,956 genome-wide contact maps from 834 biological conditions, presents the first chemical screen for changes in whole-genome 3D architecture.

We discovered that many compounds—including modulators of epigenetic enzymes, kinases, ion channels, neurotransmitter receptors, morphogen signaling, and immune response—significantly altered one or more of the 7 architectural attributes in at least one cell type (**Fig. 2b**). Most of these compounds were not previously known to alter genome architecture; the few that were^39–41^ had not been characterized in postmitotic or neural cells. Our finding highlights Plate-C’s capability to identify new modulators of DNA structure, and suggests that in both cycling and postmitotic cells, genome architecture could serve as an integrative hub that is dynamically remodeled by diverse biochemical pathways, providing a structural basis for pathway-responsive gene regulation.

### Convergent and divergent modes of restructuring

As a proof of concept for identifying structural convergence among diverse perturbations, we treated HEK293 cells with 157 compounds targeting various epigenetic pathways for 24 h. Principal component analysis (PCA) of genome-wide scA/B profiles revealed distinct modes of chemically induced genome restructuring (**Extended Data Fig. 1c**). Two of these scA/B-based clusters, corresponding respectively to histone deacetylase (HDAC) inhibitors such as TSA and SAHA and to BET inhibitors such as JQ1 and I-BET151, shared common characteristics such as increased long-range and interchromosomal contacts and weakened compartmentalization, yet remained separable by PCA. A third cluster, corresponding to certain histone demethylase (HDM) and methyltransferase (HMT) inhibitors, sirtuin inhibitors, and nucleoside analogs, led to decreased long-range contacts. These and other smaller clusters (**Extended Data Figs. 1d–f**) highlight Plate-C’s capability to capture convergent architectural signatures from a large number of perturbations, whereas previous studies only examined a few^39–41^. We confirmed selected hits in human induced pluripotent stem cells (iPSCs) (**Extended Data Figs. 2a–c**).

We next turned to postmitotic cells, which are free of cell-cycle effects yet had not been studied in 3D genome screens. We applied the 157 compounds to primary mouse cerebellar granule cells—the most abundant neurons of the brain—for 72 h. PCA and uniform manifold approximation and projection (UMAP) of scA/B profiles (**Fig. 3a, Extended Data Fig. 2d**) revealed a complex landscape of architectural responses, with 5 prominent clusters: a continuum of HDAC inhibitors that could be divided into 3 clusters based on magnitudes of scA/B changes, a mechanistically diverse cluster that included the redox-active natural compound plumbagin and several epigenetic inhibitors, and a cluster containing BET inhibitors and the antihistamine cyproheptadine (**Fig. 3b, Supplementary Table 5**).

**Fig. 3.**
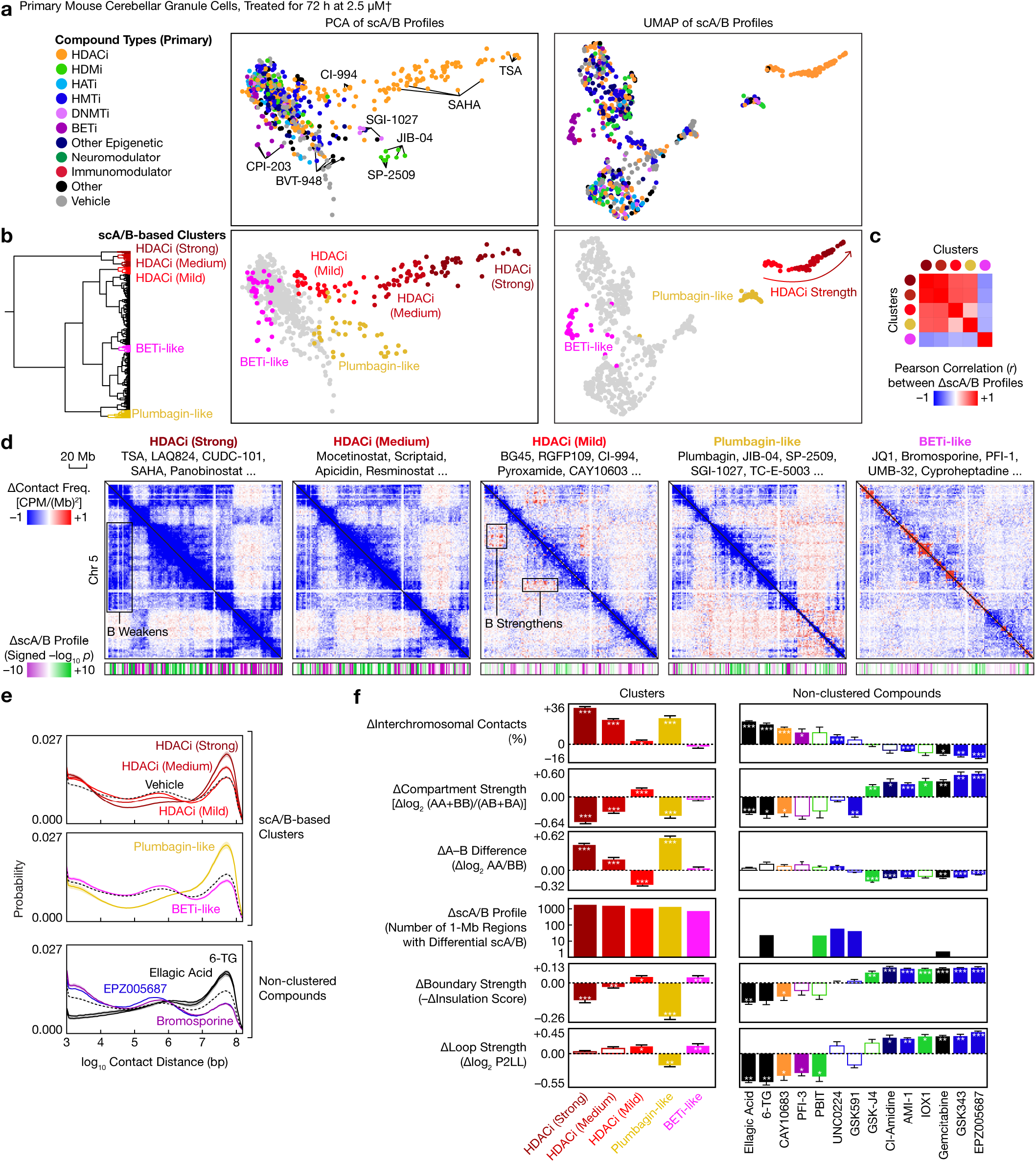
Convergent and divergent modes of chemically induced genome restructuring in primary mouse neurons. **a**, PCA and UMAP of scA/B profiles of primary mouse cerebellar granule cells treated with 157 compounds targeting various epigenetic pathways. **b**, Hierarchical clustering of scA/B profiles identified 5 modes of genome restructuring. **c**, Correlation between the 5 modes of scA/B changes. **d**, Differential chromatin contact maps (1-Mb bins) of the 5 modes, with representative compound names and differential scA/B tracks (1-Mb; *p* values from two-sided, equal-variance *t*-tests). **e**, Distribution of contact distances (mean ± s.e.m.) of the 5 clusters and of selected non-clustered compounds (colored by compound type). **f**, Changes in various architectural attributes (mean ± s.e.m.) produced by the 5 clusters and by selected non-clustered compounds. *p*_adj_ values are from two-sided, equal-variance *t*-tests with BH correction. †: Actual concentrations might be lower because of potential degradation in solution. Hollow: not significant. *: *p*_adj_ < 0.05. **: *p*_adj_ < 0.01. ***: *p*_adj_ < 0.001.

The 5 treatment clusters exhibited distinct modes of neuronal genome rewiring (**Figs. 3c–f, Extended Data Figs. 2e–i**). The strong and medium HDAC inhibitor clusters and the plumbagin-like cluster increased very-long-range (>10 Mb) and interchromosomal contacts, weakened compartmentalization especially in compartment B, and weakened domain boundaries—consistent with global chromatin decompaction and mixing. The strong and medium clusters primarily differed by magnitudes of changes, while the plumbagin-like cluster additionally weakened loops.

In contrast, the mild HDAC inhibitor cluster, consisting primarily of class-specific inhibitors such as CI-994 and RGFP109, had much smaller effects on contact distances and chromosome intermingling but, unlike the strong and medium clusters, strengthened compartmentalization especially in the most B-like regions, and also tended to strengthen boundaries and loops (**Figs. 3d–f**). This surprising finding that HDAC inhibitors can paradoxically reinforce heterochromatin interactions in mouse neurons echoes a recent report in mouse embryonic stem cells (mESCs)^39^ despite differences in cell types, concentrations, duration, and class-specificity—suggesting a general phenomenon across postmitotic and cycling mouse cells.

Finally, the BETi-like cluster acted largely orthogonal to the other 4 clusters, and tended to strengthen both boundaries and loops (**Figs. 3d–f**). Besides the 5 clusters, we observed genome restructuring driven by other compounds such as certain nucleoside analogs (6-TG and gemcitabine) and EZH2 inhibitors (GSK343 and EPZ005687, both of which strengthened compartments, boundaries, and loops) (**Figs. 3e–f**), highlighting the diverse structural plasticity of the postmitotic genome and providing a structural basis for these pathways’ functional roles in neural differentiation^42,43^.

### Architectural dose curves, time courses, and drug interactions

Motivated by the distinct but graded effects of different HDAC inhibitors along a common continuum (**Fig. 3**), we asked whether the architectural effects of a given compound could vary with dose, time, or the presence of other compounds. First, we treated granule cells for 72 h with two compounds— TSA (strong HDAC inhibitor cluster) and the HDM inhibitor JIB-04 (plumbagin-like cluster)—with a broad concentration range spanning 5–6 orders of magnitudes (**Figs. 4a–c**). For each compound alone, we charted a genome architectural dose–response curve (**Fig. 4b, Extended Data Fig. 3a**). Similar to the continuum formed by different HDAC inhibitors, both TSA and JIB-04 produced a continuum of effects as concentration increased: scA/B changes increased in magnitude, whereas most other architectural attributes switched directions.

**Fig. 4.**
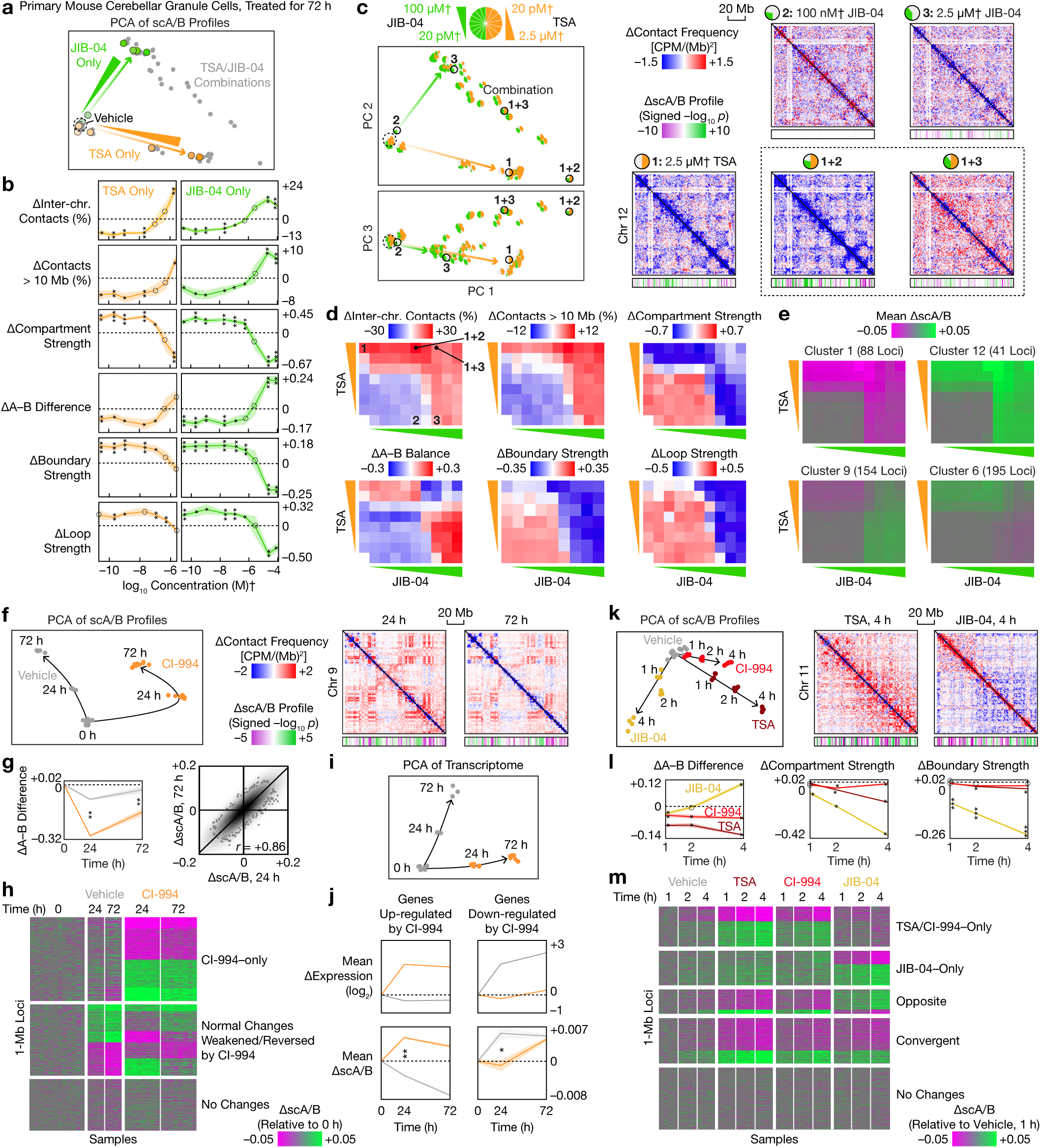
Dose-dependent, time-dependent, and combinatorial perturbation of genome architecture. **a**, PCA of scA/B profiles of primary granule cells treated by 20 pM–2.5 mM† TSA, 20 pM–100 mM† JIB-04, or their combinations for 72 h. For visual clarity, only centroids of the 4 replicates per condition were shown. **b**, Architectural dose–response curves showing changes in various genome architectural attributes (mean ± s.e.m.) produced by each compound alone. **c**, Architectural drug–drug interactions visualized on the PCA and with representative differential contact maps (2.5-Mb bins) and differential scA/B tracks (1-Mb; *p* values from two-sided, equal-variance *t*-tests). **d**, Changes in genome architectural attributes produced by different combinations of TSA and JIB concentrations. **e**, Mean scA/B changes of selected *k*-means clusters of 1-Mb loci, produced by different combinations of TSA and JIB concentrations. **f**, Left: PCA of scA/B time courses of granule cells treated with 2.5 μM† CI-994 for 0, 24, and 72 h. Right: Differential chromatin contact maps (2.5-Mb) and differential scA/B tracks (1-Mb). **g**, Left: Changes in A–B difference (mean ± s.e.m.) at 0, 24, and 72 h. Right: Comparison of CI-994–induced scA/B changes between 24 and 72 h. **h**, *k*-means clustering of 1-Mb loci based on patterns of scA/B changes at 0, 24, and 72 h. **i**, PCA of bulk RNA-seq of CI-994–treated granule cells at 0, 24, and 72 h. **j**, Mean gene expression and scA/B trajectories (mean ± s.e.m.) of genes up- and down-regulated by CI-994 at 24 h. **k**, Left: PCA of scA/B time courses of granule cells treated with 2.5 μM TSA, CI-994, or JIB-04 for 1–4 h. Right: Representative differential chromatin contact maps (2.5-Mb) and differential scA/B tracks (1-Mb). **l**, Changes in selected genome architectural attributes (mean ± s.e.m.) at 1–4 h. **m**, *k*-means clustering of 1-Mb loci based on patterns of scA/B changes at 1–4 h. †: Actual concentrations might be lower because of potential degradation in solution. *p*_adj_ values are from two-sided, equal-variance *t*-tests with BH correction. Circle: not significant. *: *p*_adj_ < 0.05. **: *p*_adj_ < 0.01. ***: *p*_adj_ < 0.001.

When the two compounds were combined, we observed diverse forms of architectural drug–drug interactions (**Figs. 4c–d**). For example, a moderate concentration of JIB-04, which had weak effects on its own, potentiated TSA–driven genome remodeling (**Fig. 4d, Extended Data Fig. 3b**). In contrast, high concentrations of both compounds combined produced weaker TSA-driven and JIB-04–driven effects than either treatment alone. This dose-dependent interaction could be explained by JIB-04’s preferential inhibition of H3K4 demethylases (KDM5s) at lower concentrations^44^, which can synergize with HDAC inhibition, while higher JIB-04 concentrations also inhibit H3K9 demethylases. In both cases, the combined perturbation additionally induced structural changes along PC 3, which were absent from either treatment alone (**Fig. 4c, Extended Data Figs. 3e**). We further showed that different genomic loci integrated signals from the two compounds differently (**Fig. 4e, Extended Data Figs. 3c–d**).

Second, we charted an architectural time course by treating granule cells for 0, 24, and 72 h with CI-994 (mild HDAC inhibitor cluster) and measuring both genome architecture and transcriptome (**Fig. 4f**, **Supplementary Table 6**). In the absence of perturbation, the baseline differentiation trajectory in vitro partially recapitulated early postnatal architectural maturation in vivo^9^ in the first 24 h but diverged by 72 h, with opposite changes in contact distances and no correlation in scA/B changes (**Extended Data Figs. 3f, 9k**). HDAC inhibition rapidly redirected this trajectory through massive remodeling of scA/B profiles and specific strengthening of compartment B, both of which plateaued by 24 h (**Figs. 4f–g**). At the locus level, HDAC inhibition both introduced new changes orthogonal to normal differentiation, and weakened or reverted normal differentiation (**Fig. 4h**). These scA/B changes correlated with gene expression changes, especially at 24 h (**Figs. 4i–j, Supplementary Table 7**).

Finally, we examined the onset of genome restructuring by treating granule cells for 1, 2, and 4 h with 3 compounds: TSA, CI-994, and JIB-04, each representing a different cluster at 72 h (**Fig. 4k**). All 3 compounds rapidly remodeled neuronal genome architecture within 1 h, suggesting that their structural effects are direct rather than downstream of transcriptional changes. While the two HDAC inhibitors produced distinct effects at 72 h (**Fig. 3**), their effects at 1–4 h were similar, including selective strengthening of compartment B and correlated scA/B changes (**Figs. 4l–m**). Together, these findings demonstrate that genome architecture is dynamically rewired in a dose- and time-dependent, locus-specific manner that integrates different biochemical signals, providing a holistic view beyond previous studies^39–41^.

### Human neuronal genome senses many biochemical signals

We next expanded beyond epigenetic pathways and mouse neurons, applying ~90 compounds or proteins—including modulators of diverse, non-epigenetic pathways and selected hits from the 157 epigenetic compounds—to 3 widely used human brain–related cell types: NGN2-induced neurons^38^, primary astrocytes, and HMC3 (**Fig. 2b, Supplementary Table 8**). These compounds cover a broad range of cellular pathways including metabolism (such as mTOR and PI3K inhibitors), other kinases (such as PKA and GSK3^23^ inhibitors), morphogen signaling (such as BMP and Hedgehog inhibitors), neurotransmitter receptors (such as neurotransmitters and modulators of AMPA, NMDA, GABA, and serotonin receptors), ion channels (such as 4-AP, KCl, and TTX), proteostasis (such as tunicamycin), immune activation [such as cGAS/STING inhibitors, antihistamines, cytokines, LPS, and poly(I:C)], and pathological protein aggregates (such as pTau and Aβ). We used well-established concentrations to minimize toxicity and profiled genome architecture at 24 h, and additionally at 2 h for some experiments.

We found that NGN2 neurons remodeled genome architecture in response to diverse biochemical pathways (**Figs. 5a–c, Extended Data Figs. 5a–d**). Epigenetic compounds such as HDAC inhibitors (including TSA, CI-994, and valproic acid), the DNA methyltransferase (DNMT) inhibitor SGI-1027, and the PRMT1 inhibitor TC-E-5003 potently drove multi-level DNA restructuring, supporting the universality of our findings in HEK and granule cells. Beyond epigenetic pathways, NGN2 neurons consistently responded to the mTOR inhibitor Torin-1, the antihistamine loratadine, the cGAS/STING inhibitor H-151, the ER stress/UPR inducer tunicamycin, the GSK3 inhibitor SB-216763, and the neurotransmitter norepinephrine, each of which altered one or more architectural attributes at 24 h. Among the protein aggregates, neurofibrillary tangles (NFTs, or pTau) also tended to remodel neuronal genome architecture (**Extended Data Fig. 5a**). These findings suggest that postmitotic human neurons reorganize genome architecture based on metabolic, stress, and signaling states.

**Fig. 5.**
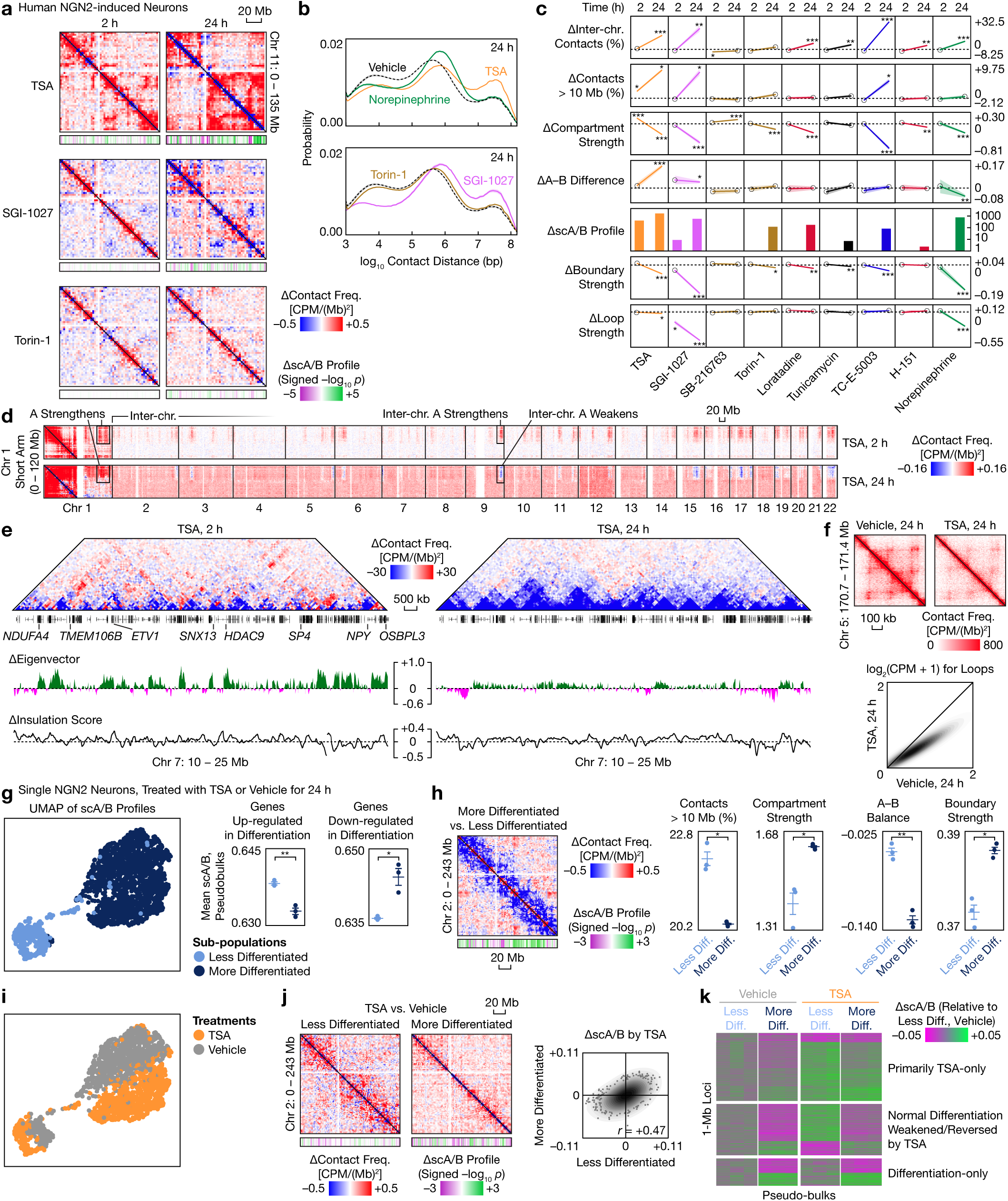
Genome architecture of human neurons responds to diverse biochemical signals. **a**, Differential contact maps (2.5-Mb bins) of NGN2 neurons treated with by selected epigenetic and non-epigenetic compounds, with differential scA/B tracks (1-Mb; *p* values from two-sided, equal-variance *t*-tests). **b**, Distribution of contact distances (mean ± s.e.m.) for selected compounds (colored by compound type) at 24 h. **c**, Time courses of changes in genome architectural attributes (mean ± s.e.m.) produced by selected compounds (colored by compound types). *p*_adj_ values are from two-sided, equal-variance *t*-tests with BH correction. Circle: not significant. *: *p*_adj_ < 0.05. **: *p*_adj_ < 0.01. ***: *p*_adj_ < 0.001. **d**, Intra- and interchromosomal differential contact maps (2.5-Mb) of the short arm of Chromosome 1 in NGN2 neurons treated by 100 nM TSA for 2 and 24 h. **e**, Representative local differential contact maps (100-kb) produced by TSA at 2 and 24 h, with compartment eigenvector (25-kb) and insulation score (25-kb) profiles. **f**, Top: Representative local contact maps (25-kb) showing global attenuation of loops by TSA at 24 h. Bottom: Comparison of loop contact frequencies between TSA and vehicle at 24 h. **g**, Left: UMAP of single-cell scA/B profiles of NGN2 neurons treated with TSA or vehicle for 24 h, colored by structural subtypes. Right: Comparison of pseudobulk scA/B (mean ± s.e.m.) of genes up- and downregulated over normal differentiation between the two subtypes (vehicle only). **h**, Left: Differential contact map (2.5-Mb) between the more and less differentiated subtypes (vehicle only), with differential scA/B tracks (1-Mb). Right: Comparison of selected genome architectural attributes between the two subtypes. **i**, UMAP in **g** but colored by treatment. **j**, Left: Differential contact maps (2.5-Mb) of TSA in the two neuronal subtypes, with differential scA/B tracks (1-Mb). Right: Comparison of TSA-induced scA/B changes between the two subtypes. **k**, *k*-means clustering of 1-Mb loci based on patterns of pseudobulk scA/B changes. *p* values in **g–j** are from two-sided, equal-variance *t*-tests. *: *p* < 0.05. **: *p* < 0.01. ***: *p* < 0.001.

Close examination of HDAC inhibition in NGN2 neurons revealed similarities and differences between species. In the first few hours, TSA selectively strengthened compartment B in mouse granule cells (**Figs. 3f, 4l**) and mESCs^39^, but tended to favor compartment A in NGN2 neurons, especially for ultra-long-range and interchromosomal contacts (**Figs. 5a, 5c–d**), echoing a recent preprint on compartment eigenvector distribution shifting toward A in the human leukemia cell line HAP1^45^. At 24 h, while this A– B difference and the overall extent of chromosome intermingling increased in NGN2 neurons (**Fig. 5c**), interchromosomal A–A contacts were selectively reduced (**Fig. 5d, Extended Data Fig. 6a**). These species differences hint toward a fundamental difference in chromatin regulation between humans and mice^46^. Deeper sequencing further visualized transient, local genome rewiring around key neuronal genes such as *TMEM106B, ETV1*, and *NDUFA4* (**Fig. 5e, Extended Data Fig. 5e**) and global weakening of domain boundaries and loops at 24 h (**Figs. 5e–f**).

We confirmed our finding at single-cell resolution using Easy Dip-C. NGN2 neurons are a heterogeneous population of transcriptionally diverse cells^47^ but had not been characterized by single-cell Hi-C. In the absence of perturbation, neurons formed two structural subtypes, which we identified as more and less differentiated cells, respectively, by correlating scA/B with published transcriptome^47^ (**Fig. 5g**). This finding establishes a baseline differentiation pseudotime for NGN2 neurons, where the two subtypes differ by many architectural attributes (**Fig. 5h**). We confirmed that HDAC inhibition remodeled both subtypes, with higher magnitudes in less differentiated cells and without altering subtype abundances (**Figs. 5i–k**). At the locus level, HDAC inhibition both induced orthogonal changes and tended to weaken or revert differentiation pseudotime (**Fig. 5k**), consistent with our finding in granule cells (**Fig. 4**) and our companion paper’s^48^ discovery on gene expression in NGN2 neurons—providing a structural mechanism for how HDAC inhibition induces neuronal reprogramming and resilience.

### Glia and neurons remodel genome differently

To understand the cell-type specificity of chemically induced genome restructuring, we compared how the 3 human brain–related cell types responded to perturbations (**Fig. 6a**). Epigenetic compounds rapidly remodeled all cell types, with generally correlated scA/B changes but notable cell-type differences. For example, 24 h of HDAC inhibition selectively weakened compartment A in the glial cells but compartment B in NGN2 neurons, potentially caused by biochemical differences between proliferating and postmitotic cells (**Extended Data Fig. 7h**).

**Fig. 6.**
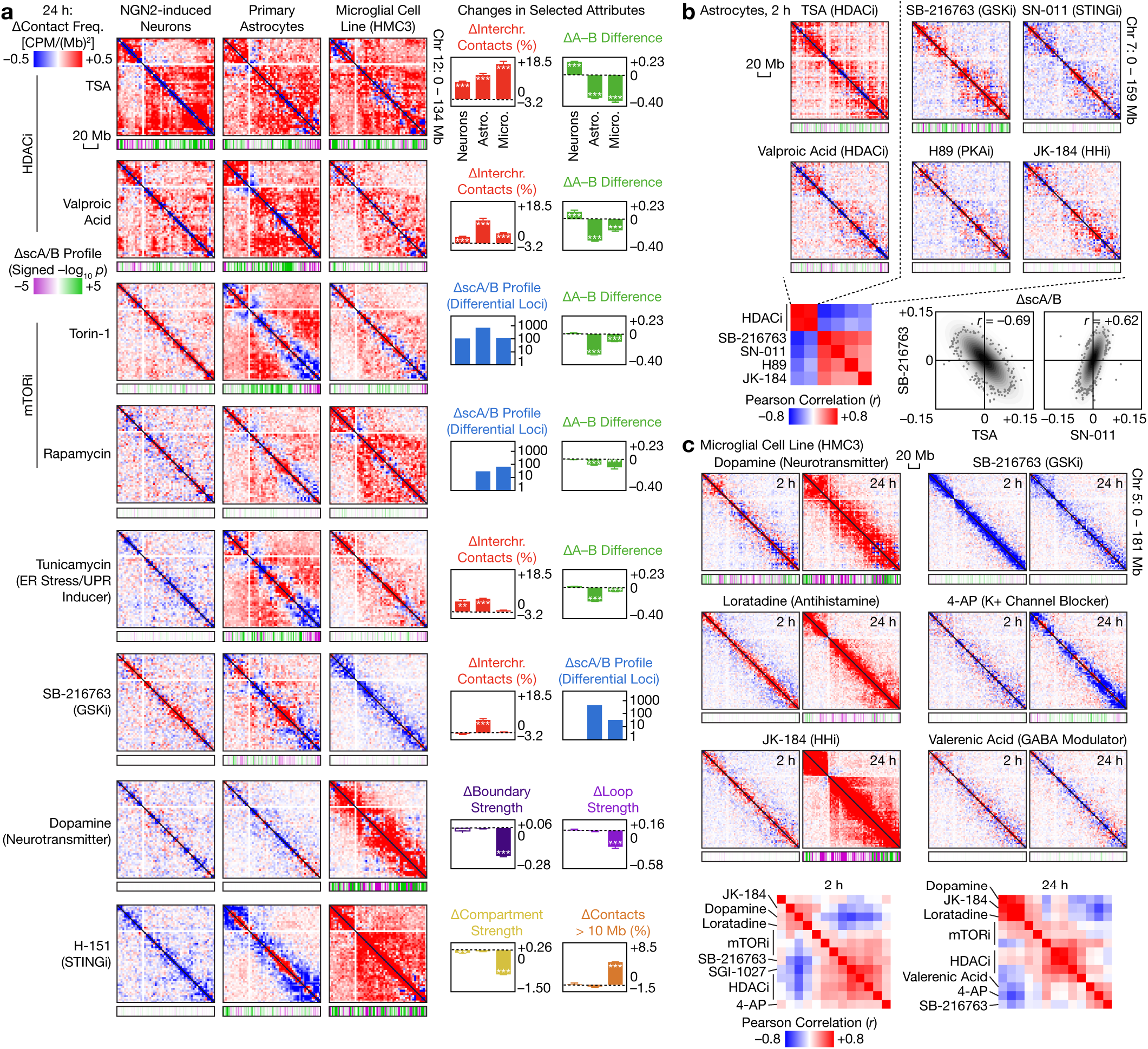
Cell type–specific genome architectural responses to a wide range of perturbations. **a**, Comparison of differential contact maps (2.5-Mb bins), with differential scA/B tracks (1-Mb; *p* values from two-sided, equal-variance *t*-tests), and selected genome architectural attributes between 3 human brain–related cell types, shown for selected compounds at 24 h. **b**, Top: Differential contact maps (2.5-Mb bins) of astrocytes treated by selected compounds for 2 h, with differential scA/B tracks (1-Mb). Bottom: Correlation between scA/B changes produced by these compounds, with representative comparisons visualized by scatter plots. **c**, Top: Differential contact maps (2.5-Mb bins) of HMC3 cells treated by selected compounds for 2 and 24 h, with differential scA/B tracks (1-Mb). Bottom: Correlation between scA/B changes produced by these compounds at 2 and 24 h. *p*_adj_ values are from two-sided, equal-variance *t*-tests with BH correction. Hollow: not significant. *: *p*_adj_ < 0.05. **: *p*_adj_ < 0.01. ***: *p*_adj_ < 0.001.

Genome architecture of primary human astrocytes and HMC3 cells responded to a wider range of perturbations than that of NGN2 neurons (**Fig. 6a**). For example, while mTOR and cGAS/STING inhibition restructured all 3 cell types, astrocytes and HMC3 cells responded to a broader set of such inhibitors including rapamycin and PQR-620 for mTOR, and C-176 and SN-011 for cGAS/STING (**Fig. 6, Extended Data Figs. 6–8**). The glial cells also consistently responded to more pathways than neurons, including PKA (inhibited by H89), Hedgehog (inhibited by JK-184), and dopamine signaling. Between the glial types, ER stress/UPR and GSK3/Wnt inhibition preferentially restructured astrocytes, while cGAS/STING inhibition and dopamine signaling preferentially restructured HMC3 cells—consistent with recent reports on cGAS/STING’s role in age-related microglial inflammation^49^ and dopamine-induced formation of extracellular traps (ETosis) in microglia^50^.

Many epigenetic and non-epigenetic perturbations restructured the glial cells at both 2 and 24 h, allowing us to analyze how their effects changed and converged or diverged over time. In astrocytes, two opposing modes emerged at 2 h, exhibiting anticorrelation in scA/B changes between HDAC inhibitors and various signaling inhibitors (GSK3/Wnt, cGAS/STING, Hedgehog, and PKA) (**Fig. 6b, Extended Data Figs. 6a–f**). This antagonism vanished by 24 h, when these perturbations became largely positively correlated with each other and with other perturbations such as mTOR inhibition and ER stress. HMC3 cells also exhibited two opposing modes at 2 h: epigenetic compounds (HDAC and DNMT inhibitors), and 3 signaling compounds (dopamine, loratadine, and JK-184) (**Fig. 7, Extended Data Figs. 7a–g**). Both modes persisted at 24 h and remained uncorrelated with each other, while a new group (the potassium channel blocker 4-AP, the GABA receptor modulator valerenic acid, and SB-216763) emerged.

**Fig. 7.**
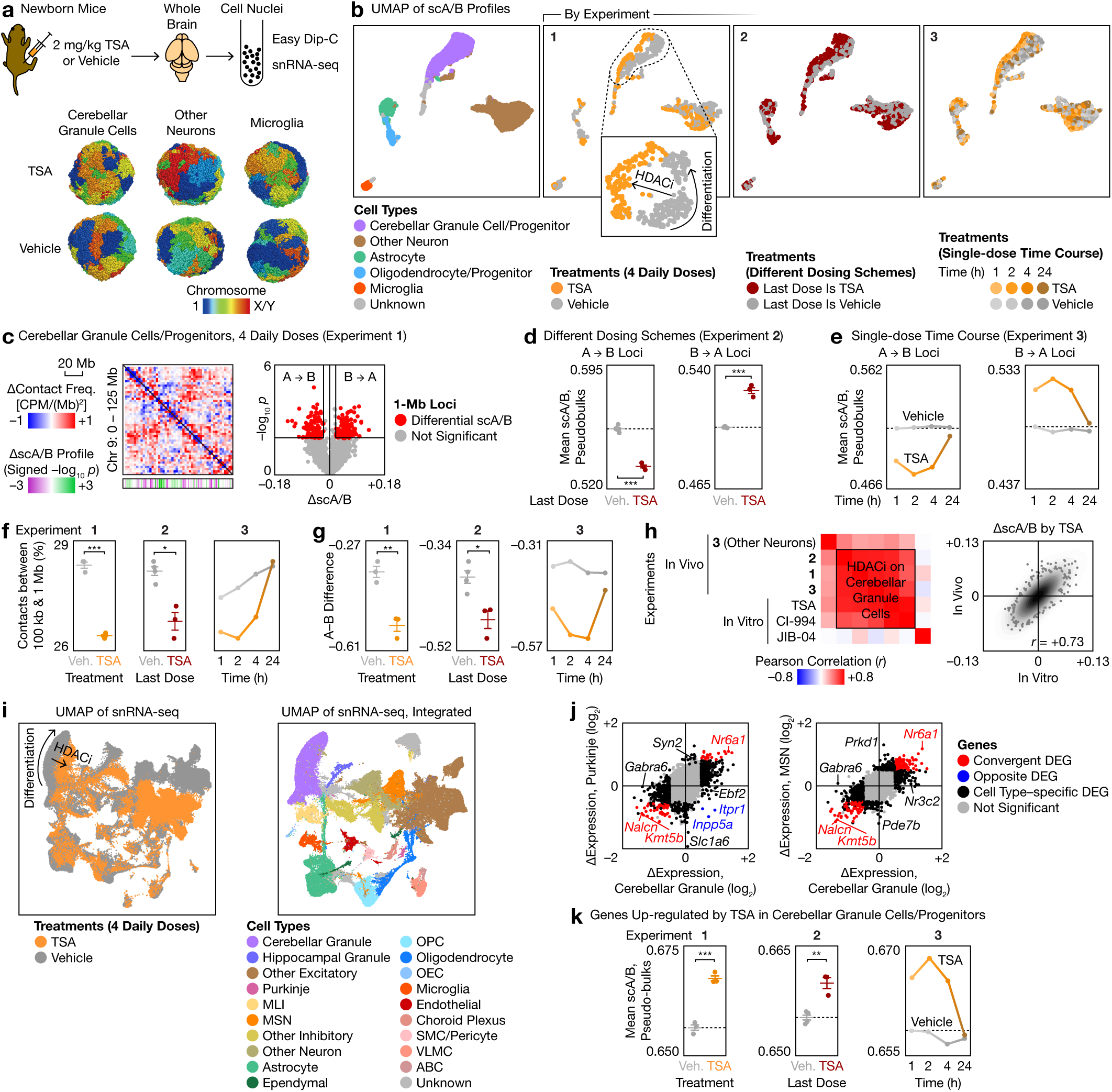
Rapid and transient chemical remodeling of brain genome architecture drives widespread transcriptomic changes in vivo. **a**, Top: Schematic of in vivo validation of Plate-C predictions in newborn mouse brains. Bottom: Representative chemically induced 3D genome structures resolved by Easy Dip-C. **b**, Combined UMAP of single-cell scA/B profiles from three in vivo 3D genome experiments, colored by cell type and by treatment. **c**, Left: Differential contact maps (2.5-Mb bins) in cerebellar granule cells or progenitors in Experiment 1, with differential scA/B tracks (1-Mb; *p* values from two-sided, equal-variance *t*-tests). Right: Volcano plot identifying 1-Mb loci with differential pseudobulk scA/B analysis. **d**, Mean scA/B value (mean ± s.e.m.) of the differential 1-Mb loci from **c** in Experiment 2. **e**, Mean scA/B time courses of the differential 1-Mb loci from **c** in Experiment 3. **f**, Changes in distribution of contact distances (mean ± s.e.m.) produced by TSA in each experiment. **g**, Changes in A–B difference (mean ± s.e.m.) produced by TSA in each experiment. **h**, Correlation between scA/B changes produced by selected perturbations in vivo and in vitro, with a representative comparison (TSA for 1–4 h in vitro vs. in vivo) visualized by a scatter plot. **i**, Left: Unintegrated UMAP of single-cell transcriptomes showing widespread gene expression changes induced by 4 daily TSA injections. Right: Integrated UMAP to identify brain cell types. **j**, Comparison of gene expression changes between representative cell types. **k**, Mean scA/B (mean ± s.e.m.) of genes upregulated by TSA in each 3D genome experiment. *p* values are from two-sided, equal-variance *t*-tests. *: *p* < 0.05. **: *p* < 0.01. ***: *p* < 0.001.

Similar to the species dependency of HDAC inhibition in neurons (**Figs. 3–5**), we found the architectural effects of certain non-epigenetic pathways to differ between humans and mice. In the human microglial cell line HMC3, neither of the immune stimulants [LPS and poly(I:C)] induced strong genome restructuring (**Fig. 2b**). To investigate this systematically, we treated 4 widely used human and mouse cell lines with two concentrations of LPS and poly(I:C) for 24 h (**Extended Data Figs. 8c–e**). Across the 7 architectural attributes, LPS and poly(I:C) had minimal effects on human microglial (HMC3) and macrophage (differentiated from THP-1) cell lines, but induced pronounced genome restructuring in mouse microglial (BV2) and macrophage (RAW 264.7) lines. Together, our findings suggest that diverse epigenetic and non-epigenetic pathways drive cell type–, time-, and species-dependent genome architectural changes across neurons and glia.

### Plate-C predicts genome restructuring in vivo

To demonstrate that Plate-C data predicts chemically induced genome restructuring in vivo, we validated our finding in newborn mouse brains at single-cell resolution using Easy Dip-C. We chose an HDAC inhibitor, TSA, because it crosses the blood–brain barrier, is widely used in neurological disease models^51–55^ and cancers, yet had not been profiled in vivo by Hi-C. We designed 3 experiments with different dosing and sampling schemes: (1) daily injections between postnatal days (P) 6–9, sampled 2 h later, (2) substituting subsets of the 4 injections with vehicle, and (3) a single injection on P9, sampled 1– 24 h later (**Fig. 7a, Extended Data Fig. 9a**). We profiled single-cell 3D genome architecture of the whole brain, resolving diverse cell types including cerebellar granule cells, other neurons, and various glia (**Fig. 7b**) and generating the first chemically perturbed 3D genome structure (**Fig. 7a**).

Consistent with our finding in vitro (**Figs. 3–4**), HDAC inhibition rapidly rewired brain genome architecture in a cell type–specific manner, producing changes in multiple architectural attributes that closely mirrored those in vitro (**Figs. 7b–h, Extended Data Fig. 9b**). In the first experiment, we identified 206 1-Mb loci with differential scA/B in cerebellar granule cells after 4 daily TSA injections, using stringent pseudobulk analysis (**Fig. 7c**). In the next two experiments, we found these scA/B changes to be largely determined by the most recent injection (**Figs. 7b, 7d**) and, after a single injection, to peak around 1–4 h before returning to baseline by 24 h (**Figs. 7b, 7e**). Similar trends were observed for other architectural attributes such as long-range contacts and selective strengthening of compartment B (**Figs. 7f–g, Extended Data Figs. 9b–c**), suggesting that HDAC inhibition produced transient and largely reversible effects, driven primarily by recent exposure. Different cell types exhibited both shared and cell type–specific scA/B changes (**Fig. 7h, Extended Data Fig. 9**).

Most importantly, despite marked differences between baseline differentiation trajectories in vitro and in vivo (**Extended Data Figs. 3f, 9k**), chemically induced scA/B changes were highly correlated between cultured neurons and the brain at the locus level (**Fig. 7h**)—demonstrating that Plate-C screens are strongly predictive of structural effects in vivo. HDAC inhibition partially mimicked juvenile–to–adult transition in vivo^9^ (*r* = +0.42) (**Extended Data Figs. 4a, 9k**), suggesting that dynamic regulation of endogenous HDAC activity may underlie structural maturation of the brain.

Finally, we examined the functional consequences of this genome restructuring by profiling single-cell transcriptome after 4 daily TSA injections. We observed widespread gene expression changes in all neuronal and glial types (**Fig. 7i**). Using stringent pseudobulk analysis, we identified both shared gene expression changes such as the sodium leak channel *Nalcn* and the nuclear receptor *Nr6a1*, and cell type– specific changes such as GABA receptor subunits *Gabra6/g2* in cerebellar granule cells, the glutamate transporter *Slc1a6* in Purkinje cells, and the phosphodiesterase *Pde7b* in medium spiny neurons (**Fig. 7j, Extended Data Fig. 10d**)—mirroring the cell type–dependency of structural changes. Genes upregulated by HDAC inhibition exhibited an average increase in scA/B, which peaked at 2 h (**Fig. 7k**), suggesting that rapid DNA restructuring could drive chemically induced transcriptional programs. In both gene expression and genome structure, HDAC inhibition induced a combination of changes orthogonal to the baseline differentiation pseudotime and complex changes along it (**Extended Data Figs. 10e–f**). Together, these findings suggest that HDAC activity broadly and dynamically controls genome organization and transcription, explaining its diverse physiological functions at the structural level.

## Discussion

Here we developed a scalable platform for measuring thousands of genome-wide architecture, and systematically interrogated the structural effects of diverse epigenetic and non-epigenetic perturbations— spanning metabolic, proteostatic, developmental, immune, and neural pathways—on neuronal and glial cells. This technology overcame a longstanding tradeoff between genome coverage and throughput, and uncovered fundamental principles of cell type–, species-, pathway-, dose-, time-, and locus-dependent genome rewiring that previous studies could not resolve because they could only measure either a small number of loci or a few biological conditions. Our findings demonstrate that in both dividing and postmitotic cells, genome architecture rapidly integrates signals from diverse biochemical pathways—most of which were not previously known to reshape chromatin organization—and that these effects are well conserved at the locus level between in vitro and in vivo.

Many of the compounds studied here are widely used in both basic neuroscience and clinical or preclinical studies of brain disorders, yet were rarely characterized in the context of genome architecture, particularly in brain-related cell types. For example, HDAC inhibitors such as valproic acid have been used in clinics since the 1960s to treat seizures, bipolar disorder, migraine, and brain tumors, in animal models of Alzheimer’s^51–53^, Huntington’s^56^, Parkinson’s^54^, and brain injury^55^, and in basic research to enhance memory^57^ and neuronal resilience to stress such as in our companion paper^48^. In other studies, BET inhibitors alleviate models of fragile X syndrome^58^, mTOR inhibitors enhance mouse longevity^59^, and cGAS/STING inhibitors alleviates inflammation and degeneration of the aging mouse brain^49^. Our discovery that these compounds rewire neuronal and glial genome architecture within hours to days reveals a new structural mechanism of action that may inform the development of precision 3D genome medicine for brain disorders.

Our observation that chemically induced genome restructuring is highly context-dependent may help explain the cell type–, cell state–, and species-specificity of brain disorders, aging, and treatment. For example, humans and mice differ profoundly in lifespan, metabolic profile, disease susceptibility, and response to interventions such as HDAC and mTOR inhibition; within each species, cell types also differ in developmental and aging trajectory, signaling pathway activity, energetic supply and demand, mitotic state, and reprogramming potential. Our findings that different species and cell types exhibit not only different baseline genome architecture^9,34^, but also different architectural responses to external perturbations, may provide a structural basis for how species and cellular contexts shape longevity, disease vulnerability, and treatment outcomes.

## Supporting information

Supplementary Protocol 1

Supplementary Protocol 2

## Data availability

Raw sequencing reads were deposited to the Sequence Read Archive (SRA) under the accession PRJNA1234645: https://www.ncbi.nlm.nih.gov/bioproject/PRJNA1234645.

Processed data was deposited to Zenodo: https://doi.org/10.5281/zenodo.15021657.

## Code availability

Code was deposited to GitHub:

- https://github.com/3d-genome/plate-c
- https://github.com/tanlongzhi/dip-c

## Acknowledgements

We thank Ritchie Chen (USCF) for help with NGN2 neuron and primary astrocyte culture, Kang Shen for discussion on HDACi and UPR, Tom Clandinin for discussion on metabolic pathways, Joe Wu for help with iPSC culture, Xin Jin (Scripps) for advice on multiplex screens, Wenfei Sun, Justus Kebschull (Johns Hopkins), and Chongyuan Luo (UCLA) for advice on liquid handlers, Steve Corsello and Hawa Racine Thiam for advice on drug screens, Dong Xing (Peking) for advice on Dip-C, Mary Hatten (Rockefeller) and Anne West (Duke) for advice on cerebellar culture, Yao Chen (WashU) for advice on PKA, Andrea Gomez (Berkeley), Liqun Luo, and Rob Malenka for advice on serotonin receptors, Serena Sanulli for advice on genetic screens, Evelyn Wong and Emma Follman for help with experiments, Arima, and Stanford core facilities (Shared FACS Facility, Protein and Nucleic Acid Facility, Genomics, Research Computing).

L.T. was supported by NIH (R35GM162529 and R01HL141371), BWF, Sanofi, Baxter, PSF, McKnight, Rita Allen, Stanford MCHRI Uytengsu-Hamilton, RCSA, Wu Tsai, Knight, and Bio-X. A.D.G. was supported by NIH (R35NS137159). B.P. and S.T. were supported by Barres. A.R.V. was supported by Stanford SGF. S.M. was supported by NSF (1828993) and NIH (T32GM136631), and SGF. Y.S. was supported by Stanford MedScholars. L.C. was supported by NIH (T32GM136568) and NSF (DGE-2146755). T.A. was supported by a grant from the Takeda Science Foundation. T.U. was supported by the Huntington’s Disease Foundation.

## Author contributions

B.P. and L.T. conceived the ideas and designed the experiments. B.P., A.R.V., L.S., Y.S., J.S., and S.T. performed Plate-C–related experiments. T.A., O.S., T.U., M.W., and A.D.G. performed NGN2 culture. B.P., A.R.V., J.P., L.S., S.M., and L.C. analyzed the data. A.H. provided guidance. B.P. and L.T. wrote the manuscript with input from all authors.

