## Supplementary material for "Whole-genome 3D architectural screen reveals modulators of brain DNA structure": Methods and Supplementary Figures

**Supplementary Table 1 | Benchmarking Plate-C against existing studies**

**Supplementary Table 2 | Information about each experiment**

**Supplementary Table 3 | Information about each reaction**

**Supplementary Table 4 | Categorization of each compound or protein**

**Supplementary Table 5 | Differential scA/B analysis of genomic region**

**Supplementary Table 6 | k-means clustering of genomic regions based on scA/B, and of genes based on RNA expression**

**Supplementary Table 7 | Differentially expressed genes in time courses of primary neuronal cultures and in vivo**

**Supplementary Table 8 | Master table for attribute changes across all Plate-C experiments**

**Supplementary Protocol 1 | Step-by-step protocol for Plate-C**

**Supplementary Protocol 2 | Step-by-step protocol for Easy Dip-C**

### Methods

Cell culture:

*Human embryonic kidney cells*

293T-derived 293AAV (293AAV) cells were maintained in Dulbecco's Modified Eagle Medium (DMEM, Gibco 10566016) supplemented with 10% fetal bovine serum (FBS, Gibco 10099141) and 1% Antibiotic-Antimycotic (Gibco 15240096). Cells were routinely cultured at 37°C in a humidified incubator containing 5% CO2.

*Mouse primary cerebellar granule cell*

Primary granule cells from mice were cultured according to established protocols[1]. Briefly, culture plates were pre-coated with poly-L-lysine (Sigma P8920, 100 µg/mL), incubated at 37°C for 2 hours, washed with Hank's Balanced Salt Solution (HBSS, Gibco 14025092) supplemented with 1% bovine serum albumin (BSA, Gemini 700-106P). Cerebella from postnatal day 6 (P6) mice were dissected, washed in HBSS-BSA, finely minced, and enzymatically dissociated using 0.025% trypsin (Gibco 15090046). The digestion was terminated by adding DMEM containing 0.1% FBS, 100 µg/ml sodium pyruvate (Gibco 11360070), and 100 µg/ml gentamycin (Sigma G1264). Following trituration, cells were counted and seeded at densities suitable for either 384-well or other culture formats previously coated with poly-L-lysine (100 µg/mL, Sigma P4707), marked as DIV0. After 24 hours (DIV1) in culture, media was changed to include 100 µg/ml gentamycin, 100 µg/ml sodium pyruvate, 0.1% FBS, 10 µM cytosine arabinofuranoside (araC, Sigma C1768), and 10 mM KCl (Invitrogen AM9640G). Specific drugs were added concurrently with this medium change and maintained for designated experimental durations.

*Induced pluripotent stem cells (iPSCs)*

Human induced pluripotent stem cells (iPSCs) were maintained on 6-cm tissue culture plates pre-coated with Matrigel (Corning 356234) under defined culture conditions using StemMACS™ iPS-Brew XF medium (Miltenyi Biotec 130-104-368). Cells were routinely passaged every 4 to 5 days utilizing Gentle Cell Dissociation Reagent (STEMCELL Technologies 100-0485).

*Human NGN2-derived Neurons*

Human H1 embryonic stem cells (hESCs) were maintained in mTeSR plus media (StemCell Technologies, 100-0276) on plates coated with Matrigel (Corning, 354230), and passaged every 3-4 days using ReLeSR (StemCell Technologies, 100-0483). hESCs were differentiated into cortical-like neurons (iNeurons) by overexpressing transcription factor neurogenin 2 (NGN2) driven by a Tet-On induction system in DMEM/F12 media supplemented with 1 ug/mL doxycycline, as previously described[2]. Cells were dissociated using Accutase (StemCell Technologies, 07920) on day 3 of differentiation and replated on plates coated with poly-D-lysine (Thermo Scientific, A3890401) and mouse laminin (Sigma-Aldrich, L2020-1MG) in Neurobasal Medium (Thermo Fisher, 21103049) containing neurotrophic factors, BDNF and GDNF (R&D Systems). Treatments were added on day 10 of differentiation.

*Human microglial cells (HMC3)*

Human microglial clone 3 (HMC3) cells (*ATCC, CRL-3304*) were cultured under standard conditions in Dulbecco's Modified Eagle Medium (DMEM, Gibco 10566016) supplemented with 10% fetal bovine serum (FBS, Gibco 10099141) and 1% Antibiotic-Antimycotic (Gibco 15240096). Cells were maintained at 37°C in a humidified incubator with 5% $CO_2$. Cells were passaged at ~70–80% confluency using 0.05% trypsin-EDTA (Gibco 25300054) and routinely seeded at densities appropriate for downstream assays. For experimental treatments, cells were plated in the desired format (e.g., 96- or 384-well plates) and allowed to adhere overnight prior to compound addition. Drug treatments were performed by replacing culture media with fresh media containing the indicated compounds and maintained for specified durations.

*Human primary astrocytes*
Primary human astrocytes (ScienCell 1800) were cultured according to the manufacturer's guidelines . Cells were maintained in Astrocyte Medium (ScienCell 1801) supplemented with 2% fetal bovine serum (FBS, ScienCell 0010), Astrocyte Growth Supplement (AGS, ScienCell 1852), and 1% penicillin/streptomycin solution (P/S, ScienCell 0503), and incubated at 37°C in a humidified atmosphere with 5% $CO_2$. Culture medium was refreshed the morning after thawing and subsequently replaced every 2–3 days. Once cultures reached ~70% confluency, medium changes were performed every other day until cells reached ~90% confluency.

For passaging, cells were subcultured at 90–95% confluency. Culture vessels were pre-coated with poly-L-lysine (ScienCell 0403, 2 μg/cm²) prior to plating. Cells were rinsed with calcium- and magnesium-free Dulbecco's phosphate-buffered saline (DPBS, ScienCell 0303) and dissociated using 0.05% trypsin-EDTA (T/E, ScienCell 0183). Enzymatic dissociation was monitored microscopically and neutralized using trypsin neutralization solution (TNS, ScienCell 0113) supplemented with fetal bovine serum (FBS, ScienCell 0500). Cells were collected, centrifuged at 1000 rpm for 5 minutes, resuspended in complete medium, and plated onto poly-L-lysine-coated vessels at a density of approximately 5,000 cells/cm². Cells were allowed to adhere overnight prior to experimental treatments.

*Murine microglial cells (BV2)*
BV2 cells (AcceGen ABC-TC212S) were maintained in Dulbecco's Modified Eagle Medium (DMEM, Gibco 10566016) supplemented with 10% fetal bovine serum (FBS, Gibco 10099141) and 1% Antibiotic-Antimycotic (Gibco 15240096). Cells were cultured at 37°C in a humidified incubator with 5% $CO_2$ and passaged at ~70–80% confluency using 0.05% trypsin-EDTA (Gibco 25300054). For experiments, cells were seeded in appropriate culture formats and allowed to adhere overnight prior to treatment. Drug treatments were performed by replacing culture medium with fresh medium containing the indicated compounds for specified durations.

*Human monocytic THP-1 cells (macrophage differentiation)*
THP-1 cells (AcceGen ABC-TC1234) were maintained in Roswell Park Memorial Institute medium (RPMI-1640, Gibco 11875093) supplemented with 10% fetal bovine serum (FBS, Gibco 10099141) and 1% Antibiotic-Antimycotic (Gibco 15240096), and cultured at 37°C with 5% $CO_2$. Cells were maintained in suspension and passaged every 2–3 days to maintain a density between $2 \times 10^5$ and $1 \times 10^6$ cells/mL. For macrophage differentiation, THP-1 cells were seeded and treated with phorbol 12-myristate 13-acetate (PMA, Sigma P8139, 100 nM) for 24 hours, followed by a 24-hour rest period in PMA-free medium prior to experimental treatments. Differentiated cells were then exposed to indicated compounds under specified conditions.

*Murine macrophage-like cells (RAW264.7)*

RAW264.7 cells (AcceGen ABC-TC0934) were cultured in Dulbecco's Modified Eagle Medium (DMEM, Gibco 10566016) supplemented with 10% fetal bovine serum (FBS, Gibco 10099141) and 1% Antibiotic-Antimycotic (Gibco 15240096). Cells were maintained at 37°C in a humidified atmosphere with 5% $CO_2$ and passaged at ~70–80% confluency using gentle scraping, as these cells are sensitive to trypsinization. For experimental assays, cells were plated at appropriate densities and allowed to adhere overnight prior to compound treatment. Media was replaced with fresh medium containing indicated treatments and maintained for specified durations.

Chemical Treatment In Vitro:

*Treatments in 293AAV cells*

Epigenetic drug screening was performed using cultured 293T cells. 293 cells were cultured in two 96 well plates in standard DMEM supplemented with 10% fetal bovine serum (FBS) and 1% penicillin/streptomycin. For screening experiments, cells were seeded and maintained at 37°C in a humidified incubator. Cells were plated to reach approximately 40% confluency by the time of drug treatment. The following day, an Epigenetics Screening Library (Cayman Chemical, catalog #11076), containing 158 epigenetic drugs dissolved at 10 mM in DMSO distributed across two 96 well plates, were thawed at room temperature. Drugs were administered directly to the cells at a final concentration of 2.5 μM. After an additional 24 hours of incubation, when cells had reached approximately 70% confluency, the samples were proceeded for Plate-C assays.

*Treatments in iPSCs*

During passaging (designated as Day 0), cells were seeded in a 96 well plate at a 1:10 dilution ratio to ensure optimal growth density. On Day 1 post-seeding, cultures underwent a half-media exchange, and specified drugs were introduced at this point. Briefly, Trichostatin A, CI-994, JQ1, Pyroxamide, TC-E-5003 was dissolved in DMSO to a stock concentration of 10 mM. Drugs were added to iPSC cells at a final concentration of 2.5 μM. After an additional 24 hours of incubation, when cells had reached approximately 50% confluency, the samples were harvested and processed for Plate-C assays.

*Treatments in cerebellar granule cells*

Mouse primary granule cells were cultured in two 384 well plates at a density of 37,500 cells/well on DIV0. The next day (DIV1) media was replaced with one containing araC and KCl. The Epigenetics Screening Library (Cayman Chemical, catalog #11076), containing 158 epigenetic drugs dissolved at 10 mM in DMSO distributed across two 96 well plates, were thawed at room temperature. Drugs were administered directly to the cells at a final concentration of 2.5 μM. After an additional 72 hours of incubation (DIV4), the samples were proceeded for Plate-C assays.

*Combinatorial treatment of HDACi and HDMi in cerebellar granule cells*

Mouse primary cerebellar granule neurons were plated in two 384-well plates at a density of 37,500 cells per well on DIV0. On the following day (DIV1), the culture medium was replaced with a medium supplemented with AraC and KCl. Trichostatin A (TSA) and JIB-04 were obtained from the Epigenetics Screening Library (Cayman Chemical, catalog #11076). Serial dilutions of TSA were applied across rows (2500, 500, 100, 20, 1, 0.2, 0.02, and 0 nM), while serial dilutions of JIB-04 were applied across columns (100000, 20000, 2500, 500, 100, 20, 1, 0.2, 0.02, and 0 nM), generating a combinatorial dose matrix. Cells were incubated with compounds for 72 hours prior to initiating Plate-C.

*Time course of HDACi treatment in cerebellar granule cells*

Mouse primary granule cells were cultured in three 384 well plates at a density of 37,500 cells/well, and in three 96 well plates at a density of 150k cells/well on DIV0. The next day (DIV1) media was replaced with one containing araC and KCl. CI-994 was dissolved in DMSO to a stock concentration of 10 mM. To understand the effect of CI-994 on granule cells across differentiation, cells were treated with vehicle or CI994 at a final concentration of 2.5 μM. One plate each was processed for plate-c assay on DIV 1, 2 and 4. The 384 well plate was used for plate-C assay and the 96 well plate was used for bulk transcriptomics.

*Fast time course of HDACi and HDMi treatment in cerebellar granule cells*

Mouse primary granule cells were cultured in three 384 well plates at a density of 37,500 cells/well on DIV0. The next day (DIV1) media was replaced with one containing araC and KCl. Trichostatin A, CI-994 and JIB-04 was dissolved in DMSO to a stock concentration of 10 mM. To understand the direct effect of these epigenetics drugs on the genome in granule cells, cells were treated with vehicle (DMSO) or epigenetic drugs at a final concentration of 2.5 μM. One plate each was processed for plate-C assay at 1h, 2h or 4h. Single cells can be sequenced deeper to obtain more contacts.

*Treatments in NGN2-Neuron, astrocytes, and HMC3 microglia for Plate-C and Easy Dip-C*

Individual compounds were obtained from commercial vendors. Compounds were prepared according to the manufacturers' instructions and dissolved or diluted in either DMSO or water as appropriate. Treatments were performed using a half-media exchange strategy, in which half of the existing medium was removed and replaced with fresh medium containing 2× the desired final drug concentration. Plates were gently mixed to ensure uniform distribution of compounds and incubated for either 2 or 24 hours prior to termination of treatment and downstream processing for Plate-C. Trichostatin A treatment was performed for 24 hours prior to Easy Dip-C.

*Treatments in HMC3, Activated THP-1, BV2, RAW264.7*

The four cell lines were treated with either two different concentrations of bacterial lipopolysaccharide (LPS, 100 ng/mL or 1 μg/mL) or polyinosinic:polycytidylic acid (PolyI:C, 10 μg/mL or 100 μg/mL) on a 96 well plate for 24 hours. Cells were processed for Plate-C afterwards.

Animals:

Animals were maintained following the guidelines provided by the U.S. National Research Council's Guide for the Care and Use of Laboratory Animals. All animal protocols were reviewed and approved by the Administrative Panel on Laboratory Animal Care (APLAC) at Stanford University. Mice were housed in groups of up to five per cage in enriched housing conditions, with pregnant mice housed individually and with a consistent 12-hour light and 12-hour dark cycle. All animals were in good health, exhibited normal immune function, and had no previous exposure to experimental procedures, drugs, or testing.

The majority of mice used in these experiments belonged to the inbred C57BL/6J strain, obtained from Jackson Laboratory (stock number: 000664). To facilitate 3D genome modeling using Dip-C, F1 hybrid mice were bred in-house by crossing the CAST/EiJ strain (Jackson Laboratory 000928) with the C57BL/6J strain (Jackson Laboratory 000664).

Chemical treatment in vivo:

*4 daily IP injections of HDACi*
Mice received repeated intraperitoneal injections of 2 mg/Kg TSA (Trichostatin A, 3 mice) or vehicle (DMSO, 3 mice) starting from day 6 (P6). Specifically, animals were administered four consecutive injections, with a 24-hour interval between injections (from P6 to P9). Two hours following the final injection, mice were euthanized, and whole brains were immediately isolated for Easy Dip-C assay. Single cells can be sequenced deeper to obtain more contacts.

*Different dosing schemes*
Mice received intraperitoneal injections of either 2 mg/Kg TSA (Trichostatin A) or vehicle (DMSO) on each day depending on the design scheme starting from day 6 (P6). The day mice received TSA was marked as '+', and DMSO was marked as '–'. With one mouse per injection scheme, the series were '+ + + +', '– + + +', '– – + +', '+ + + –', '+ + – –', '+ – – –', '– – – –' (from P6 to P9). Two hours following the final injection, mice were euthanized, and whole brains were immediately isolated for Easy Dip-C assay. Single cells can be sequenced deeper to obtain more contacts.

*Time course after a single dose*
Mice received one intraperitoneal injection of 2 mg/Kg TSA (Trichostatin A, 4 mice) or vehicle (DMSO, 4 mice) on day 9 (P9). One mouse each receiving TSA or DMSO was sacrificed at 1h, 2h, 4h or at 24 h, and whole brains were immediately isolated for Easy Dip-C assay. Single cells can be sequenced deeper to obtain more contacts.

Plate-C:
We have provided a detailed, step-by-step protocol of Plate-C as **Supplementary Protocol 1**. We have also summarized the method below:

*Fixation*
Media was fully aspirated from cells cultured in either 96- or 384-well plates. Cells were fixed using 2% paraformaldehyde (PFA, EMS 15714) in PBS (Invitrogen 10010023) at room temperature (RT) with shaking (800 rpm, 10 min). The reaction was quenched with 10% BSA (GeminiBio 700-101P) in PBS, followed by centrifugation at 1000 ×g for 5 minutes at 4°C. After supernatant removal, pellets were resuspended in 10% BSA in PBS, incubated at room temperature for 5 minutes, and centrifuged again at 2500 ×g for 5 minutes at 4°C. Supernatants were discarded, and fixed cells can be stored at –80°C.

*Permeabilization, Digestion, and Ligation in 384 well plate*
For a similar protocol in 96-well plate, scale all the numbers by 4-fold. Cells were permeabilized with 5 µL of Plate-C cell-lysis buffer (10 mM Tris–HCl pH 7.5 (Invitrogen 15567-027), 10 mM NaCl (Invitrogen AM9760G), 3 mM $MgCl_2$ (Invitrogen AM9530G) and 0.1% IGEPAL CA-630 (Sigma I8896), and 1% BSA (GeminiBio 700-101P)) on ice-water for 20 minutes, followed by sequential treatments with 5 µL of water and 5 µL of 1% SDS (final 0.33% SDS, Sigma 71736), incubated at 62°C (10 min, 800 rpm). Samples were subsequently incubated with 5 µL of 10% Triton X-100 (final 3.3% Triton X-100, Thermo Scientific 85111) at 37°C (15 min, 800 rpm). 7 µL of digestion mix (1X rCutSmart, 10U MboI (NEB R0147L), 10U NlaIII (NEB R0125L)) was then added, and samples were incubated at 37°C (1 hour, 800 rpm), followed by

enzyme inactivation at 65°C (20 min, 800 rpm). One well was marked as digestion QC. Samples were centrifuged (2500 ×g, 5 min, 4°C), supernatant volume adjusted, and either 23 µL of NEB ligation mix (48.9 (v/v)% NEBNext Ultra II Ligation Master Mix (NEB E7595L), and 1.63 (v/v)% NEBNext Ligation Enhancer (NEB E7595L)) or 20 µL of the Vazyme ligation mix (1X Rapid ligation buffer, 73 U/µL of T4 ligase (Vazyme N103)) added. Reactions proceeded at room temperature for 15–30 minutes with shaking (800 rpm). One well was marked as ligation QC. Post-ligation, plates were centrifuged again (2500 g, 5 min, 4°C), supernatants removed, and plates stored at –80°C for longer term storage. Plates were then lysed with 25 µL of Dip-C lysis buffer (25 mM DTT ((Sigma 646563), 20 mM Tris pH 8.0 (ThermoFisher AM9855G), 0.15% Triton X-100 (Thermo Scientific 85111), 500 nM carrier ssDNA: TCAGGTTTTCCTGAA (IDT), 20 mM NaCl (ThermoFisher AM9760G), 1 mM EDTA (ThermoFisher AM9260G)) at 50°C for 16 hours, followed by 70°C for 20 min). 2 µL of samples were aliquoted into V-bottom 384 plates (Biorad HSP3801) for library amplification and stored at 4°C overnight or –80°C for longer-term storage.

*Quality Control*
Lysates from wells designated for digestion and ligation QC were transferred to DNA-LoBind microcentrifuge tubes. For 384-well plates, lysates from four QC wells were combined. DNA was purified using Zymo columns (D4013) with a fivefold volume of DNA Binding Buffer and eluted into TE buffer (Invitrogen AM9849).

*Whole-genome Amplification and Tn5 Optimization*
Optimal Tn5 concentration was determined by performing whole-genome amplification on cell lysate with varying Tn5 amounts. Briefly, 2 µL of samples were aliquoted into a 8-strip tube (USA Scientific 1402-4780). The lysate was mixed with 8 µL of Transposition Buffer (10% PEG 8000 (Hampton Research HR2-535), 12.5 mM TAPS pH8.5 (Boston Bio Products BB-2375), 6.25 mM $MgCl_2$ (ThermoFisher AM9530G)) containing serial dilutions of Tn5 (TTE Mix V50 from Vazyme TD501-01) transposome at a concentration of 50, 12.5, 6.25, 3.13, 1.6, 0.8, 0.4, 0.2 µL per 850 µL Transposition Buffer). The reactions were proceeded through transposition stop and amplification using Nextera i7 index primer: CAAGCAGAAGACGGCATACGAGATTCGCCTTAGTCTCGTGGGCTCGG and Nextera i5 index primer: AATGATACGGCGACCACCGAGATCTACACTAGATCGCTCGTCGGCAGCGTC, amplified for 7 cycles following the amplification step described below. Samples were purified using DNA Clean & Concentrator-5 (Zymo D4014) following manufacturer's protocol. Purified libraries were run on the Agilent 2100 Bioanalyzer to find the Tn5 concentration that gives the flattest Bioanalyzer profile (a little higher on the long side is the most ideal).

*Transposition*
The 384 well plates containing 2 µL lysates were thawed, mixed with 8 µL of freshly prepared Transposition Mix (Transposition buffer containing optimized Tn5 concentration) from the previous step, and incubated at 55°C for 10 minutes.

*Transposition Stop*
The reaction was stopped by adding Stop Mix (45 mM EDTA (ThermoFisher AM9260G, 300 mM NaCl (ThermoFisher AM9760G), 0.01% Triton X-100 (Sigma 93443), 480 µg/mL Qiagen Protease (Qiagen

19157)), followed by incubation steps at 50°C (40 min) and 70°C (20 min). Plates were stored at 4°C overnight.

*Amplification*

PCR Mix (1X Q5 reaction buffer (NEB M0491S), 1X GC enhancer (NEB M0491S), 538 µM dNTP each (Vazyme P031-02), 5.38 mM $MgCl_2$ (ThermoFisher AM9530G), 448 µg/mL BSA (NEB B9000S), 0.0448 U/µLQ5 polymerase (NEB M0491S)) and Nextera Primer Mix (IDT) were added to each well, vortexed, and subjected to PCR amplification for 7 cycles (incubated at 98°C for 30 s, four cycles of (4°C for 3 min, 72°C for 3 min, 98°C for 20 s, 7 cycles of (98°C for 10 s, 62°C for 1 min, 72°C for 2 min) and a final 72°C for 5 min.). Plates were stored at 4°C overnight or –80°C for long-term storage.

*Purification*

PCR reactions were pooled with DNA Binding Buffer (Zymo D4004-1-L), mixed, and stored at –20°C. DNA purification utilized Zymo columns, eluting DNA into TE buffer. Final library concentrations and fragment lengths were assessed using Qubit and Bioanalyzer. Optimal libraries underwent double-sided size selection (0.45X right, 0.6X left) twice, and yield percentages were recorded to confirm transposition efficiency.

Easy Dip-C:

We have provided a detailed, step-by-step protocol of Easy Dip-C as **Supplementary Protocol 2**. We have also summarized the method below:

*Nuclei isolation from P9 mice*

A chilled 2 mL Dounce homogenizer (Sigma D8938) was used to homogenize tissue in ice-cold Nuclei Isolation Buffer with Triton (250 mM sucrose (Sigma 84097), 25 mM KCl (ThermoFisher AM9640G), 10 mM HEPES pH 7 (ThermoFisher 15630080), 5 mM $MgCl_2$ (ThermoFisher AM9530G), 1 µM DTT (Sigma 646563), 0.1% Triton X-100 (Sigma 93443)) for 20 strokes with the tight pestle B. The homogenate was transferred, and centrifuged at 100 ×g for 8 min at 4°C. Supernatants were discarded, and pellets resuspended in Nuclei Isolation Buffer without Triton, followed by another identical centrifugation. Final pellets were resuspended in 2 mL of Nuclei Isolation Buffer without Triton and filtered through 40 µm filters (Fisherbrand 22-363-547).

*Fixation*

Nuclei were fixed using 2% paraformaldehyde (PFA, EMS 15714) in Nuclei Isolation Buffer without Triton at room temperature (RT) for 10 min. The reaction was quenched with 0.5% BSA (Miltenyi Biotec 130-091-376), followed by centrifugation at 1000 ×g for 5 minutes at 4°C. After supernatant removal, pellets were resuspended in 1% BSA in PBS, incubated at RT for 5 minutes, counted, aliquoted to 250k nuclei per reaction, centrifuged, and fixed pellets can be stored at –80°C.

followed by sequential treatments with 5 µL of water and 5 µL of 1% SDS (final 0.33% SDS, Sigma 71736), incubated at 62°C (10 min, 800 rpm). Samples were subsequently incubated with 5 µL of 10% Triton X-100 (final 3.3% Triton X-100, Thermo Scientific 85111) at 37°C (15 min, 800 rpm). 4 µL of digestion mix (1X rCutSmart, 10U MboI (NEB R0147L), 10U NlaIII (NEB R0125L)) was then added, and samples were incubated at 37°C (1 hour, 800 rpm), followed by enzyme inactivation at 65°C (20 min, 800 rpm). One well was marked as digestion QC. Samples were centrifuged (2500 ×g, 5 min, 4°C), supernatant volume adjusted, and either NEB ligation mix (48.9 (v/v)% NEBNext Ultra II Ligation Master Mix (NEB E7595L), and 1.63 (v/v)% NEBNext Ligation Enhancer (NEB E7595L))

*Permeabilization, Digestion, and Ligation*
Nuclei were resuspended in 20 µL of water and 22 µL of 1% SDS (Sigma 71736), incubated at 62°C (10 min, 800 rpm). Samples were subsequently incubated with 22 µL of 10% Triton X-100 (Thermo Scientific 85111) at 37°C (15 min, 800 rpm). 28 µL of digestion mix (1X rCutSmart, 40U MboI (NEB R0147L), 40U NlaIII (NEB R0125L)) was then added,, and samples were incubated at 37°C (1 hour, 800 rpm), followed by enzyme inactivation at 65°C (20 min, 800 rpm). A 10 µL aliquot was taken out for digestion related QC. Ligation mix (48.9 v/v% NEBNext Ultra II Ligation Master Mix (NEB E7595L), and 1.63 v/v% NEBNext Ligation Enhancer (NEB E7595L)) was added, and reactions proceeded at room temperature for 15 minutes. Another 10 µL was taken out for ligation related QC. Post-ligation, samples were centrifuged again (2500 ×g, 5 min, 4°C), supernatants removed, and ligated nuclei are resuspended in 300 nM in 4′,6-diamidino-2-phenylindole, stored at –80°C until single nuclei FACS sorting.

*Quality Control*
Digestion QC and ligation QC tubes were centrifuged at 1000 ×g for 5 minutes, pellets were resuspended in 4U proteinase K (NEB P8107S) in 100 µL PBS, and lysed at 65C for 1 hour. DNA was purified using Zymo columns (D4013) with a fivefold volume of DNA Binding Buffer and eluted into TE buffer (Invitrogen AM9849). Samples were sent to Bioanalyzr or run on an e-gel to validate successful digestion and ligation.

*Microwell based Fluorescence-Activated Nuclei Sorting and lysis*
Nuclei were then passed through cell strainers (35 µm, Fisher Scientific 08-771-23) and sorted into a 384 well plate containing 1 µL of Dip-C lysis buffer (25 mM DTT ((Sigma 646563), 20 mM Tris pH 8.0 (ThermoFisher AM9855G), 0.15% Triton X-100 (Thermo Scientific 85111), 500 nM carrier ssDNA: TCAGGTTTTCCTGAA (IDT), 20 mM NaCl (ThermoFisher AM9760G), 1 mM EDTA (ThermoFisher AM9260G)) using a 100 µm chip, vortexed, spun down at 500 ×g for 1 minute. Plates were incubated at 50°C for 1 hour, followed by 70°C for 15 minutes to lyse the nuclei, and the plates were stored at –80°C for long-term storage.

*Whole-genome Amplification and Tn5 Optimization*
Optimal Tn5 concentration was determined by performing whole-genome amplification on cell lysate with varying Tn5 amounts. Lysate (2 µL) was mixed with a Transposition Buffer (10% PEG 8000 (Hampton Research HR2-535), 12.5 mM TAPS pH 8.5 (Boston Bio Products BB-2375), 6.25 mM $MgCl_2$ (ThermoFisher AM9530G)) containing serial dilutions of Tn5 (TTE Mix V50 from Vazyme TD501-01) transposome. DNA was quantified (Qubit) and analyzed for fragment distribution (Bioanalyzer/e-gel) to identify optimal Tn5 levels.

*Transposition*
Lysates were thawed, mixed with 4 µL of freshly prepared Transposition Mix (10% PEG 8000 (Hampton Research HR2-535), 12.5 mM TAPS pH8.5 (Boston Bio Products BB-2375), 6.25 mM $MgCl_2$ (ThermoFisher AM9530G)) containing optimized Tn5 concentration, sealed, and incubated (55°C, 10 min).

The reaction was stopped by adding 1 µL Stop Mix (45 mM EDTA (ThermoFisher AM9260G, 300 mM NaCl (ThermoFisher AM9760G), 0.01% Triton X-100 (Sigma 93443), 480 µg/mL Qiagen Protease (Qiagen 19157)), followed by incubation steps at 50°C (40 min) and 70°C (20 min). Plates can be stored at 4°C overnight.

*Amplification*

6 µL of PCR Mix (1X Q5 reaction buffer (NEB M0491S), 1X GC enhancer (NEB M0491S), 538 µM dNTP each (Vazyme P031-02), 5.38 mM $MgCl_2$ (ThermoFisher AM9530G), 448 µg/mL BSA (NEB B9000S), 0.0448 U/µL Q5 polymerase (NEB M0491S)) and 1 µL of 10 bp UDI Nextera Primer Mix (IDT) were added to each well and subjected to PCR amplification for 16 cycles (incubated at 98 °C for 30 s, four cycles of (4°C for 3 min, 72°C for 3 min, 98°C for 20 s, 16 cycles of (98°C for 10 s, 62°C for 1 min, 72°C for 2 min) and a final 72°C for 5 min.). Plates were stored at 4°C overnight or –80°C for long-term storage.

*Purification*

PCR reactions were pooled with DNA Binding Buffer (Zymo D4004-1-L), mixed, and stored at –20°C. DNA purification utilized Zymo columns, eluting DNA into TE buffer. Final library concentrations and fragment lengths were assessed using Qubit and Bioanalyzer. Optimal libraries underwent 0.6X left side size selection once, and the library was sequenced on the Illumina platform Novaseq X-Plus.

*Bulk directional RNA sequencing*

Supernatant was completely removed from each well, gently washed once with 100 µL PBS, and cells were lysed and total RNA was isolated using the Direct-zol RNA Miniprep Kits (Zymo R2053) following manufacturer's protocol. RNA was quantified using the Equalbit RNA HS Assay Kit (Vazyme EQ211) following the manufacturer's protocol for 'Protocol A: Poly(A)-based mRNA enrichment'. 100 ng RNA was processed for Poly(A)-based mRNA enrichment using VAHTS mRNA Capture Beads 2.0 (Vazyme N403), and mRNA was directly used for subsequent library preparation using VAHTS Universal V8 RNA-seq Library Prep Kit for Illumina (Vazyme NR605) following the manufacturer's protocol with the following modifications. 1) RNA was fragmented at 94 C for 5 min to obtain a 200 - 300 bp insert size. 2) 2nd Strand Buffer 2 (with dUTP) was used during second strand cDNA synthesis for strand-specific mRNA libraries. 3) 0.5 µL of VAHTS RNA Adapter-S for illumina was used during adapter ligation to minimize excess adapter in the final library, 4) Option B: For libraries with >200 bp inserts (suitable for mRNA fragmented by incubation at 94°C for 5 min) was used during Purification/ Size-selection Protocol. 5) VAHTS RNA Multiplex Oligos Set 1 - Set 2 for Illumina (Vazyme #N323/N324) was used during the library amplification step, and the library was amplified for 15 cycles. Final library was quantified using Equalbit 1 × dsDNA HS Assay Kit (Vazyme EQ121), and was sequenced on the Illumina platform Novaseq X-Plus.

*Single-cell RNA sequencing*

Mice received repeated intraperitoneal injections of 2 mg/Kg TSA (Trichostatin A) or vehicle control starting from day 6 (P6). Specifically, animals were administered four consecutive injections, with a 24-hour interval between injections (from P6 to P9). Two hours following the final injection, mice were euthanized, and whole brains were immediately isolated for single cell transcriptomics experiments. The collected brain tissues were dissociated in 3 mL of homogenization buffer (10 mM Tris pH 8.0, 5 mM $MgCl_2$, 25 mM KCl, 250 mM sucrose, 1 µM DTT, 0.5x protease inhibitor [cOmplete Protease Inhibitor

Cocktail, MilliporeSigma 11697498001], and 0.2 U/µL RNase inhibitor), 0.1% Triton X-100. Tissue homogenates were filtered through a 100-µm cell strainer, and centrifuged at 1000 ×g for 5 minutes at 4°C. The supernatants were discarded, and cell pellets were gently resuspended in 450 µL of cold homogenization buffer.

For density gradient separation, an equal volume of 50% iodixanol solution (146 mM sucrose, 10 mM Tris pH 8, 5 mM $MgCl_2$, and 50% w/v iodixanol) was mixed gently with the cell suspension, achieving a final iodixanol concentration of 25%. This mixture was carefully layered onto 900 µL of a cold 29% iodixanol solution and centrifuged at 13,500 ×g for 20 minutes at 4°C. After centrifugation, the upper layers were discarded, and the resulting pellet was gently resuspended in approximately 50 µL of EB buffer (Qiagen 19086) supplemented with 0.5% bovine serum albumin (BSA) and 0.2 U/µL RNase inhibitors (Vazyme R301), counted with a hemocytometer, and adjusted to 1500 cells/µL. For each sample, ~20 k nuclei were loaded into a Chromium GEM-X Single Cell 3' Gene Expression Kits v4 following its user guide (CG000731 Rev A).

*Plate-C data analysis*

*Pre-processing*

Plate-C data were pre-processed similarly to Dip-C[3], with the mouse reference mm10 (GRCm38 from GENCODE) and human reference hg19 (the "hs37d5" version) with 2 minor modifications: (1) threshold for removal of PCR duplicates was reduced to "--dup-dist=1" (during the "hickit" step), because bulk data has fewer PCR cycles and artifacts than single cells; (2) contacts that overlapped with public blacklists (https://github.com/Boyle-Lab/Blacklist/blob/master/lists/mm10-blacklist.v2.bed.gz and https://github.com/Boyle-Lab/Blacklist/blob/master/lists/hg19-blacklist.v2.bed.gz) were removed with "hickit.js bedflt."

*Library complexity estimation for contacts*

Library complexity was estimated for each sample by integrating Picard-derived duplication metrics with contact yield from the Plate-C processed contact files. Briefly, the total number of read pairs examined, duplicate read pairs, and the estimated library size were obtained from Picard (EstimateLibraryComplexity). The number of valid contacts was extracted from Plate-C processing. To approximate contact-level library complexity, we scaled the Picard-estimated library size by the fraction of unique read pairs contributing to contacts, calculated as the ratio of total contacts to non-duplicate read pairs (READ_PAIRS_EXAMINED − READ_PAIR_DUPLICATES). The resulting value represents the estimated number of unique contacts in the library. Where available, the total number of sequenced reads was independently computed from raw FASTQ files for reference.

*Contact histogram calculation*

For each sample, distance-dependent contact probability was computed from the pairs.gz file. Only autosomal cis contacts were considered, excluding interactions involving sex chromosomes (chrX/chrY) and all trans contacts. Genomic distances were calculated for each contact pair and filtered to retain interactions ≥1 kb. Distances were transformed to log10 scale and binned using a fixed bin width of 0.05 over the range 1kb onward. For each bin, the number of contacts was counted and normalized by the total number of included cis contacts, yielding a probability distribution of contact frequency as a function of genomic distance.

*Statistical comparison of contact distance distributions*

To assess differences in distance-dependent contact distributions between treatments and vehicle controls, we applied a group-level Kolmogorov–Smirnov (KS) test on empirical cumulative distribution functions (ECDFs). For each treatment, replicate-level histograms were subsampled to a fixed number of contacts per replicate to ensure balanced comparison. ECDFs were computed on a common grid. Statistical significance was assessed using permutation testing, in which replicate labels were randomly shuffled between treatment and vehicle groups, generating a null distribution of KS statistics. Empirical p-values were computed based on the position of the observed KS statistic within this null distribution. To account for multiple hypothesis testing across treatments, p-values were adjusted using the Benjamini–Hochberg false discovery rate (FDR) procedure.

*scA/B analysis*

Plate-C scA/B matrices were analyzed similarly to Dip-C[3] with a minor modification that differential scA/B analysis was performed with 2-sided t-tests, rather than U-tests, with FDR < 1% and $|\Delta scA/B| > 0.02$, because the rank-based U-tests did not perform well with low replicate numbers. The first 10 PCs were used for hierarchical clustering with Ward's method.

*Quantification of overall and relative strengths of chromatin A/B compartmentalization*

First, merged .pairs.gz and .hic files for a given treatment/condition/experiment were prepared from the data corresponding to the individual replicates. Merging of .pairs.gz files for individual/treatment/condition/experiment was done via hickit version r29 and, for merging several individual files, parallel merged batch-sorting was deployed. Conversion from .pairs.gz file to .hic was facilitated by juicer_tools_1.22.01.jar with mm10 and hg19 genomes used for M. musculus and H. sapiens data respectively. Conversion from .hic to .mcool was done via HiCExplorer v3.6[4]. A resolution of 1 Mb (using 100kb does not change our conclusions) was used for compartment calling; balancing was performed via cooler v0.9.3 iterative correction and eigenvector decomposition (ICE) normalization with the default parameters (parameters: mad_max = 5, min_nnz = 10, min_count = 0) as also described in the compartment calling tutorial (https://cooltools.readthedocs.io/en/latest/notebooks/compartments_and_saddles.html) to correct for low coverage regions; the files were also filtered to exclude mitochondrial chromosomes (chrMT, chrM). Eigenvector decomposition was performed using cooltools.eigs_cis (cooltools v0.7.1)[5], which computes the first principal component (E1) of the observed/expected Hi-C contact matrix. The resulting eigenvector values were assigned to genomic bins, and genome-wide A/B compartment profiles were inferred. GC content data for the reference genome (mm10 or hg19) was used to validate the E1 compartment signal (GC coverage files were generated via bioframe v0.7.2).

To quantify the genome-wide strength of compartmentalization, we employed the saddle function in the cooltools v0.7.1 package to compute contact enrichment as a function of eigenvector values. Genomic bins were grouped into 50 quantiles based on E1 values, with extreme 2% (parameters: Q_LO = 0.02, Q_HI = 0.98) discarded to reduce noise (that is, the number of remaining groups is 48). The saddle point matrix was then computed to visualize preferential interactions between genomic regions with similar compartment identities. Importantly, saddle calculations were performed using eigenvector tracks from the all-merged dataset, which represents data from the entire experiment in question. Each .cool file corresponding to a given replicate/condition/treatment was compared against the eigenvector track from the all-merged dataset to compute interaction enrichment. This approach ensured robust normalization and allowed for direct comparisons across experimental conditions. All compartment interaction strengths (A–A, B–B, A–B, B–A) were extracted only at extent = 10 for downstream analysis; these values were used to calculate the saddle strengths, (AA+BB)/(AB + BA). The selection of extent = 10 corresponds to approximately 20% of the ranked compartment groups in the 48-group scheme here. This selection ensures that interactions across moderately compartmentalized regions are captured while avoiding biases arising from proximity to the diagonal (artefactual effects). Python v3.9.20 was used for calculations.

*Quantification of TAD and Loop strengths*

Hi-C read pairs from each replicate were loaded at 5 kb resolution with "cooler cload" pairs against the appropriate genome assembly (hg19 or mm10) and filtered to canonical autosomes, removing mitochondrial and sex chromosomes. Multi-resolution .mcool files were generated with "cooler zoomify" at 10, 25, 50, and 100 kb, and each resolution was balanced by iterative correction (cooler balance). TADs and chromatin loops were called on merged contact maps of vehicle-treatment replicates (one per condition/cell line), generated by upstream merging and pre-processing described prior, which served as reference feature sets used for feature quantification in individual replicates.

For TAD calling, merged .mcool files were converted to HDF5 with hicConvertFormat and TADs were identified at 50 kb using hicFindTADs (--minDepth = 3 × resolution, --maxDepth = 5 × resolution, --step = resolution, --thresholdComparisons 0.01, --delta 0.01, Benjamini–Hochberg FDR correction $q \leq 0.01$). Insulation scores were then computed on each replicate at 50kb using cooltools.insulation (300 kb diamond window, ignore_diags = 2). The resulting insulation table was filtered to exclude bins flagged as bad and those with fewer than 50% valid pixels within the diamond. Reference boundaries were then joined to the filtered replicate table, and those whose corresponding bin was missing were excluded. The $\log_2$(mean) and $\log_2$(median) of the remaining per-boundary insulation scores were reported as per-replicate boundary-strength summaries. Pileup maps were obtained by piling up the replicate's balanced matrix at the merged-reference boundaries with coolpup.pileup in local mode (flank = 30 × resolution, rescale = False), normalized by the replicate's cis-expected (cooltools.expected_cis).

Chromatin loops were called at 25 kb with cooltools.dots, using a cis-expected background (cooltools.expected_cis) computed over chromosome arms in human and whole chromosomes in mouse, with anchor separation ≤ 10 Mb. Loop strength was quantified at 25 kb by piling up the replicate's matrix at the reference loop anchors (no rescaling) and computing a peak-to-lower-left ratio (P2LL) on the resulting $\log_2$(O/E) mean map. The peak window is a centered quarter-width square, rounded up to the nearest odd integer; the lower-left (LL) background is an equivalently-sized square in the lower-left corner of the map. P2LL is the mean of the peak pixels minus the mean of the LL pixels on this $\log_2$ map. All analyses performed in Python 3.9.0 using cooler v0.9.3, cooltools v0.7.1, bioframe v0.8.0, coolpuppy v1.1.0, and HiCExplorer v3.7.6.

*Differential contact maps*

Differential contact maps were created from .hic files after dividing the Juicer-generated contact matrices with the total number of contacts in each sample (to generated contacts per million [CPM] values), and subtracting the 2 normalized matrices of interest.

*E1, insulation, and differential track generation*
Hi-C contact matrices were converted to .cool format and balanced using iterative correction. All analyses were performed at 25 kb resolution, unless otherwise mentioned. Compartment profiles (E1) were computed using "cooltools eigs-cis", with GC content used as a phasing track to ensure consistent orientation of eigenvectors across samples. Insulation scores were calculated using "cooltools insulation" with a 200 kb sliding window, excluding the first two diagonals to minimize short-range contact bias. These steps yielded genome-wide tracks of compartment identity (E1) and local chromatin insulation strength at each genomic bin. To assess perturbation-induced changes, differential tracks were generated by subtracting the matched DMSO control from treatment conditions (e.g., TSA − DMSO) at each genomic bin, producing ΔE1 and Δinsulation profiles. Bins flagged as low-quality were excluded prior to computing insulation differences. The resulting delta tracks capture locus-specific shifts in compartmentalization and boundary strength upon treatment. All tracks, including Hi-C contact maps, E1, insulation, and their corresponding differential profiles, were visualized using pyGenomeTracks.

*Dip-C data analysis*
Dip-C data were analyzed as previously described[3] with 2 minor modifications: (1) differential scA/B analysis was performed with 2-sided t-tests, rather than U-tests, with FDR < 1% and |ΔscA/B| > 0.02 (same as Plate-C); (2) the scA/B matrix was visualized with PCA-initialized UMAP, rather than t-SNE.

*Bulk RNA data analysis*
Bulk transcriptome data were pre-processed with STAR and the mouse reference mm39 and analyzed following the DESeq2 (v1.44.0) manual (https://bioconductor.org/packages/release/bioc/vignettes/DESeq2/inst/doc/DESeq2.html) with "design = ~ batch + condition". Protein coding genes were filtered with "smallestGroupSize <- 3" and "keep <- rowSums(counts(dds) >= 10) >= smallestGroupSize." For PCA and k-means clustering, DESeq2 normalized expression values were generated with "rlog" (parameters: "blind=FALSE").

*Single-cell RNA data analysis*
Single-cell transcriptome data were pre-processed with 10x Genomics Cell Ranger (v8.0.1) using the mouse reference "refdata-gex-GRCm39-2024-A." For cell type annotation, the 4 samples were integrated following the Seurat (v5.2.1) tutorial "Introduction to scRNA-seq integration" (https://satijalab.org/seurat/articles/integration_introduction.html), and clustered with "FindClusters" (parameters: "resolution = 1"). Clusters were manually merged and annotated based on public databases. Differentially expressed genes were identified from all protein coding genes following the Seurat tutorial "Differential expression testing" (https://satijalab.org/seurat/articles/de_vignette.html) with 2 minor modifications to ensure stringency: (1) threshold for padj were set to 0.01 (FDR < 1%) for both single-cell and pseudo-bulk tests, rather than thresholding on p; (2) a threshold for |log2FC| (> 0.5) was added for both singlecell and pseudo-bulk results

**a** Cost and time-scale for Plate-C

| Step | Key Cost Items | Cost | Time |
|---|---|---|---|
| Fixation & Permeabilization | | Negligible | < 1 h |
| Restriction Digestion | Restriction Enzymes | $1.6 / Reaction | 1 h |
| Proximity Ligation | DNA Ligase | $1.5 / Reaction | < 1 h |
| Lysis | | Negligible | 16 h |
| Library Preparation | Tn5 Transposome | $0.7 / Reaction | 5 h |
| DNA Sequencing | Sequencing Service (NovaSeq X Plus) | $2.8 / Gb | 48 h |
| | **Total:** | **$3.7 / Reaction (Plate-C) + $2.8 / Gb (Sequencing)** | **< 24 h (Plate-C) + 48 h (Sequencing)** |

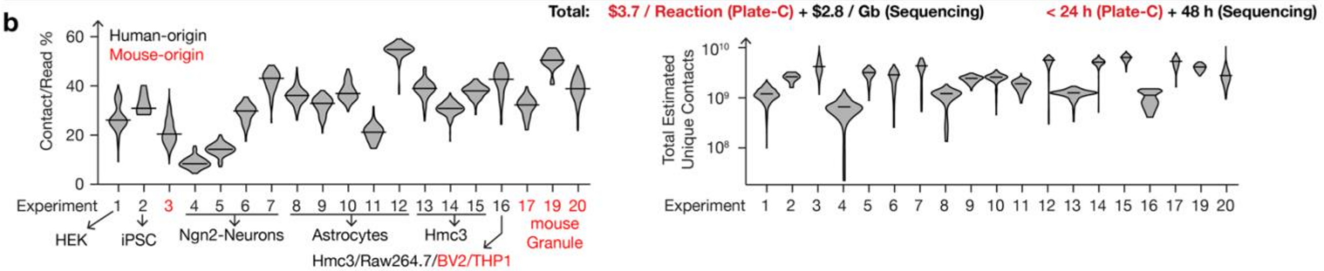


**c** Validating Plate-C in HEK cells

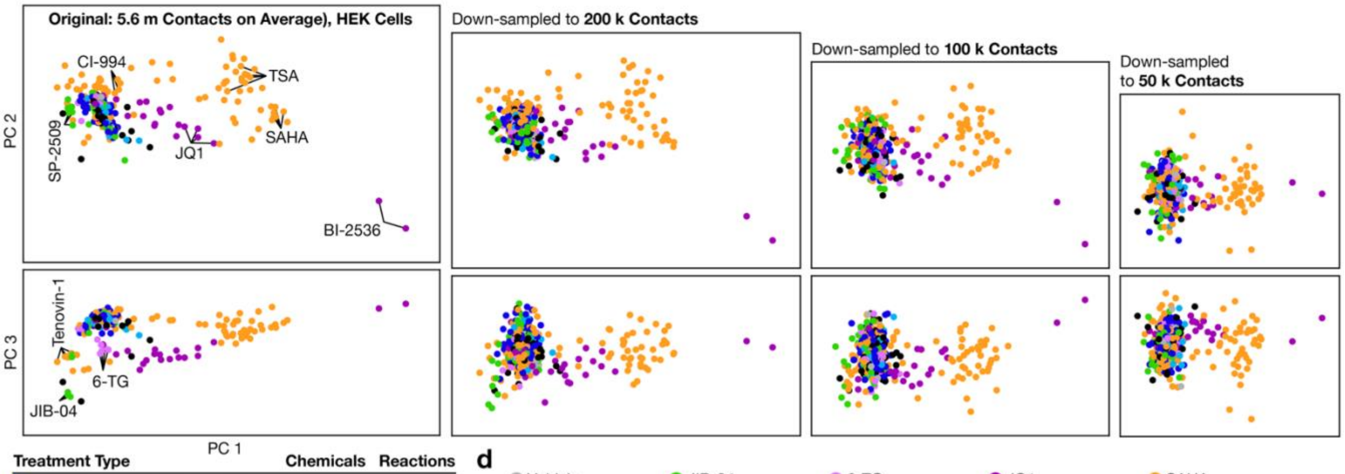


| Treatment Type | Chemicals | Reactions |
|---|---|---|
| Histone Deacetylase Inhibitor (HDACi) | 60 | 134 |
| Histone Demethylase Inhibitor (HDMi) | 17 | 42 |
| Histone Acetyltransferase Inhibitor (HATi) | 11 | 24 |
| Histone Methyltransferase Inhibitor (HMTi) | 31 | 75 |
| DNA Methyltransferase Inhibitor (DNMTi) | 8 | 22 |
| BET Domain Inhibitor (BETi) | 13 | 28 |
| Other | 17 | 43 |
| Vehicle | 1 | 12 |

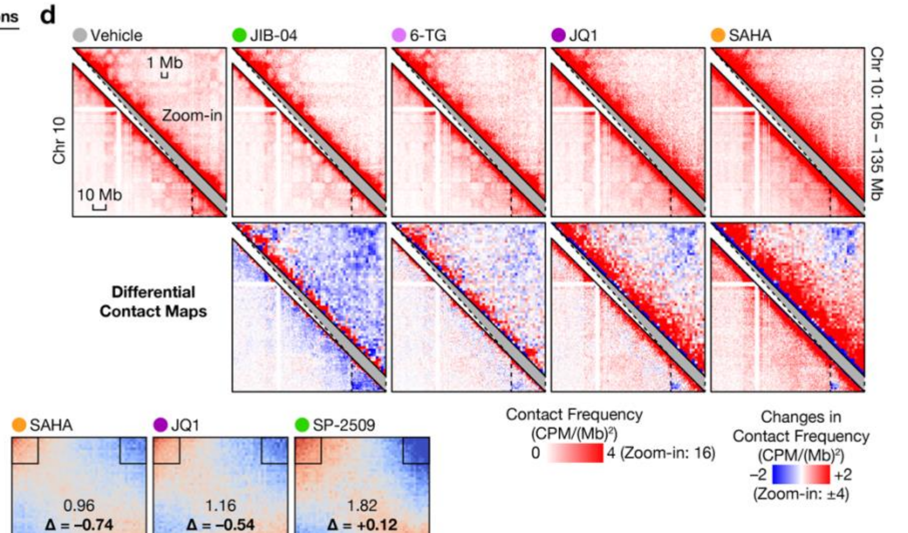


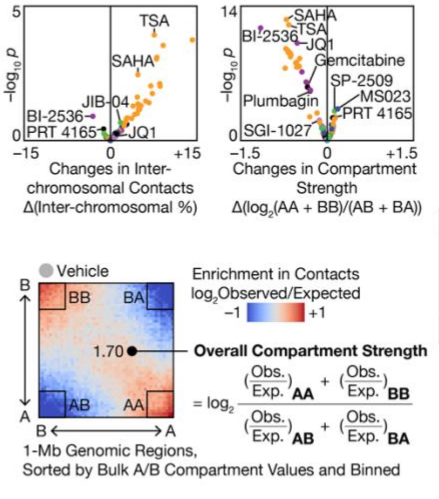


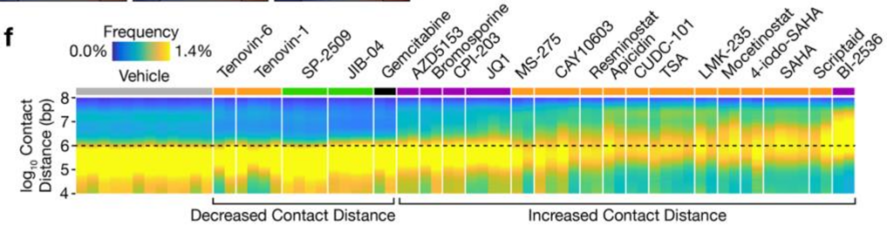

**Extended Data Fig. 1 | Performance, scalability, and robustness of Plate-C**

**a**, Estimated cost and time associated with each step of the Plate-C workflow. Total per-reaction cost and processing time are indicated. **b**, (Left) Distribution of contact efficiency (fraction of reads with valid contacts) and (Right) estimated total number of unique contacts across experiments spanning multiple cell types and species. **c**, PCA of scA/B profiles in HEK cells at full sequencing depth (mean ~5.6 million contacts per sample) and following down-sampling to 200k, 100k, and 50k contacts, demonstrating stability of clustering and separation of perturbations, colored by treatment type. **d**, Aggregated (top) and differential (bottom) chromatin 3D contact maps quantitatively visualized local and global patterns of chemically induced genome restructuring. Bin sizes were 1 Mb for whole chromosome maps, 500 kb for differential zoom-in maps for selected perturbations. **e**, Volcano plots revealed chemically induced changes in overall 3D genome characteristics such as extents of inter-chromosomal contacts (left) and strengths of chromatin A/B compartmentalization (right). *p*-values were from 2-sided t-tests. (Bottom) Quantification of the overall strengths of chromatin A/B compartmentalization with saddle plots generated by cooltools. Colors denoted contact frequencies normalized by expected frequencies based on contact distances (red: high, blue: low). Small boxes denoted parts of the saddle matrix that were used to calculate contact strengths of the A compartment (A–A contacts, or "AA"), of the B compartment ("BB"), and between the 2 compartments ("AB" and "BA"). Δ values denoted changes with respect to vehicle controls. **f**, Histograms of intra-chromosomal contact distances for each Plate-C reaction, showing chemically induced shifts in contact distances.

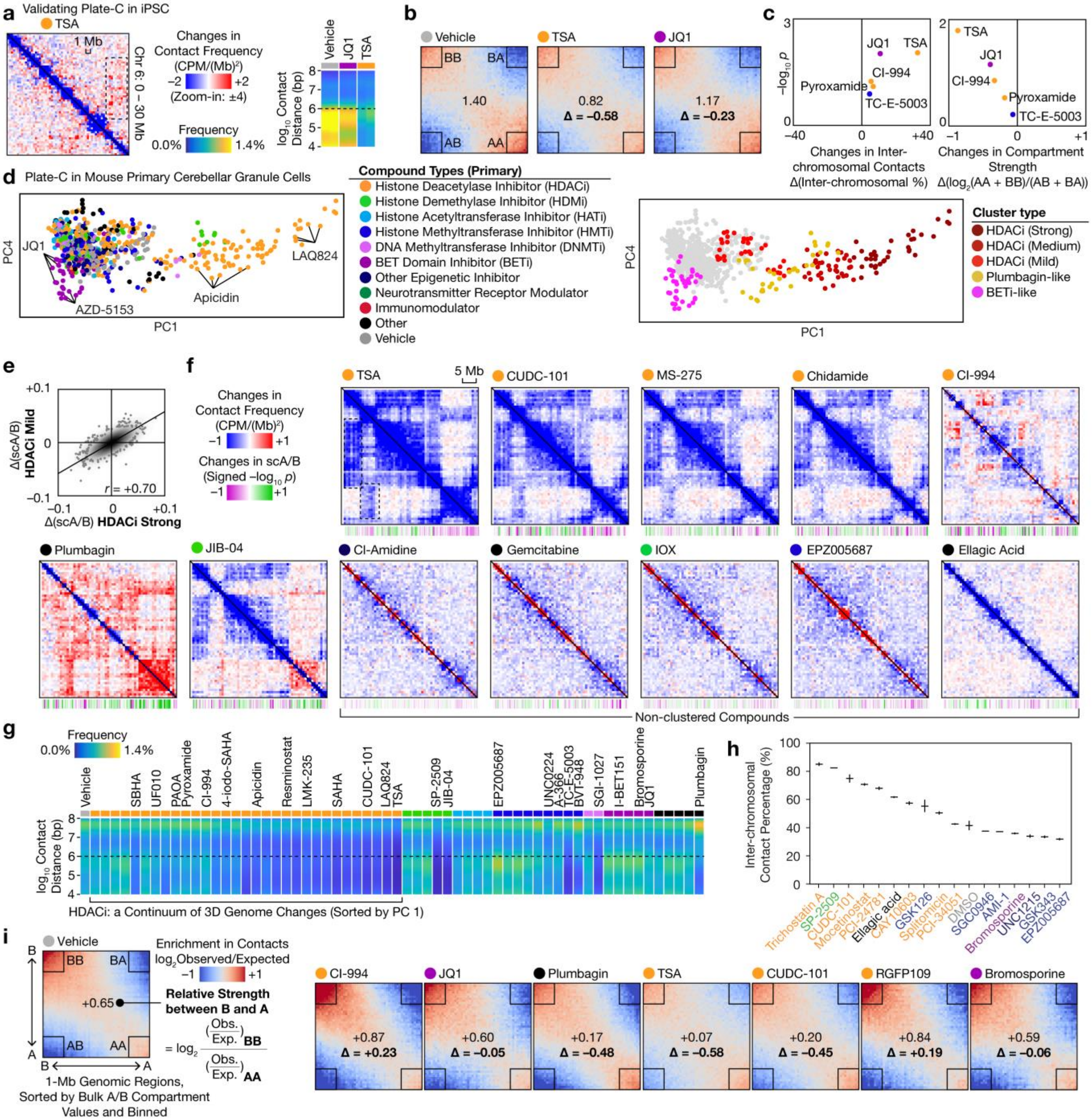

a
Validating Plate-C in iPSC
TSA
1 Mb
Chr 6: 0 – 30 Mb
Changes in Contact Frequency (CPM/(Mb)²)
−2 +2
(Zoom-in: ±4)
Frequency
0.0% 1.4%
Vehicle
JQ1
TSA
log10 Contact Distance (bp)
b
Vehicle
TSA
JQ1
BB
BA
AB
AA
1.40
0.82
Δ = −0.58
1.17
Δ = −0.23
c
−log10 p
JQ1
TSA
CI-994
Pyroxamide
TC-E-5003
Changes in Inter-chromosomal Contacts Δ(Inter-chromosomal %)
Changes in Compartment Strength Δ(log2(AA + BB)/(AB + BA))
d
Plate-C in Mouse Primary Cerebellar Granule Cells
PC4
PC1
JQ1
AZD-5153
Apicidin
LAQ824
Compound Types (Primary)
Histone Deacetylase Inhibitor (HDACi)
Histone Demethylase Inhibitor (HDMi)
Histone Acetyltransferase Inhibitor (HATi)
Histone Methyltransferase Inhibitor (HMTi)
DNA Methyltransferase Inhibitor (DNMTi)
BET Domain Inhibitor (BETi)
Other Epigenetic Inhibitor
Neurotransmitter Receptor Modulator
Immunomodulator
Other
Vehicle
Cluster type
HDACi (Strong)
HDACi (Medium)
HDACi (Mild)
Plumbagin-like
BETi-like
e
Δ(scA/B) HDACi Mild
Δ(scA/B) HDACi Strong
r = +0.70
f
Changes in Contact Frequency (CPM/(Mb)²)
−1 +1
Changes in scA/B (Signed −log10 p)
−1 +1
5 Mb
TSA
CUDC-101
MS-275
Chidamide
CI-994
Plumbagin
JIB-04
Cl-Amidine
Gemcitabine
IOX
EPZ005687
Ellagic Acid
Non-clustered Compounds
g
Frequency
0.0% 1.4%
Vehicle
SBHA
UF010
PAOA
Pyroxamide
CI-994
4-iodo-SAHA
Apicidin
Resminostat
LMK-235
SAHA
CUDC-101
LAQ824
TSA
SP-2509
JIB-04
EPZ005687
UNC0224
A-366
TC-E-5003
BVT-948
SGI-1027
I-BET151
Bromosporine
JQ1
Plumbagin
log10 Contact Distance (bp)
HDACi: a Continuum of 3D Genome Changes (Sorted by PC 1)
h
Inter-chromosomal Contact Percentage (%)
Trichostatin A
SP-2509
CUDC-101
Mocetinostat
PCI-24781
Ellagic acid
CAY10603
GSK126
Splitomicin
PCI-34051
DMSO
SGC0946
AMI-1
Bromosporine
UNC1215
GSK343
EPZ005687
i
Vehicle
BB
BA
AB
AA
+0.65
Enrichment in Contacts log2Observed/Expected
−1 +1
Relative Strength between B and A
= log2 (Obs./Exp.)BB / (Obs./Exp.)AA
1-Mb Genomic Regions, Sorted by Bulk A/B Compartment Values and Binned
CI-994
+0.87
Δ = +0.23
JQ1
+0.60
Δ = −0.05
Plumbagin
+0.17
Δ = −0.48
TSA
+0.07
Δ = −0.58
CUDC-101
+0.20
Δ = −0.45
RGFP109
+0.84
Δ = +0.19
Bromosporine
+0.59
Δ = −0.06

**Extended Data Fig. 2 | Additional characterization of genome architectural responses across perturbations**
**a**, Differential chromatin contact maps for 2.5 mM TSA treatment in human iPSC cells, with contact distance distribution (right). Differential maps highlight local and long-range restructuring induced by perturbation in iPSC, bin size: 2.5 Mb. **b**, Saddle plots; values represent $\log_2$(observed/expected) contact frequencies. Insets indicate regions used to quantify compartment strength; Δ values denote changes relative to vehicle. **c**, Volcano plots showing changes in inter-chromosomal contacts (left) and compartment strength (right) across compounds. *p*-values were calculated using two-sided t-tests. **d**, PCA of scA/B profiles colored by compound types (left) and by cluster type (right), illustrating separation of perturbations along principal components. **e**, Correlation of ΔscA/B profiles between HDACi strong and mild clusters, showing concordance of compartment-level changes across perturbation strengths. **f**, Representative differential contact maps for selected clustered (top; HDACi) and non-clustered (bottom) compounds, bin size: 2.5 Mb, with signed *p*-values as tracks. **g**, Heatmap of intra-chromosomal contact distance distributions ($\log_{10}$ genomic distance) across compounds, ordered along a continuum defined by PC1. **h**, Inter-chromosomal contact frequency across perturbations. **i**, Saddle plots for selected compounds, showing changes in relative A/B compartment interaction strengths ($\log_2$ observed/expected) and Δ values relative to vehicle. Signed p-values of 2-sided t-tests for changes in scA/B profile (green: increase, magenta: decrease).

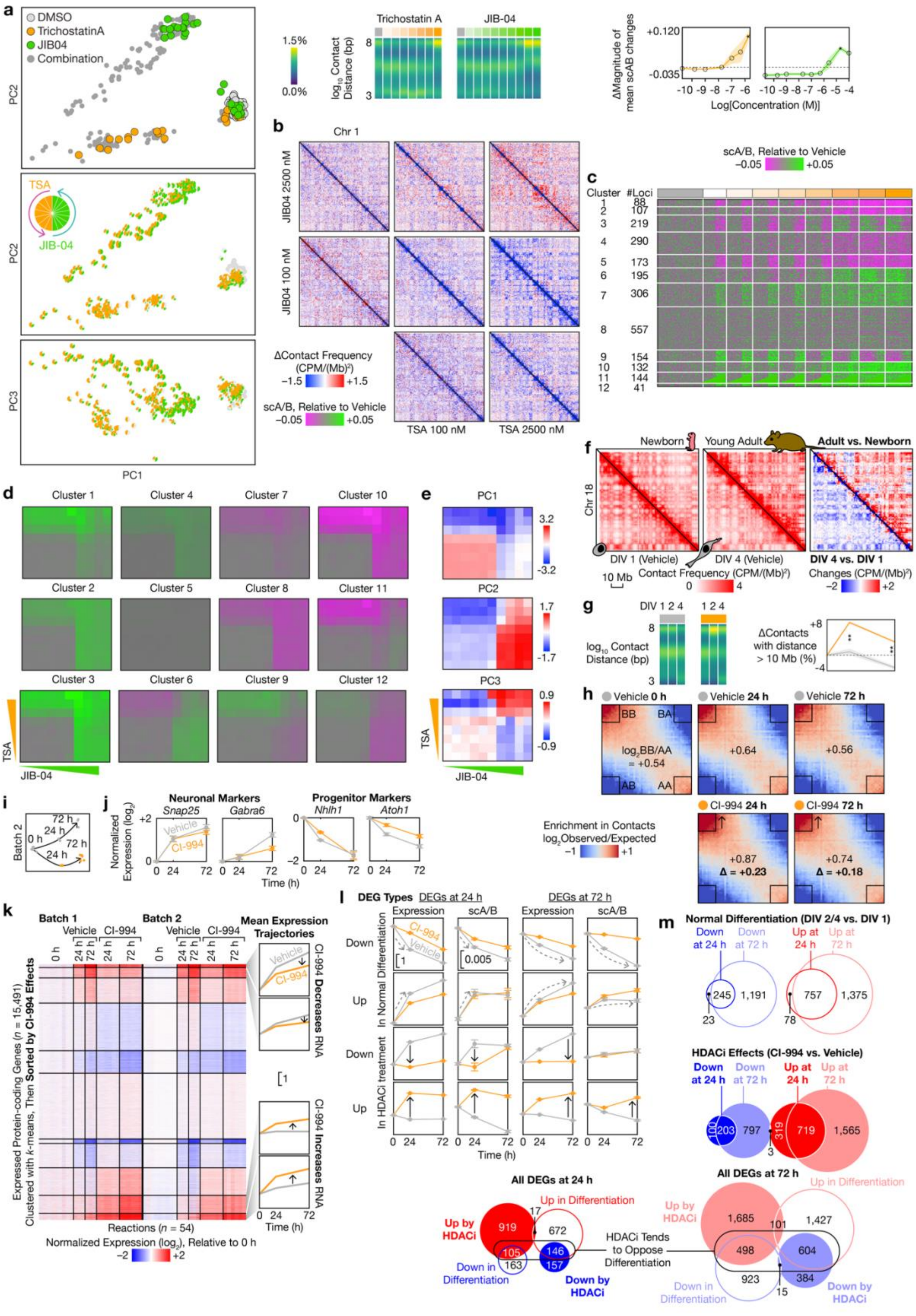

**Extended Data Fig. 3 | Dose-, locus-, and time-dependent effects of perturbations on genome architecture and transcription**

**a**, (Left) Replicate level PCA of the drug combo treatment. (Right) Contact distance distributions and magnitude of scA/B changes found in granule primary screening across concentration gradients for TSA and JIB-04, showing dose-dependent architectural responses. **b**, Differential contact maps across concentrations of JIB-04 and TSA, illustrating non-linear and dose-dependent restructuring patterns. **c**, k-means clustering of all 1-Mb autosomal regions (rows) for different patterns of scA/B changes, k = 12. scA/B values were centered to the mean of DMSO. **d**, Plotting mean scab changes as a heatmap for combinatorial treatments for selected clusters. **e**, Principal component loadings (PC1–PC3) for combinatorial TSA and JIB-04 treatments, capturing orthogonal axes of genome architectural variation. **f**, Aggregated (left) and differential (right) chromatin 3D contact maps visualized the drastic DNA structural differences between neuronal differentiation in vitro (lower left triangles) and in vivo (upper right triangles), bin size: 1 Mb. **g**, Contact distance distributions and quantification of long-range contacts during early neuronal maturation. The CI-994 used here was from the drug library and may be prone to degradation. **h**, Saddle plots for CI-994 treatment, with Δ values relative to vehicle. Arrows (up: increase) denoted specific strengthening of the B compartment by HDACi. **i**, PCA of transcriptomic profiles across time points and treatments, showing temporal trajectories of gene expression. **j**, Transcriptional changes (orange: HDACi, gray: vehicle) of known neuronal (left) and progenitor (right) marker genes confirmed successful differentiation of mouse cerebellar granule cells in vitro. Normalized expression was calculated with DESeq2 as regularized log2 values and centered at 0 h (DIV 1). Error bars denoted SEM. DESeq2 tests for |log2FC| > 0.5. **k**, k-means clustering of all protein-coding genes (rows) revealed different patterns of transcriptional changes, including HDACi-induced changes that were either orthogonal or opposite to changes in normal differentiation. Left: Heatmap of gene expression in each bulk RNA-seq reaction (columns). Right: Average gene expression trajectories of representative clusters. **l**, Average gene expression and scA/B trajectories of various categories of DEGs in vitro showed correlation between transcriptional and architectural changes, especially at 24 h. Top: DEGs in normal differentiation. Bottom: DEGs induced by HDACi. Left: 24 h. Right: 72 h. Error bars denoted SEM. 2-sided t-tests for HDACi induced scA/B changes for DEGs up- and down-regulated by HDACi at 24 h, respectively. **m**, Venn diagrams comparing differentially expressed genes (DEGs; red: up-regulated, blue: down-regulated) between time points (top; bright colors: 24 h, faint colors: 72 h) and between HDACi effects (filled circles) and normal differentiation (hollow circles) in vitro (bottom). Transcriptional changes in differentiation and HDACi induced changes both intensified over time and were negatively correlated to each other (Fisher's exact tests $p = 6 \times 10^{-57}$ and $< 10^{-60}$ at 24 and 72 h). All comparisons are made relative to the vehicle, with $p$ values adjusted using BH-FDR. Significance thresholds: hollow circle, not significant; $*p < 0.05$, $**p < 0.01$.

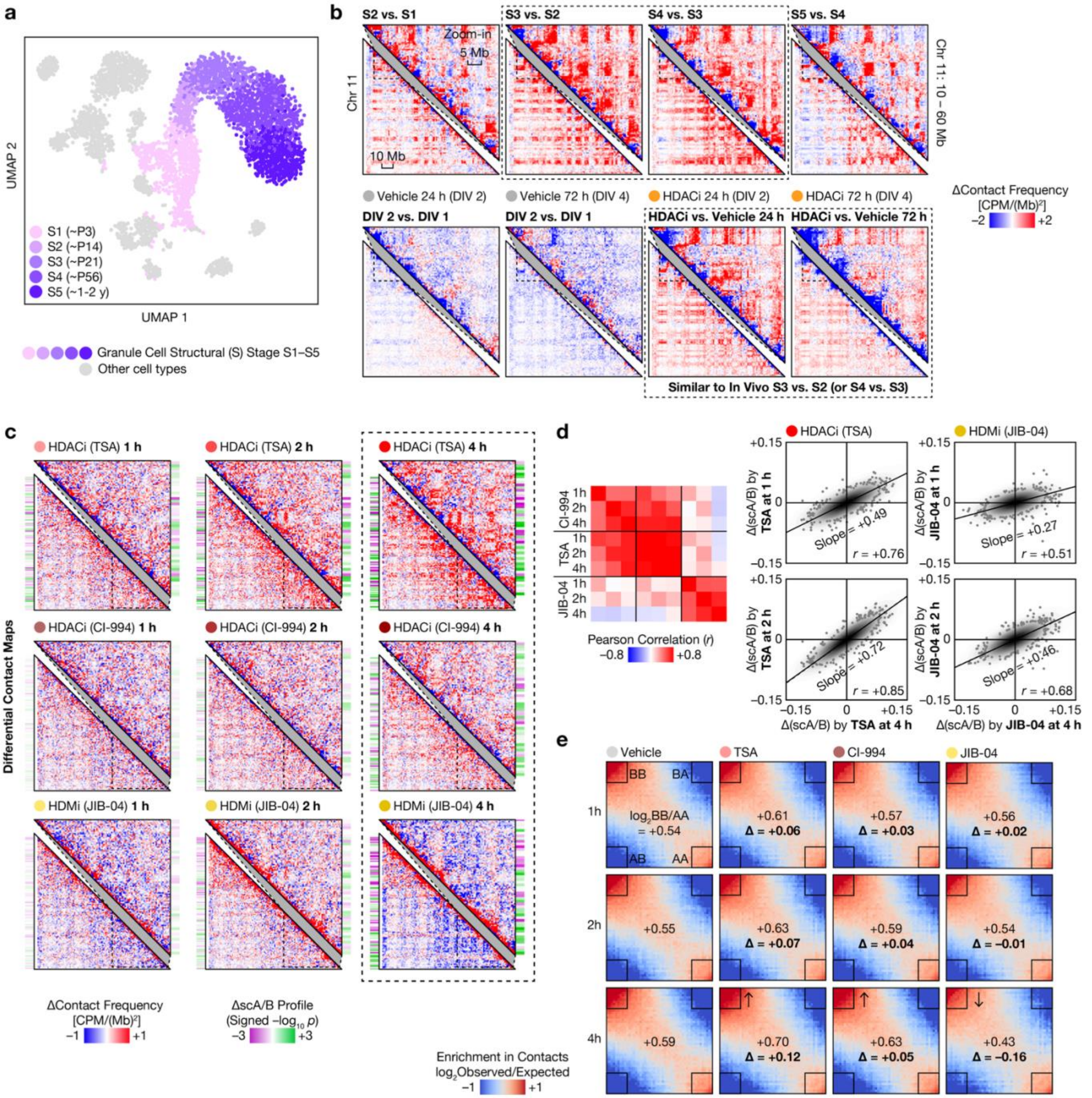

a
UMAP 2
UMAP 1
S1 (~P3)
S2 (~P14)
S3 (~P21)
S4 (~P56)
S5 (~1-2 y)
Granule Cell Structural (S) Stage S1–S5
Other cell types
b
S2 vs. S1
S3 vs. S2
S4 vs. S3
S5 vs. S4
Zoom-in
5 Mb
10 Mb
Chr 11
Chr 11: 10 – 60 Mb
Vehicle 24 h (DIV 2)
Vehicle 72 h (DIV 4)
HDACi 24 h (DIV 2)
HDACi 72 h (DIV 4)
DIV 2 vs. DIV 1
DIV 2 vs. DIV 1
HDACi vs. Vehicle 24 h
HDACi vs. Vehicle 72 h
Similar to In Vivo S3 vs. S2 (or S4 vs. S3)
ΔContact Frequency [CPM/(Mb)²]
−2 +2
c
HDACi (TSA) 1 h
HDACi (TSA) 2 h
HDACi (TSA) 4 h
HDACi (CI-994) 1 h
HDACi (CI-994) 2 h
HDACi (CI-994) 4 h
HDMi (JIB-04) 1 h
HDMi (JIB-04) 2 h
HDMi (JIB-04) 4 h
Differential Contact Maps
ΔContact Frequency [CPM/(Mb)²]
−1 +1
ΔscA/B Profile (Signed −log₁₀ p)
−3 +3
d
CI-994 1h 2h 4h
TSA 1h 2h 4h
JIB-04 1h 2h 4h
Pearson Correlation (r)
−0.8 +0.8
HDACi (TSA)
HDMi (JIB-04)
Δ(scA/B) by TSA at 1 h
Δ(scA/B) by JIB-04 at 1 h
Δ(scA/B) by TSA at 2 h
Δ(scA/B) by JIB-04 at 2 h
Δ(scA/B) by TSA at 4 h
Δ(scA/B) by JIB-04 at 4 h
Slope = +0.49
r = +0.76
Slope = +0.27
r = +0.51
Slope = +0.72
r = +0.85
Slope = +0.46
r = +0.68
+0.15
0
−0.15
−0.15 0 +0.15
e
Vehicle
TSA
CI-994
JIB-04
1h
2h
4h
BB
BA
AB
AA
log₂BB/AA = +0.54
+0.61 Δ = +0.06
+0.57 Δ = +0.03
+0.56 Δ = +0.02
+0.55
+0.63 Δ = +0.07
+0.59 Δ = +0.04
+0.54 Δ = −0.01
+0.59
+0.70 Δ = +0.12
+0.63 Δ = +0.05
+0.43 Δ = −0.16
Enrichment in Contacts log₂Observed/Expected
−1 +1

**Extended Data Fig. 4 | Relationship between perturbation-induced and developmental genome architectural changes**

**a**, UMAP of scA/B profiles only highlighting granule cell structural states (S1–S5) along a developmental trajectory from our previous work. **b**, Differential contact maps comparing adjacent developmental states (S2–S1, S3–S2, S4–S3, S5–S4) and in vitro time points (DIV2 vs. DIV1, DIV4 vs. DIV2), alongside HDACi-induced changes relative to the vehicle. Large, dashed boxes highlighted partial resemblance between juvenile-to-adult development in vivo and HDACi effects in vitro, bin size: 1 Mb. **c**, Differential contact maps following short-term treatment (1–4 h) with HDAC inhibitors (TSA, CI-994) and HDM inhibitor (JIB-04), showing rapid and progressive architectural changes, bin size: 1 Mb. **d**, Pearson correlation (*r*) of ΔscA/B profiles across time points and treatments, with pairwise comparisons highlighting temporal consistency and similarity between perturbations. Scatter plots show correlations between early and later time points for each compound. **e**, Saddle plots showing A–A, B–B, and inter-compartment contacts ($\log_2$ observed/expected) at 1 h, 2 h, and 4 h following treatment. Δ values indicate changes relative to vehicle, Arrows (up: increase) denoted specific strengthening of the B compartment by HDACi.

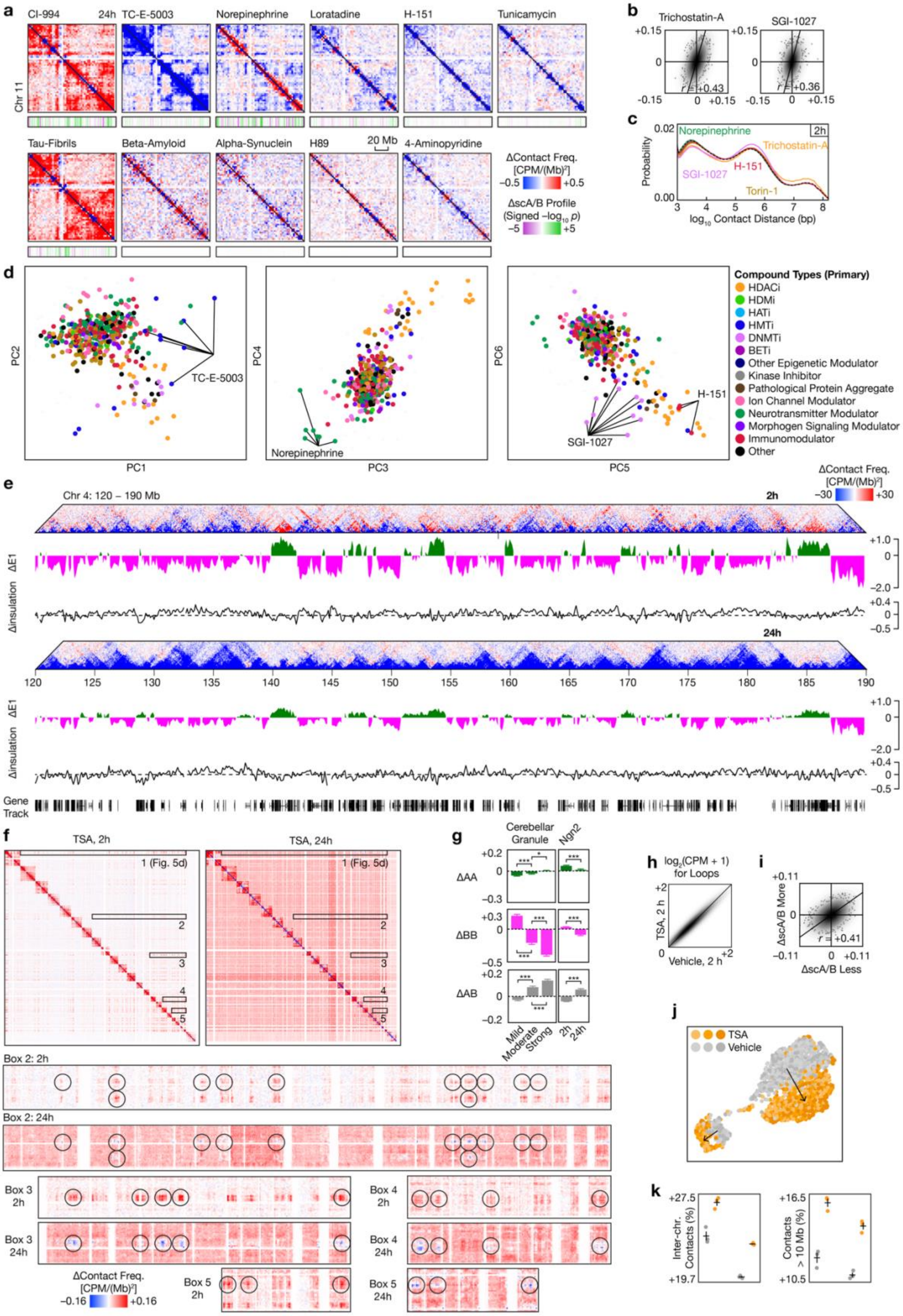
a
CI-994
24h
TC-E-5003
Norepinephrine
Loratadine
H-151
Tunicamycin
Chr 11
Tau-Fibrils
Beta-Amyloid
Alpha-Synuclein
H89
20 Mb
4-Aminopyridine
ΔContact Freq. [CPM/(Mb)²]
−0.5
+0.5
ΔscA/B Profile (Signed −log10 p)
−5
+5
b
Trichostatin-A
SGI-1027
+0.15
0
−0.15
r = +0.43
r = +0.36
c
Norepinephrine
2h
Trichostatin-A
SGI-1027
H-151
Torin-1
Probability
log10 Contact Distance (bp)
d
PC1
PC2
TC-E-5003
PC3
PC4
Norepinephrine
PC5
PC6
H-151
SGI-1027
Compound Types (Primary)
HDACi
HDMi
HATi
HMTi
DNMTi
BETi
Other Epigenetic Modulator
Kinase Inhibitor
Pathological Protein Aggregate
Ion Channel Modulator
Neurotransmitter Modulator
Morphogen Signaling Modulator
Immunomodulator
Other
e
Chr 4: 120 – 190 Mb
2h
24h
ΔContact Freq. [CPM/(Mb)²]
−30
+30
ΔE1
Δinsulation
+1.0
−2.0
+0.4
−0.5
120
125
130
135
140
145
150
155
160
165
170
175
180
185
190
Gene Track
f
TSA, 2h
TSA, 24h
1 (Fig. 5d)
Box 2: 2h
Box 2: 24h
Box 3 2h
Box 3 24h
Box 4 2h
Box 4 24h
Box 5 2h
Box 5 24h
ΔContact Freq. [CPM/(Mb)²]
−0.16
+0.16
g
Cerebellar Granule
Ngn2
ΔAA
ΔBB
ΔAB
Mild
Moderate
Strong
2h
24h
h
log2(CPM + 1) for Loops
TSA, 2 h
Vehicle, 2 h
i
ΔscA/B More
ΔscA/B Less
+0.11
−0.11
r = +0.41
j
TSA
Vehicle
k
Inter-chr. Contacts (%)
+27.5
+19.7
Contacts > 10 Mb (%)
+16.5
+10.5

**Extended Data Fig. 5 | Characterization of perturbation-induced genome architectural changes in neurons**
**a**, Differential contact maps in NGN2 Neurons across diverse perturbations at 24 h, bin size: 2.5 Mb with signed p-values as tracks. **b**, Pearson correlation (r) of ΔscA/B profiles between compounds at 24 h. **c**, Contact distance distributions ($log_{10}$ genomic distance) following 2 h treatments, showing perturbation-specific shifts relative to vehicle. **d**, PCA of scA/B profiles colored by compound types. **e**, Representative locus showing temporal changes (2 h and 24 h) in differential contact frequency, eigenvalue (ΔE1), and insulation following perturbation with 100 nM TSA. **f**, Genome-wide differential contact maps of inter-chromosomal interactions following TSA treatment at 2 h and 24 h. **g**, Quantification of compartment-specific contact changes (ΔAA, ΔBB, ΔAB) in mouse cerebellar granule cells and NGN2 neurons, showing differential modulation of compartment interactions. **h**, Comparison of loop-level contact frequencies between treatment with vehicle and TSA at 2 h. **i**, Correlation of ΔscA/B profiles for TSA treated population between less and more differentiated cells. **j**, UMAP of single-cell scA/B profiles showing replicate consistency of TSA-treated and vehicle-treated Ngn2. **k**, Quantification of inter-chromosomal contacts and long-range contacts (>10 Mb) following TSA treatment. All comparisons are made relative to vehicle, with p values adjusted using BH-FDR. Significance thresholds: *$p < 0.05$, **$p < 0.01$, ***$p < 0.001$. Signed p-values of 2-sided t-tests for changes in scA/B profile (green: increase, magenta: decrease).

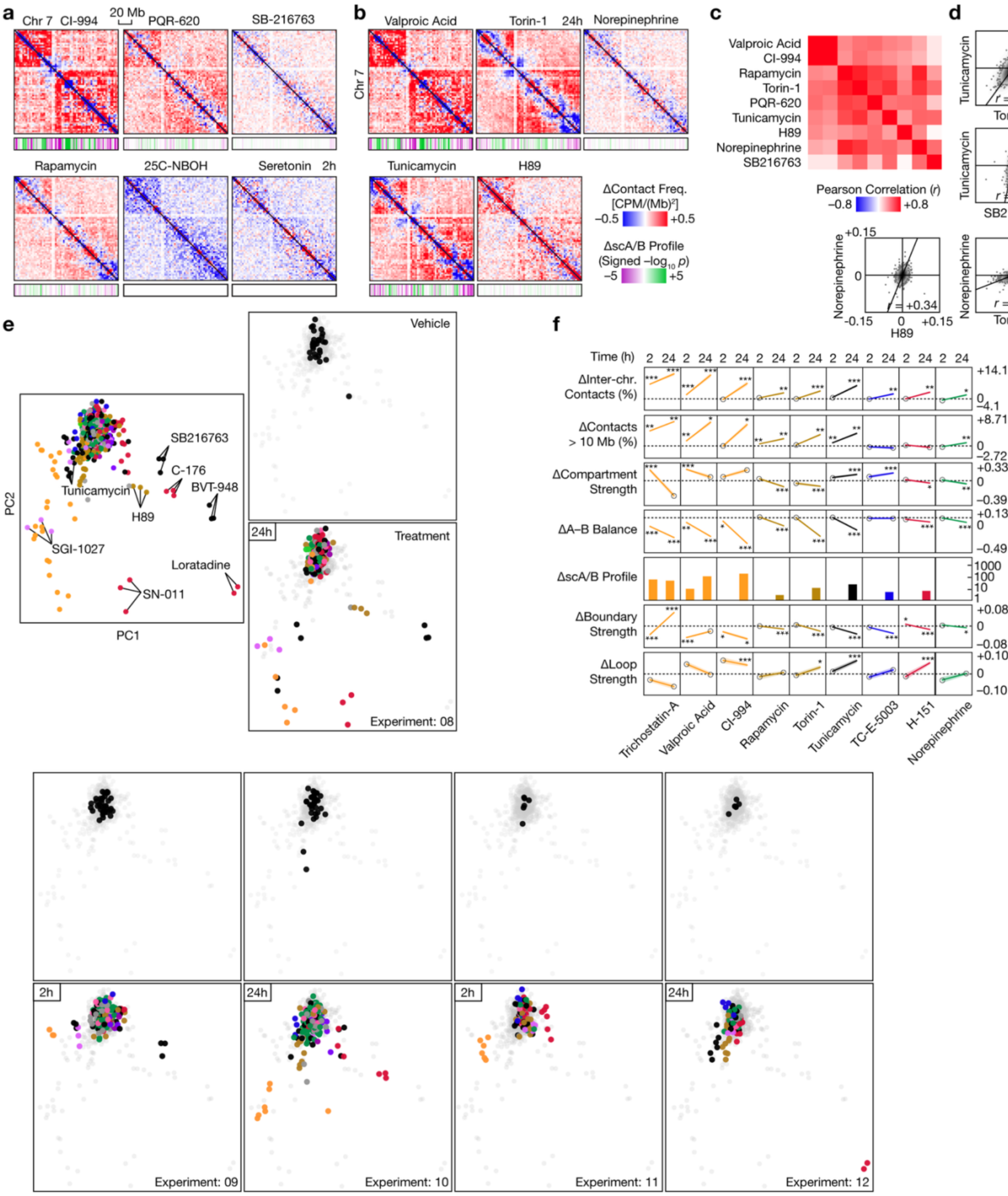
a
Chr 7
CI-994
20 Mb
PQR-620
SB-216763
Rapamycin
25C-NBOH
Seretonin
2h
b
Valproic Acid
Torin-1
24h
Norepinephrine
Chr 7
Tunicamycin
H89
ΔContact Freq. [CPM/(Mb)²]
−0.5
+0.5
ΔscA/B Profile (Signed −log₁₀ p)
−5
+5
c
Valproic Acid
CI-994
Rapamycin
Torin-1
PQR-620
Tunicamycin
H89
Norepinephrine
SB216763
Pearson Correlation (r)
−0.8
+0.8
d
Tunicamycin
Torin-1
r = +0.63
Tunicamycin
SB216763
r = +0.48
Norepinephrine
H89
r = +0.34
+0.15
0
−0.15
0
+0.15
Norepinephrine
Torin-1
r = +0.61
e
PC1
PC2
SB216763
C-176
BVT-948
Tunicamycin
H89
SGI-1027
Loratadine
SN-011
Vehicle
24h
Treatment
Experiment: 08
f
Time (h)
2 24
ΔInter-chr. Contacts (%)
+14.1
0
−4.1
ΔContacts > 10 Mb (%)
+8.71
0
−2.72
ΔCompartment Strength
+0.33
0
−0.39
ΔA–B Balance
+0.13
0
−0.49
ΔscA/B Profile
1000
100
10
1
ΔBoundary Strength
+0.08
0
−0.08
ΔLoop Strength
+0.10
0
−0.10
Trichostatin-A
Valproic Acid
CI-994
Rapamycin
Torin-1
Tunicamycin
TC-E-5003
H-151
Norepinephrine
2h
24h
2h
24h
Experiment: 09
Experiment: 10
Experiment: 11
Experiment: 12

**Extended Data Fig. 6 | Characterization of perturbation-induced genome architectural changes in human primary astrocytes**

**a, b**, Differential contact maps across diverse perturbations for scA/B, bin size: 2.5 Mb with signed *p*-values as tracks, illustrating compound-specific architectural responses at 2h and 24h. **c**, Pearson correlation (*r*) matrix of ΔscA/B profiles across treatments, highlighting relationships between signaling pathways. **d**, Pairwise correlations (*r*) of ΔscA/B profiles between selected perturbations, illustrating shared and distinct responses. **e**, PCA of aggregated and individual scA/B profiles across multiple experiments and time points, demonstrating reproducibility of perturbation-induced changes. For each experiment, top PCA shows the vehicle (black) and treatment (grey), and bottom PCA colored by compound type. **f**, Quantification of changes in genome architectural features across perturbations at 2h and 24h, including (from top to bottom) inter-chromosomal contacts, long-range contacts, compartment strength, A–B balance, scA/B profiles, boundary strength, and loop strength. All comparisons are made relative to vehicle, with P values adjusted using BH-FDR. Significance thresholds: *$p < 0.05$, **$p < 0.01$, ***$p < 0.001$. Signed *p*-values of 2-sided t-tests for changes in scA/B profile (green: increase, magenta: decrease).

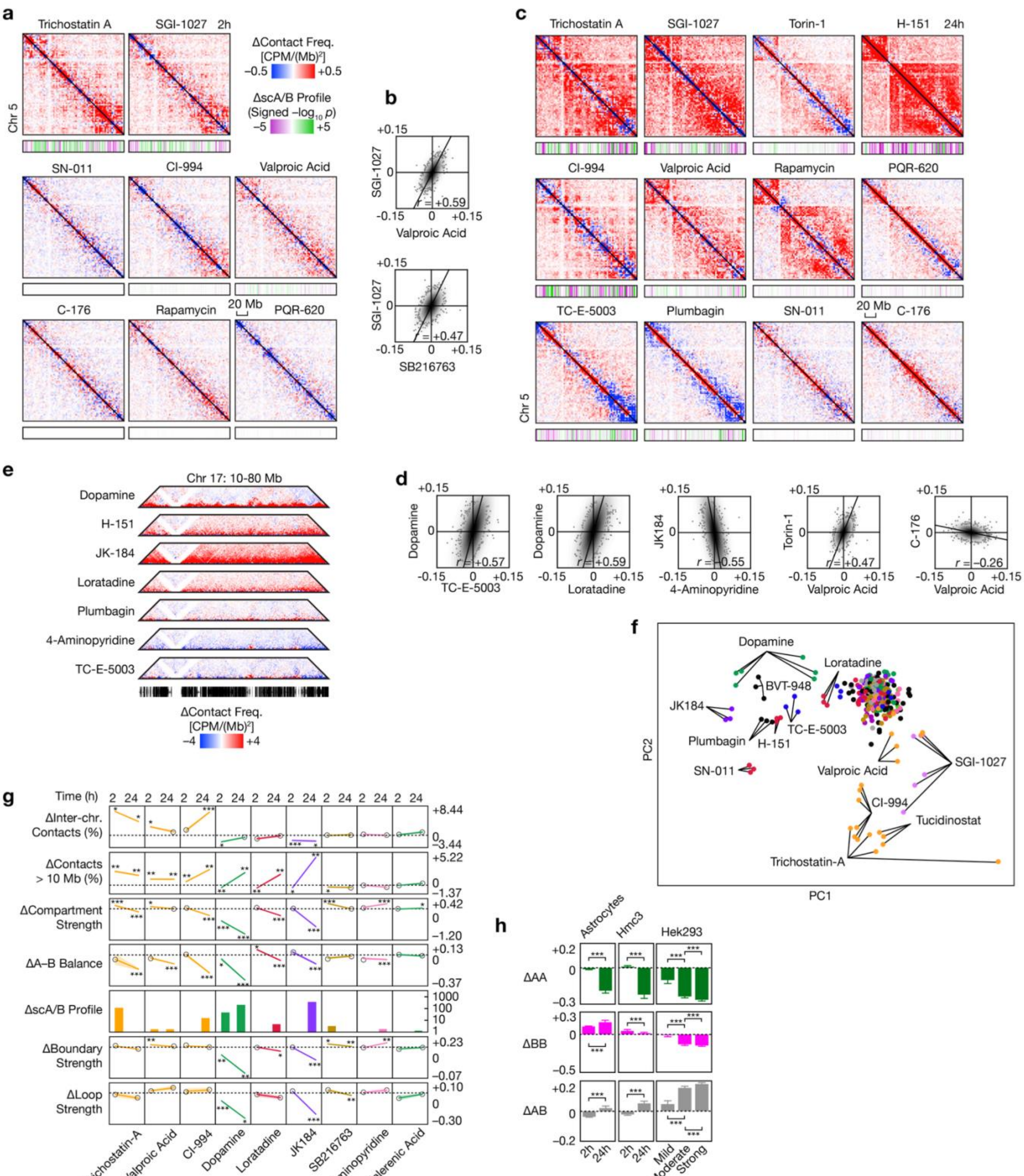

a
Trichostatin A
SGI-1027
2h
ΔContact Freq. [CPM/(Mb)²]
−0.5
+0.5
ΔscA/B Profile (Signed −log₁₀ p)
−5
+5
Chr 5
SN-011
CI-994
Valproic Acid
C-176
Rapamycin
20 Mb
PQR-620
b
+0.15
SGI-1027
0
r = +0.59
−0.15
+0.15
Valproic Acid
SGI-1027
r = +0.47
SB216763
c
Trichostatin A
SGI-1027
Torin-1
H-151
24h
CI-994
Valproic Acid
Rapamycin
PQR-620
TC-E-5003
Plumbagin
SN-011
20 Mb
C-176
Chr 5
e
Chr 17: 10-80 Mb
Dopamine
H-151
JK-184
Loratadine
Plumbagin
4-Aminopyridine
TC-E-5003
ΔContact Freq. [CPM/(Mb)²]
−4
+4
d
Dopamine
r = +0.57
TC-E-5003
Dopamine
r = +0.59
Loratadine
JK184
r = −0.55
4-Aminopyridine
Torin-1
r = +0.47
Valproic Acid
C-176
r = −0.26
Valproic Acid
f
Dopamine
Loratadine
BVT-948
JK184
Plumbagin
H-151
TC-E-5003
SN-011
Valproic Acid
SGI-1027
CI-994
Tucidinostat
Trichostatin-A
PC2
PC1
g
Time (h)
2
24
ΔInter-chr. Contacts (%)
+8.44
0
−3.44
ΔContacts > 10 Mb (%)
+5.22
0
−1.37
ΔCompartment Strength
+0.42
0
−1.20
ΔA–B Balance
+0.13
0
−0.37
ΔscA/B Profile
1000
100
10
1
ΔBoundary Strength
+0.23
0
−0.07
ΔLoop Strength
+0.10
0
−0.30
Trichostatin-A
Valproic Acid
CI-994
Dopamine
Loratadine
JK184
SB216763
4-Aminopyridine
Valerenic Acid
h
Astrocytes
Hmc3
Hek293
ΔAA
+0.2
0
−0.3
ΔBB
+0.3
0
−0.5
ΔAB
+0.2
0
−0.2
2h
24h
2h
24h
Mild
Moderate
Strong

**Extended Data Fig. 7 | Perturbation-induced genome architectural changes in human microglia**
**a,c**, Differential contact maps across diverse perturbations for scA/B in HMC3 cells, bin size: 2.5 Mb with signed *p*-values as tracks, illustrating compound-specific architectural responses at 2h and 24h. **b,d**, Correlation of ΔscA/B profiles between selected perturbations at 2h and 24h **e**, Zoomed-in differential contact maps across selected treatments, highlighting locus-specific patterns of restructuring at 24 h. **f**, PCA of aggregated scA/B profiles across multiple experiments and time points. **g**, Quantification of changes in genome architectural features across perturbations at 2h and 24h, including (from top to bottom) inter-chromosomal contacts, long-range contacts, compartment strength, A–B balance, scA/B profiles, boundary strength, and loop strength. **h,** Quantification of compartment-specific contact changes (ΔAA, ΔBB, ΔAB) in human primary astrocytes, HMC3 and HEK293 cells, showing differential modulation of compartment interactions. All comparisons are made relative to the vehicle, with P values adjusted using BH-FDR. Significance thresholds: $^{*}p < 0.05$, $^{**}p < 0.01$, $^{***}p < 0.001$. Signed *p*-values of 2-sided t-tests for changes in scA/B profile (green: increase, magenta: decrease).

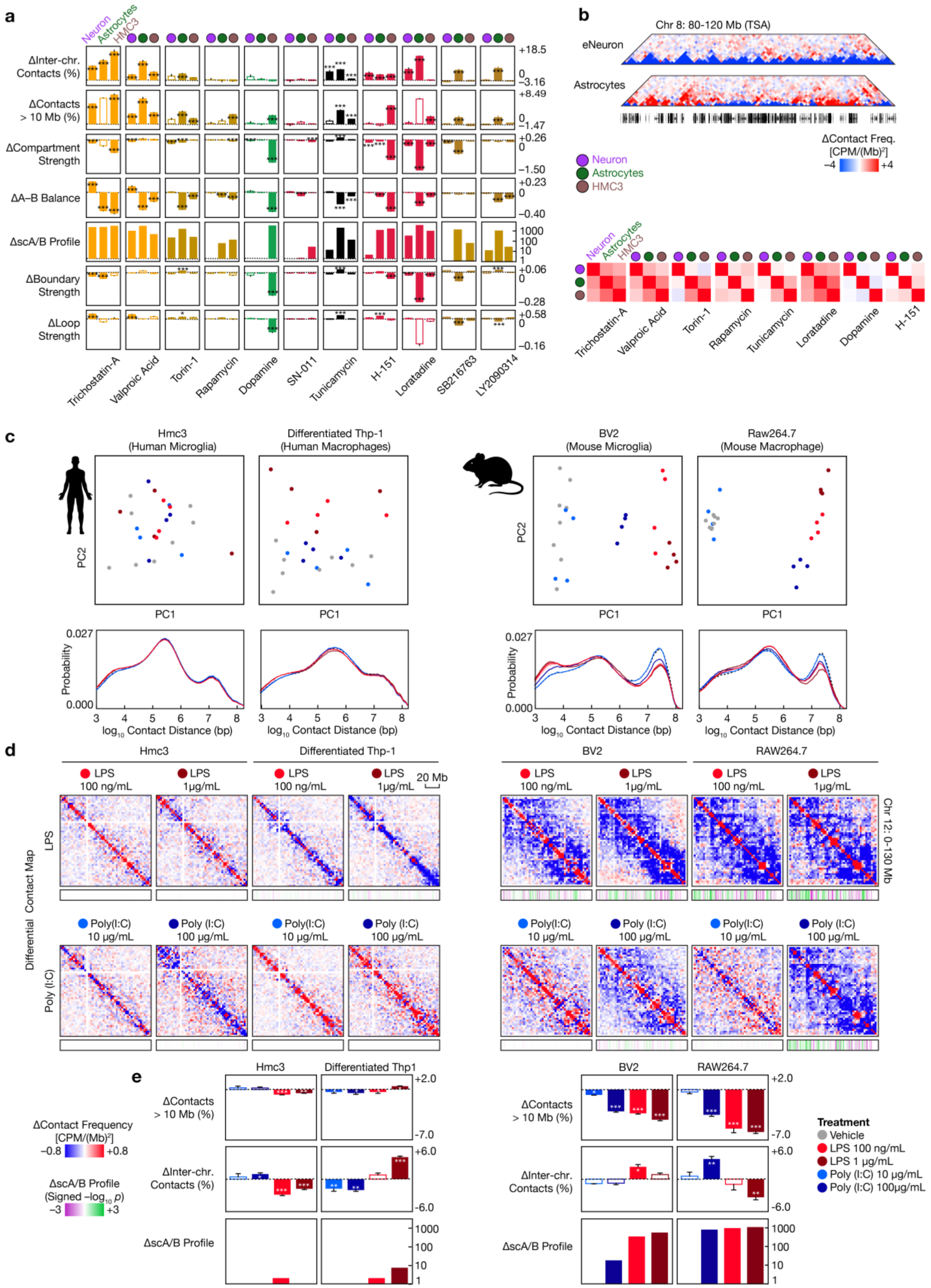

a
Neuron
Astrocytes
HMC3
ΔInter-chr. Contacts (%)
ΔContacts > 10 Mb (%)
ΔCompartment Strength
ΔA–B Balance
ΔscA/B Profile
ΔBoundary Strength
ΔLoop Strength
+18.5
0
–3.16
+8.49
0
–1.47
+0.26
0
–1.50
+0.23
0
–0.40
1000
100
10
1
+0.06
0
–0.28
+0.58
0
–0.16
Trichostatin-A
Valproic Acid
Torin-1
Rapamycin
Dopamine
SN-011
Tunicamycin
H-151
Loratadine
SB216763
LY2090314
b
Chr 8: 80–120 Mb (TSA)
eNeuron
Astrocytes
ΔContact Freq. [CPM/(Mb)²]
–4
+4
Neuron
Astrocytes
HMC3
Trichostatin-A
Valproic Acid
Torin-1
Rapamycin
Tunicamycin
Loratadine
Dopamine
H-151
c
Hmc3 (Human Microglia)
Differentiated Thp-1 (Human Macrophages)
BV2 (Mouse Microglia)
Raw264.7 (Mouse Macrophage)
PC2
PC1
Probability
0.027
0.000
log₁₀ Contact Distance (bp)
d
Hmc3
Differentiated Thp-1
BV2
RAW264.7
LPS 100 ng/mL
LPS 1μg/mL
20 Mb
Chr 12: 0–130 Mb
Differential Contact Map
LPS
Poly (I:C)
Poly(I:C) 10 μg/mL
Poly (I:C) 100 μg/mL
e
Hmc3
Differentiated Thp1
BV2
RAW264.7
ΔContacts > 10 Mb (%)
ΔInter-chr. Contacts (%)
ΔscA/B Profile
+2.0
-7.0
+6.0
-6.0
1000
100
10
1
ΔContact Frequency [CPM/(Mb)²]
–0.8
+0.8
ΔscA/B Profile (Signed –log₁₀ p)
–3
+3
Treatment
Vehicle
LPS 100 ng/mL
LPS 1 μg/mL
Poly (I:C) 10 μg/mL
Poly (I:C) 100μg/mL

**Extended Data Fig. 8 | Cell type– and species-specific genome architectural responses to perturbations**

**a**, Quantification of genome architectural changes across perturbations in multiple attributes, including inter-chromosomal contacts, long-range contacts (>10 Mb), compartment strength, A–B balance, scA/B profiles, boundary strength, and loop strength at 24h. **b**, Representative locus showing differential contact maps in NGN2 Neurons and astrocytes following 100 nM TSA treatment at 24h, bin size: 2.5 Mb with signed p values as track, highlighting cell type–specific responses. **c**, PCA of scA/B profiles (top) and contact distance distributions (bottom) across human (HMC3, THP-1) and mouse (BV2, RAW264.7) immune-derived cell lines following stimulation with 2 doses of LPS and Poly(I:C) treatments. **d**, Differential contact maps for LPS and Poly(I:C) treatments across human and mouse cell lines, bin size: 2.5 Mb with signed *p*-values for scA/B as tracks, illustrating differential genome architectural responses. **e**, Quantification of perturbation-induced changes in long-range contacts, inter-chromosomal contacts, and scA/B profiles across cell lines, highlighting species-specific differences in genome architectural remodeling. All comparisons are made relative to the vehicle, with *p* values adjusted using BH-FDR. Significance thresholds: $*p < 0.05$, $**p < 0.01$, $***p < 0.001$. Signed *p*-values of 2-sided t-tests for changes in scA/B profile (green: increase, magenta: decrease).

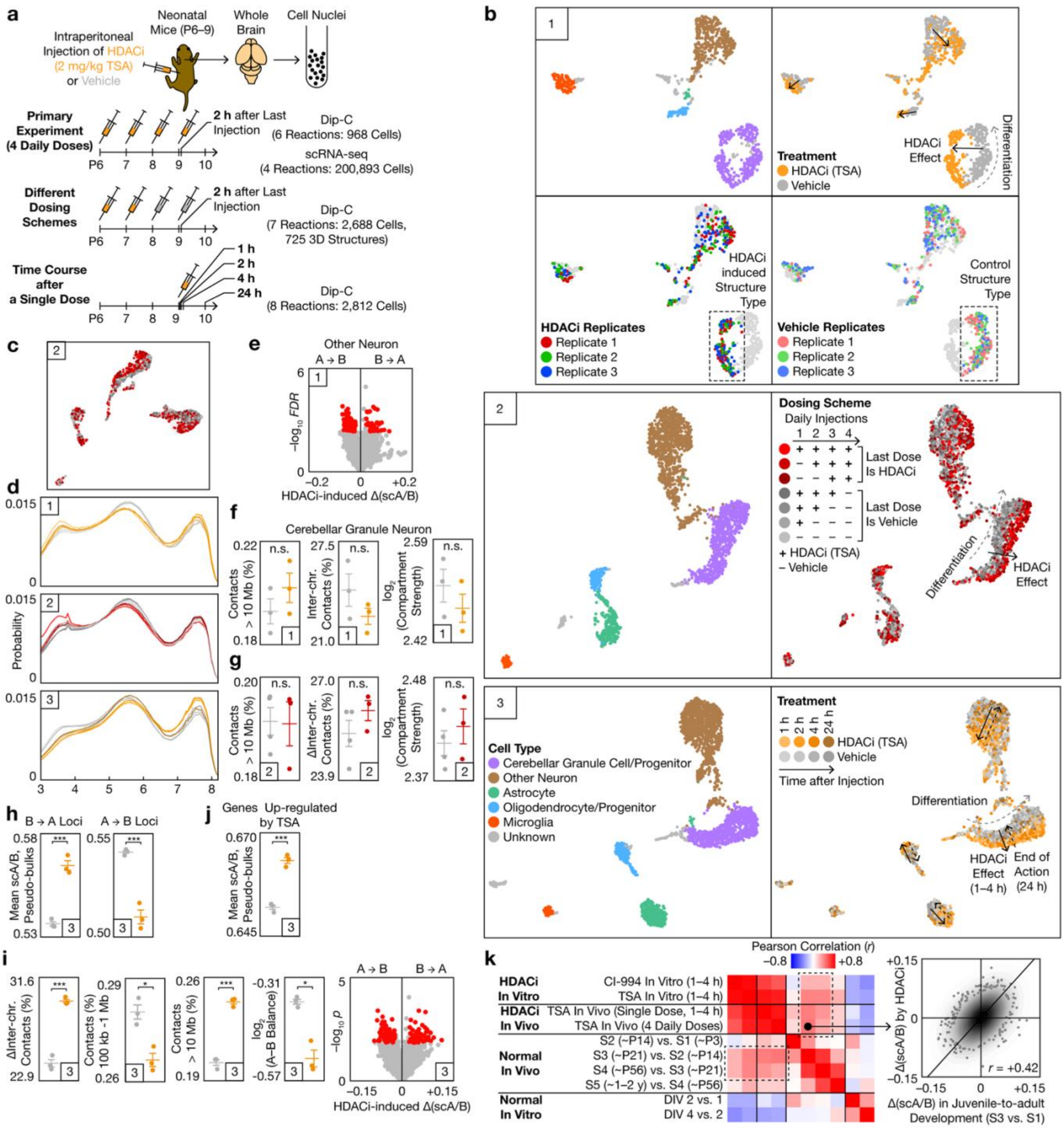
a
Intraperitoneal Injection of HDACi (2 mg/kg TSA) or Vehicle
Neonatal Mice (P6–9)
Whole Brain
Cell Nuclei
Primary Experiment (4 Daily Doses)
2 h after Last Injection
Dip-C (6 Reactions: 968 Cells)
scRNA-seq (4 Reactions: 200,893 Cells)
Different Dosing Schemes
Dip-C (7 Reactions: 2,688 Cells, 725 3D Structures)
Time Course after a Single Dose
1 h
2 h
4 h
24 h
Dip-C (8 Reactions: 2,812 Cells)
b
Treatment
HDACi (TSA)
Vehicle
HDACi Effect
Differentiation
HDACi induced Structure Type
Control Structure Type
HDACi Replicates
Vehicle Replicates
Replicate 1
Replicate 2
Replicate 3
Dosing Scheme
Daily Injections
Last Dose Is HDACi
Last Dose Is Vehicle
+ HDACi (TSA)
– Vehicle
Cell Type
Cerebellar Granule Cell/Progenitor
Other Neuron
Astrocyte
Oligodendrocyte/Progenitor
Microglia
Unknown
Time after Injection
HDACi Effect (1–4 h)
End of Action (24 h)
c
d
Probability
e
Other Neuron
A → B
B → A
HDACi-induced Δ(scA/B)
f
Cerebellar Granule Neuron
Contacts > 10 Mb (%)
Inter-chr. Contacts (%)
log2 (Compartment Strength)
n.s.
g
ΔInter-chr. Contacts (%)
h
B → A Loci
A → B Loci
Mean scA/B, Pseudo-bulks
j
Genes Up-regulated by TSA
i
ΔInter-chr. Contacts (%)
Contacts (%) 100 kb -1 Mb
Contacts > 10 Mb (%)
log2 (A–B Balance)
k
Pearson Correlation (r)
HDACi In Vitro
CI-994 In Vitro (1–4 h)
TSA In Vitro (1–4 h)
HDACi In Vivo
TSA In Vivo (Single Dose, 1–4 h)
TSA In Vivo (4 Daily Doses)
Normal In Vivo
S2 (~P14) vs. S1 (~P3)
S3 (~P21) vs. S2 (~P14)
S4 (~P56) vs. S3 (~P21)
S5 (~1–2 y) vs. S4 (~P56)
Normal In Vitro
DIV 2 vs. 1
DIV 4 vs. 2
Δ(scA/B) by HDACi
r = +0.42
Δ(scA/B) in Juvenile-to-adult Development (S3 vs. S1)

**Extended Data Fig. 9 | In vivo HDAC inhibition reshapes genome architecture across cell types and time**
**a**, Experimental design for in vivo HDAC inhibition using TSA in neonatal mice (P6–P9), including primary dosing, varied dosing schemes, and single-dose time course, followed by Easy Dip-C and scRNA-seq profiling. **b**, Individual UMAP of scA/B profiles colored by cell type, treatment, replicate, dosing scheme, and time after injection, showing separation of HDACi-induced structural states and preservation of underlying differentiation trajectories. **c**, aggregated UMAP highlighting experiment 2 showing individual experiments. **d**, Contact distance distributions across experiments 1 to 3. **e**, Volcano plot of HDACi-induced scA/B changes in neuronal populations, identifying loci transitioning from A to B and B to A loci. **f,g**, Quantification of genome architectural features for experiment 1 and 2, including long-range contacts, inter-chromosomal contacts, and compartment strength, showing modest but consistent changes across replicates. **h**, Changes in compartment switching following HDAC inhibition for experiment 3 at 1, 2 and 4 hours. **i**, Quantification of inter-chromosomal contacts, long-range contacts, and A–B balance experiment 3 merging 1,2, and 4 h time points. **j**, scA/B changes at loci corresponding to genes upregulated by TSA in experiment 3 merging 1,2, and 4 h time points. **k**, Correlation of HDACi-induced structural changes in vivo with in vitro perturbations and normal developmental trajectories, demonstrating conservation of architectural responses across systems. All comparisons are made relative to vehicle; $p$ values denote 2-sided t test. Significance thresholds: *$p < 0.05$, **$p < 0.01$, ***$p < 0.001$.

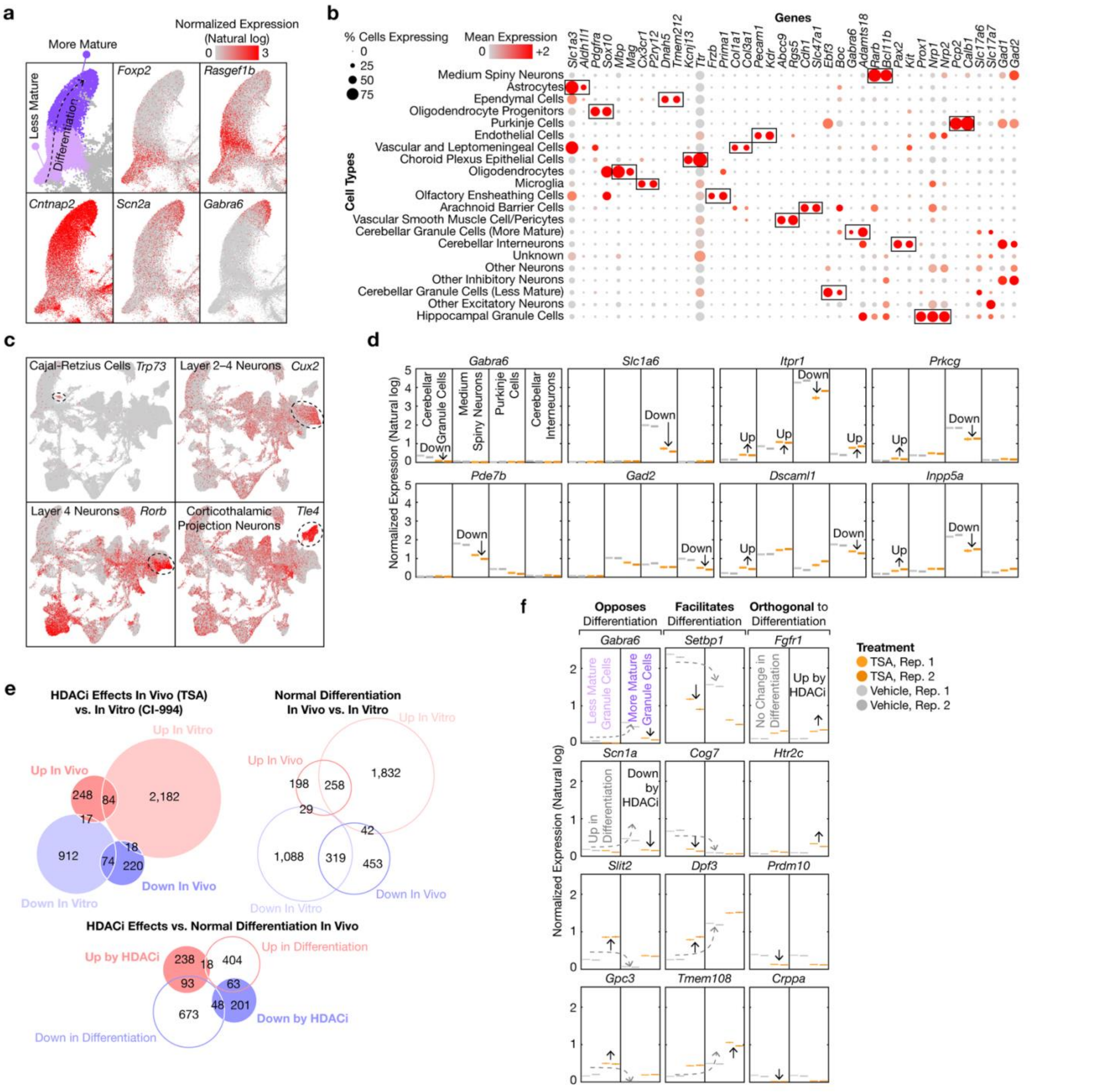
a
Normalized Expression (Natural log)
More Mature
Less Mature
Differentiation
Foxp2
Rasgef1b
Cntnap2
Scn2a
Gabra6
b
Genes
% Cells Expressing
Mean Expression
Cell Types
Medium Spiny Neurons
Astrocytes
Ependymal Cells
Oligodendrocyte Progenitors
Purkinje Cells
Endothelial Cells
Vascular and Leptomeningeal Cells
Choroid Plexus Epithelial Cells
Oligodendrocytes
Microglia
Olfactory Ensheathing Cells
Arachnoid Barrier Cells
Vascular Smooth Muscle Cell/Pericytes
Cerebellar Granule Cells (More Mature)
Cerebellar Interneurons
Unknown
Other Neurons
Other Inhibitory Neurons
Cerebellar Granule Cells (Less Mature)
Other Excitatory Neurons
Hippocampal Granule Cells
c
Cajal-Retzius Cells Trp73
Layer 2–4 Neurons Cux2
Layer 4 Neurons Rorb
Corticothalamic Projection Neurons Tle4
d
Normalized Expression (Natural log)
Cerebellar Granule Cells
Medium Spiny Neurons
Purkinje Cells
Cerebellar Interneurons
Gabra6
Slc1a6
Itpr1
Prkcg
Pde7b
Gad2
Dscaml1
Inpp5a
e
HDACi Effects In Vivo (TSA) vs. In Vitro (CI-994)
Up In Vitro
Up In Vivo
Down In Vitro
Down In Vivo
Normal Differentiation In Vivo vs. In Vitro
HDACi Effects vs. Normal Differentiation In Vivo
Up by HDACi
Up in Differentiation
Down in Differentiation
Down by HDACi
f
Opposes Differentiation
Facilitates Differentiation
Orthogonal to Differentiation
Less Mature Granule Cells
More Mature Granule Cells
No Change in Differentiation
Up by HDACi
Up in Differentiation
Down by HDACi
Normalized Expression (Natural log)
Gabra6
Setbp1
Fgfr1
Scn1a
Cog7
Htr2c
Slit2
Dpf3
Prdm10
Gpc3
Tmem108
Crppa
Treatment
TSA, Rep. 1
TSA, Rep. 2
Vehicle, Rep. 1
Vehicle, Rep. 2

**Extended Data Fig. 10 | Single-cell transcriptomic characterization of HDAC inhibition in vivo**

**a**, Cerebellar granule cells/progenitors formed a continuum of different maturation stages, as shown on the CCA-integrated UMAP (partial). Top left: Louvain clustering of cerebellar granule cells/progenitors into less mature (lighter purple) and more mature (darker purple) cells. Others: Seurat-normalized expression levels of marker genes of various differentiation stages. **b**, Dot plot showing mean expression levels (after normalization and scaling by Seurat; gray: low, red: high) of marker genes across cell types. The radius of each dot was proportional to the percentage of cells that expressed a gene. Boxes highlighted key marker genes of each cell type. **c**, Seurat-normalized expression levels of marker genes of additional neuronal sub-types, shown on the CCA integrated UMAP. **d**, Mean expression levels (after normalization by Seurat) of HDACi-induced DEGs in each reaction (orange: HDACi, gray: vehicle; different shades denoted replicates) across cell types (divided by vertical lines). Arrows (up: increased expression by HDACi, down: decreased expression) denoted statistically significant HDACi induced changes (FDR $< 1\%$ and $|\log2FC| > 0.5$ for both single-cell U-tests and pseudo-bulk DESeq2 tests). Error bars represented SEM. **e**, Venn diagrams of DEGs in cerebellar granule cells/progenitors showed a positive correlation between HDACi-induced DEGs in vivo (4 daily injections of TSA) and in vitro (72 h treatment by CI-994) (Fisher's exact test $p = 10^{-19}$) (top left) and between normal differentiation in vivo (more mature vs. less mature cells on P9) and in vitro (DIV 4 vs. 1) ($p < 10^{-60}$) (top right), and a negative correlation between HDACi effects and normal differentiation in vivo ($p = 2 \times 10^{-10}$) (bottom). **f**, Similar to panel d, but comparing less (left) and more (right) mature cerebellar granule cells/progenitors rather than comparing between cell types. Dashed arrows denoted statistically significant changes in normal differentiation (less vs. more mature cells in control mice). Solid arrows denoted statistically significant changes by HDACi (less and mature cells combined).
