## Supplementary Protocol 1 for "Whole-genome 3D architectural screen reveals modulators of brain DNA structure"

### Tan Lab Plate-C Protocol

Updated Apr 15, 2026

#### Important Notes

NOTE: This version uses half volume for 96-well plates and skips the IGEPAL CA-630 permeabilization.

Use **swinging-bucket centrifuge** to minimize cell loss.

This protocol describes the use of Plate-C for either a full 96- or 384-well plate, accommodating 3k–100k cells per well.

Use a multichannel pipette (P200 for 96-well plates; P20 for 384-well plates) to aspirate reagents from the plate. **<u>Prop the plate at a 45-degree angle and gently position the pipette tips along the well wall closest to you, keeping the tips at the edge of the well bottom to minimize disruption.</u>**

#### Plate Preparation

1. Aim for 25k cells/well in a 96-well plate.

2. Different layouts can be used; ensure that the DMSO is present throughout the plate and not segregated to one row or column to avoid evaporation-related issues.

3. The drugs are prepared in the media in appropriate concentration so that during media change, the final concentration of the drug becomes 1X.

4. Gently shake the plate to mix after drug addition; DO NOT pipette mix in the wells

#### Fixation

1. Freshly prepare **2% PFA in PBS** (recipe below for 12 mL):
    a. 11.25 mL PBS (Invitrogen 10010023)
    b. 750 µL 32% PFA (EMS 15714; if unopened, each glass ampule can be stored indefinitely at RT; once opened, store at 4 C for 4 weeks)
    c. Vortex to mix.
    d. Recipe for 40 mL (4 96-well + overhead) – 37.5 mL PBS + 2.5 mL 32% PFA

2. Completely aspirate media from each well by a gentle flick to get rid of most of the supernatant, followed by a slightly more vigorous flick to remove the rest of the supernatant, leaving cells intact.

3. Fix cells in each well by adding 100 uL **2% PFA in PBS** to each well of a 96-well plate (or 20 uL for 384-well plates), adding to the side wall while tilting the plate.

NOTE: Adding to the side wall is very important as adding directly to the well can blast a hole in the cell layer.

4. Incubate at RT for 10 minutes.

NOTE: If the cells are not suspected to have become less adherent, consider pressing the plate gently against a paper towel. The suction created suffices to remove a considerable amount of the PFA. This may be preferred against flicking multiple times as it is harder to remove 100 μL in this step than the larger volume in step 2.

5. Prepare **10% BSA in PBS** (recipe below for 10 mL):
    a. 5 g BSA (GeminiBio 700-101P)
    b. 30 mL PBS (Invitrogen 10010023)
    c. Heat the solution at 40 C for 1 h to dissolve it, vortex intermittently.
    d. Add PBS to make up the volume to 50 mL.
    e. Vortex to mix. Aliquot and store indefinitely at –20 C.

6. Remove all the supernatants by flicking the plate. Add 100 uL **10% BSA in PBS** to each well of a 96-well plate (or 20 uL for 384-well plates), adding to the side wall while tilting the plate

7. Incubate at RT for 10 minutes.

8. Remove supernatant by flicking the plate.

NOTE: A final round of pipetting may be needed to remove the last drops of BSA (<~40 μL). **Ensure the pipetting is done from the side to avoid creating a void/hole in the cell monolayer.**

PAUSE POINT: Fixed cells can be stored indefinitely at –80 C.

### Chromatin Conformation Capture (3C/Hi-C)

#### Permeabilization

1. Thaw plates on ice.

2. Prepare **0.355% SDS** (recipe below for 12.4 mL):
    a. 440 uL 10% SDS (Sigma 71736)
    b. 11.96 mL water (ThermoFisher 10977023)
    c. Vortex to mix. Store indefinitely at RT.

3. Add 31 uL **0.355% SDS** to each well of a 96-well plate (or 15.5 uL for 384-well plates), adding to the side wall while tilting the plate, and seal with film (the transparent 'B' type: Bio-Rad MSB1001 and roller (Bio-Rad MSR0001).

NOTE: Make sure liquid covers the whole well.

NOTE: The two side flaps (non-sticky) of the film must be removed, to prevent peeling off during incubation/shaking.

4. Incubate at 62 ºC and 800 RPM for 10 min. Gently remove film.

5. Add 11 uL 10% Triton X-100 (Sigma 93443) to each well of a 96-well plate (or 5.5 uL for 384-well plates), adding to the side wall while tilting the plate, and seal with film (the transparent ‘B’ type: Bio-Rad MSB1001 and roller (Bio-Rad MSR0001).

NOTE: The two side flaps (non-sticky) of the film must be removed, to prevent peeling off during incubation/shaking.

6. Incubate at 37 ºC and 800 rpm for 15 min. Gently remove film.

#### Digestion

7. Freshly prepare **Digestion Mix** (recipe below for 3 mL, suitable for 2 96-well plates with 312 uL or 10% overhead):
    a. 964 uL 10X rCutSmart (NEB B9200)
    b. 750 uL water (ThermoFisher 10977023)
    c. 857 uL 5 U/uL MboI (NEB R0147L)
    d. 429 uL 10 U/uL NlaIII (NEB R0125L)
    e. Pipette to mix.

NOTE: NlaIII is stored at –80 ºC.

8. Add 14 uL **Digestion Mix** to each well of a 96-well plate (or 7 uL for 384-well plates), adding to the side wall while tilting the plate, and seal with film (the transparent ‘B’ type: Bio-Rad MSB1001 and roller (Bio-Rad MSR0001).

NOTE: The two side flaps (non-sticky) of the film must be removed, to prevent peeling off during incubation/shaking.

9. Incubate at 37 ºC and 800 RPM for 1 h.

10. Incubate at 65 ºC and 800 RPM for 20 min, Gently remove film.

NOTE: Remove the plate and the heated lid when changing temperature from 37 C to 65 C. Return both after temperature reaches 65 C. The Thermomixer will NOT heat with the heated lid on when the target temperature is set above 62 C.

NOTE: Set the temperature of the plate mixers to 25ºC afterwards in preparation for ligation.

#### Ligation

11. Freshly prepare **Ligation Mix** (recipe below for 4.9 mL, suitable for 1 96-well plates with 484 uL or 10% overhead):
    a. 4.2 mL 2 X Rapid Ligation Buffer (from Vazyme N103-01)
    b. 0.7 mL T4 DNA Ligase (Rapid) (Vazyme N103-01)

    c. Pipette to mix.

NOTE: 2 X Rapid Ligation Buffer is the bigger bottle (NOT the red-capped one), and takes ~1 h to thaw.

12. Mark 1 well of a 96 well-plate (or 4 wells for 384 well-plates) as **Digestion QC**, and 1 other well as **Ligation QC**.

13. Add 46 uL **Ligation Mix** to each well of a 96-well plate (or 23 uL for 384-well plates), EXCEPT the well marked as **Digestion QC** (namely, no ligation), adding to the side wall while tilting the plate, and Seal with film (the transparent 'B' type: Bio-Rad MSB1001 and roller (Bio-Rad MSR0001). Pipette to mix 10 times.

NOTE: Do NOT add **Ligation Mix** to the well marked as **Digestion QC**.

NOTE: Pipette to mix thoroughly because the reagent is viscous.

NOTE: The two side flaps (non-sticky) of the film must be removed, to prevent peeling off during incubation/shaking.

14. Incubate at RT and 800 RPM for 30 min. Gently remove film.

15. Centrifuge at 2500 g for 5 min at 4 C. Remove supernatant by pipetting.

PAUSE POINT: Ligated plates can be stored indefinitely at –80 C.

#### Lysis

16. Prepare **Dip-C Lysis Buffer** (recipe below for 50 mL, sufficient for at least 4 96-well plates):
    a. 46.45 mL water (ThermoFisher 10977023)
    b. 1.25 mL 1 M DTT (Sigma 646563; if unopened, each glass ampule can be stored indefinitely at RT; once opened, store indefinitely at –20 C) (final: 25 mM)
    c. 1 mL 1 M Tris pH 8.0 (ThermoFisher AM9855G) (final: 20 mM)
    d. 750 uL 10% Triton X-100 (Sigma 93443) (final: 0.15%)
    e. 250 uL **100 µM Carrier ssDNA**, TCAGGTTTTCCTGAA (final: 500 nM)
    f. 200 uL 5 M NaCl (ThermoFisher AM9760G) (final: 20 mM)
    g. 100 uL 0.5 M EDTA (ThermoFisher AM9260G) (final: 1 mM)
    h. Vortex to mix. Store indefinitely at –20 C as shared lab stock.

17. Prepare **60 mg/mL Qiagen Protease (QP)**:
    a. 1 vial (7.5 AU) of Qiagen Protease (Qiagen 19157)
    b. 2.78 mL water (ThermoFisher 10977023)
    c. Vortex to mix. Aliquot into ~500 uL per tube and store indefinitely at 4 C as shared lab stock. QP can have batch effect; the powder can be stored for a long time but the dissolved reagent can grow white precipitate at 4ºC when opened many times (no evidence of impacting performance).

NOTE: Always test the whole reaction when a new batch of QP is made, together with an old QP. Switching QP without testing can lead to failure.

18. Freshly prepare **Plate-C Lysis Mix** (recipe below for 11 mL, suitable for 1 96-well plate with 1.4 mL or 15% overhead):
    a. 11 mL **Dip-C Lysis Buffer** (can thaw at 37 C and cool to room temperature)
    b. 110 µL **60 mg/mL Qiagen Protease** (final: 600 µg/mL)
    c. Vortex to mix.

NOTE: This is a key difference between Plate-C and Dip-C, with QP concentration much higher in Plate-C.

19. Add 100 uL **Plate-C Lysis Mix** to each well of a 96-well plate (or 25 uL for 384-well plates), and seal with film
    a. NOTE: the metallic film works better than the transparent 'B' type: Bio-Rad MSB1001 and roller (Bio-Rad MSR0001).

NOTE: The two side flaps (non-sticky) of the film must be removed, to prevent peeling off during incubation/shaking.

20. Incubate at 50 C and 800 RPM for 16 h.

NOTE: The plate has to be perfectly sealed to avoid evaporation at either this step or the following step.

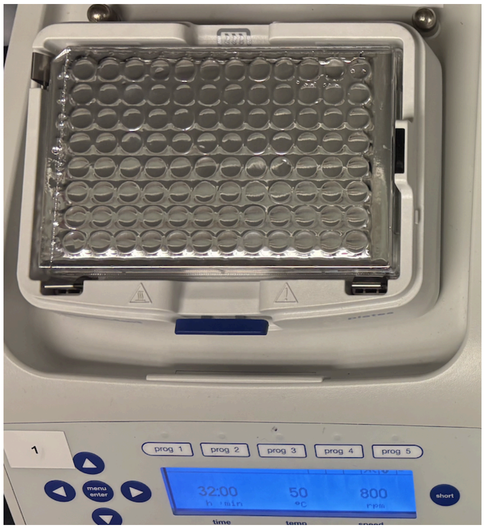


21. Incubate at 70 C and 800 RPM for 20 min.

NOTE: Remove the plate and the heated lid when changing temperature from 50 C to 70 C. Return both after temperature reaches 70 C. The Thermomixer will NOT heat with the heated lid on when the target temperature is set above 62 C.

NOTE: Cells should dissolve during the above steps.

NOTE: A common test for lysis is to check under the phase-contrast microscope for presence of un-lysed nuclei. **<u>Putting the film back on is NOT recommended; use fresh films!</u>** Else, the film could peel and well contents could evaporate.

22. Transfer 2 µL lysate per well to a 96-well (or 384-well) PCR plate, EXCEPT the wells marked as **Digestion QC** and **Ligation QC**. For the 2 wells marked for QC, transfer the entire lysate (100 uL; combine the 4 wells of 25 uL each for 384-well plates) to separate tubes named **Digestion QC** and **Ligation QC**.

PAUSE POINT: Lysate, including the 2 uL in PCR plates, the remaining 98 uL in the original plates, and tubes for QC, can be stored at 4 C overnight, or for a few months at –80 C. Store them in one of the six plate racks at the top of the –80 C. NOTE: The ‘B’ film might peel a little, or become difficult to thaw and peel off, when stored for a long time at –80 C. Feel free to swap it with the ‘F’ film (Bio-Rad MSF1001) before storing at –80 C.

#### Quality Control

23. Purify the DNA from each of the **Digestion QC** and **Ligation QC** with a column (Zymo D4013; DNA Clean & Concentrator-5), using 5 times volume (500 µL) of DNA Binding Buffer, washing 2X with 200 µL wash buffer, and eluting into 10 µL of TE (Invitrogen AM9849).

24. Use 1 µL for quantification using Qubit HS DNA kit and dilute the rest of volume with 1X TE to have a final concentration of (less than) 5 ng/uL, load all of it to e-gel and inspect the trace.

NOTE: Typical concentration is ~20 ng/uL.

NOTE: Prepare samples – load 50 ng of DNA (minimum) + 2 µL 1X E-gel loading buffer; make-up the total volume to 20 µL with water (ThermoFisher 10977023). Use “E-Gel™ 1 Kb Plus DNA Ladder”.

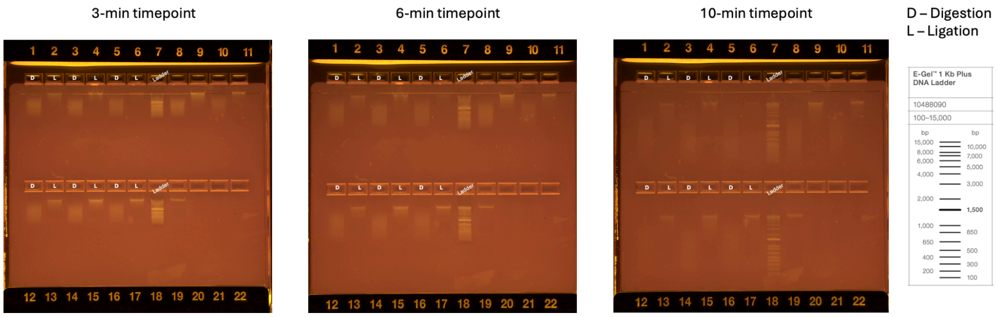


### Whole-genome Amplification

While this section can be easily done by hand, automated liquid handling can greatly accelerate the procedure.

NOTE: The reaction is very similar to Dip-C, but with double the volume, higher QP concentration, and fewer PCR cycles.

#### Determining Optimal Tn5 Concentration

Determine the optimal Tn5 concentration by performing whole-genome amplification (on cells or cell lysate) with varying amounts of Tn5, and identifying the concentration that leads to a flat Bioanalyzer/e-gel profile.

This is also a good positive control (or practice reaction) for new users.

1. Pool 6 µL of lysate each from 3 random control wells into the first tube of a PCR 8-strip
    a. Averaging wells to avoid well-specific issues

2. Transfer 2 µL to each of the remaining 7 tubes of the strip. Remove the excess lysate from the first well ensuring 2 µL in the first tube (just as the remaining 7 tubes).

3. Perform whole-genome amplification with varying amounts of Tn5 transposome (e.g., 50, 25, 12.5, 6.25, 3.13, 1.6, 0.8, 0.4 µL per 850 µL **Transposition Buffer** or following the table below (values slightly different but not posing an issue), *adjusting volumes to 200 µL*. Next, follow the sections Transposition, Stopping, Amplification, and Purification but eluting into 10 µL TE (Invitrogen AM9849) per reaction.
    a. Single-cell Tn5 is ~1.2 µL per 2 mL
    b. Prepare tube 1, then add 100 µL of the Tn5+TB mix to 100 µL TB in tube 2 and so on to do serial dilution
    c. PCR mix (reduced volume as only 8 tubes):
        i. 89.7 µL Q5 reaction buffer
        ii. 89.7 µL Q5 high GC enhancer
        iii. 10.7 µL of dNTP mix
        iv. 1.07 µL of $MgCl_2$
        v. 4.5 µL BSA
        vi. 4.5 µL Q5 DNA polymerase

NOTE: This Tn5 gradient can be set up using serial dilution with a 1:1 or 1:0.5 dilution factor. 1:1 shown here. Follow the 'Instructions' row and set up accordingly.

| | 1 | 2 | 3 | 4 | 5 | 6 | 7 | 8 |
|---|---|---|---|---|---|---|---|---|
| Tn5 (µL/mL) | 50 | 25 | 12.5 | 6.25 | 3.125 | 1.5625 | 0.7812 | 0.3906 |
| 1:X | 1:20 | 1:40 | 1:80 | 1:160 | 1:320 | 1:640 | 1:1280 | 1:2560 |
| *Instructions* | Add 10 uL Tn5 to 190 uL TB | 1:1 dilution in TB | 1:1 dilution in TB | 1:1 dilution in TB | 1:1 dilution in TB | 1:1 dilution in TB | 1:1 dilution in TB | 1:1 dilution in TB |

4. Measure DNA concentration with Qubit.

5. Measure DNA length distribution with Bioanalyzer HS DNA or e-gel (if needed, dilute to less than 5 ng/uL; to send – 5 ng in a total of 3-5µL).

6. Identify optimal Tn5 concentration as the one that leads to the **flattest** Bioanalyzer profile (a little higher on the long side is the most ideal), usually corresponding to a smear centered between 500 bp and 1 kb (the two brightest ladder bands). Perform a finer optimization if needed.

#### Transposition

1. Thaw lysate in a PCR plate. Wipe off condensation from the seal.

NOTE: Unlike Dip-C, if amplification failed, there will be plenty of leftover lysate (98 uL) in the original plate for a redo.

2. Prepare **Transposition Buffer** (recipe below for 50 mL):
    a. 39.06 mL water (ThermoFisher 10977023)
    b. 10 mL 50% PEG 8000 (Hampton Research HR2-535) (final: 10%)
    c. 625 uL 1 M TAPS pH 8.5 (Boston Bio Products BB-2375) (final: 12.5 mM)
    d. 312.5 uL 1 M MgCl2 (ThermoFisher AM9530G) (final: 6.25 mM)
    e. Vortex vigorously (because PEG is viscous) to mix. Aliquot into 2 mL per tube and store indefinitely at –20 C as shared lab stock, labeled “TB”.

3. Freshly prepare **Transposition Mix** (recipe below for 2 mL):
    a. 2 mL **Transposition Buffer** (can thaw at 37 C and cool to room temperature)
    b. Optimized amount of TTE Mix V50 from Vazyme TD501-01
    c. Pipette to mix. NOTE: Do not vortex!

4. Add 8 µL **Transposition Mix** per well:
    a. Seal with film (the transparent ‘B’ type: Bio-Rad MSB1001 and roller (Bio-Rad MSR0001).
    b. Vortex the plate thoroughly.

5. **Vortex the plate thoroughly (at 12 roughly evenly spaced points along the plate) and spin down the plate.**

6. Transpose the genome
    a. 55 C for 10 min
    b. 4C forever.

### Stopping

7. Prepare **Stop Buffer** (recipe below for 50 mL):
    a. 42.45 mL water (ThermoFisher 10977023)
    b. 4.5 mL 0.5 M EDTA (ThermoFisher AM9260G) (final: 45 mM)
    c. 3 mL 5 M NaCl (ThermoFisher AM9760G) (final: 300 mM)
    d. 50 uL 10% Triton X-100 (Sigma 93443) (final: 0.01%)
    e. Vortex to mix. Aliquot into 2 mL per tube (25 tubes) and store indefinitely at –20 C as shared lab stock.

8. Freshly prepare **Plate-C Stop Mix** (recipe below for 1 mL):
    a. 1 mL **Stop Buffer** (can thaw at 37 C and cool to room temperature)
    b. 8 µL **60 mg/mL Qiagen Protease** (final: 480 µg/mL, Qiagen 19157)
    c. Vortex to mix.

NOTE: This is another difference between Plate-C and Dip-C, with QP concentration a bit higher in Plate-C.

9. Add 2 µL **Stop Mix** per well.
    a. **Vortex the plate thoroughly (at 12 roughly evenly spaced points along the plate) and spin down the plate.**

10. Stop transposition:
    a. 50 C for 40 min
    b. 70 C for 20 min
    c. 4 C forever.

PAUSE POINT: Plates can be stored at 4 C overnight. NOTE: Do not store at –20 C, which will greatly reduce the yield!

**NOTE** on Amount of DNA Calculation:
- 96 wells → pool → Take 1/3 of the pooled sample for Zymo → elute: 200 uL, at ~ 20 ng/uL
- 4000 ng / (1/3 of the pooled 96-well plate) → 4000/32 or ~100 ng/well (desired output from library prep)
- Titration: 10 cycles at ~ 4 ng/uL = 40 ng

### Amplification

11. Freshly prepare **PCR Mix** (recipe below for 5 mL) in a regular 15 mL tube:
    a. 2.242 mL Q5 Reaction Buffer (NEB M0491S) (might have white precipitate when thawed for the first time; dissolve by heating with palm and vortexing)
    b. 2.242 mL Q5 High GC Enhancer (NEB M0491S)
    c. 268 μL 10 mM (each) dNTP mix (Vazyme P031-02 or NEB N0447S) (final: 538 μM (each))
    d. 26.8 μL 1 M MgCl2 (ThermoFisher AM9530G) (final: 5.38 mM)
    e. 112 μL 20 mg/mL BSA (NEB B9000S) (final: 448 ug/mL)
    f. 112 μL 2 U/μL Q5 DNA Polymerase (NEB M0491S) (final: 0.0448 U/μL)
    g. Vortex to mix.

12. Add 11 μL **PCR Mix**.

13. Use multi-channel pipette or Bravo to add 2 uL **6.25 uM (each) Nextera Primer Mix** (thaw at room temperature, and centrifuge at 3,000 g for 5 min) per well. NOTE: Set speed (both aspirating and dispensing speeds) of the motorized pipette (Eppendorf) to the slowest (1) to pipette such low volume (1 uL) reliably. Mark the plate with the UDI used.

NOTE: For 96-well plates, we have UDI 1-8. UDI 1-4 is the same as UDI A for single cells (the 384-well UDI) and UDI 5-8 is the same as UDI B.

14. Seal with film (the transparent ‘B’ type: Bio-Rad MSB1001 and roller (Bio-Rad MSR0001). Vortex the plate thoroughly (at 12 roughly evenly spaced points along the plate) and spin down the plate.

15. Run the following 7 cycle (up to 10) PCR:
    a. 4ºC for 3 min
    b. 72ºC for 3 min
    c. 98ºC for 20 s
    d. 7 cycles of 98ºC for 10 s, 62ºC for 1 min, 72ºC for 2 min
    e. 72ºC for 5 min
    f. 4ºC forever.

PAUSE POINT: Plates can be stored at 4ºC overnight or for a few months at –80ºC.

### Purification

16. Pool reactions from the whole plate into 5 times volume of DNA Binding Buffer (Zymo D4004-1-L) (12 mL per 96-well plate, 48 mL per 384-well plate). Attach the plate up-side-down into a V-bottom plate reservoir (ClickBio CBVBLOK200-1) pre-filled with DNA Binding Buffer, centrifuge at 50 g for 5 min, Vortex to mix.

NOTE: If the V-bottom plate reservoir (ClickBio CBVBLOK200-1) does not fit the centrifuge, it will get stuck if put directly; to avoid this, put the plate reservoir on top of a stack of 3 empty 384-well PCR plates.

NOTE: You MUST visually examine the V-bottom reservoir for **cracks** before use, otherwise liquid can be entirely lost. PAUSE POINT: Pooled reaction mixed with Binding Buffer can be stored indefinitely at –20 C.

17. Purify 7.2 mL of mixture by loading 3 columns (each can hold 800 uL) 3 times, and elute into 200 uL TE total across all columns. (Zymo D4013) and elute in TE (Invitrogen AM9849).
    a. 25 µL reaction volume * 96 wells * 6 (as 5X volume of DNA binding buffer added) = 14.4 mL. One-third is 4.8 mL.

NOTE: Please **write the index** (such as “UDI 3”) on every tube to avoid confusion.

18. Measure DNA concentration with Qubit, using the 1x dsDNA HS kit.

NOTE: Typical concentration is ~40 ng/uL. If exceeding 100 ng/uL, the sample is over-amplified (with longer concatemers forming) and PCR cycles should be reduced.

19. Measure DNA length distribution with Bioanalyzer, using a High Sensitivity DNA chip (dilute to less than 5 ng/uL if needed). NOTE: Adjust future Tn5 concentration if length is not optimal.

20. Remove shorter fragments with 0.45X right-side and 0.55X left-side size selection. Do the size selection twice. Record the percentage yield after size selection: ~20% indicates optimal transposition (higher indicates insufficient transposition, lower indicates excessive transposition). We typically size select 100 uL of DNA and elute into 25 uL TE for the final library.

### Size-selection

Plate-C here uses a two-time, double size selection

#### Double-side size-selection (Plate-C)

The protocol here does 0.45X (right side), 0.6X (0.45+0.15=0.60, left side)

1. 100 µL transferred to PCR tube (one per sample)
2. Vortex the SPRI beads thoroughly

NOTE: The beads have to be very well-mixed

3. Add 45 µL of the SPRI beads to each PCR tube; pipette mix well (~10 times)
4. 1 min wait

5. To magnetic rack; wait for 3-5min until solution transparent and beads stuck to the magnet
6. **Transfer supernatant** to fresh PCR tubes (145 µL)
7. Add 15 µL beads to each tube, pipette mix well, incubate for 1 min, transfer to magnetic rack, wait for 3-5min until solution transparent and beads stuck to the magnet
8. Aspirate supernatant, discard it
9. Wash twice with 200 µL 80% Ethanol without disturbing the beads
10. Remove all ethanol completely; use P10 pipette to remove the last drops
11. Dry for 2-3 min until the beads appear slightly shiny (a glossy texture)
12. Add 105 µL TE (22 uL for the second time; use warm TE). Vortex, wait 2min, then spin down, put on magnet until clear
13. Take 100 µL (20 uL for the second time), avoid taking any brown beads
14. Repeat this entire protocol in case of two-time double-side size selection, 20uL elution second time around.
15. NOTE FOR SELF: Estimate input:output DNA amount ratio and keep a record

### Appendix A: E-Gel Quality Control

If many samples need to be checked for quality control, one option is to run agarose gels. We have precast, prestained E-Gel™ Agarose Gels with SYBR™ Safe DNA Gel Stain (ThermoFisher A42100 which has 10 lanes, A42347/double-comb which has 11*2 = 22 lanes; ThermoFisher G820801 with 24*2 = 48 lanes; 96-lane gels and more).

1. Purify DNA as described in *Quality Control*.

2. Thaw 1X E-gel loading buffer (ThermoFisher 10482055) and the ladder of choice. ([https://assets.thermofisher.com/TFS-Assets/BID/Reference-Materials/e-gel-ladder-reference-card.pdf](https://assets.thermofisher.com/TFS-Assets/BID/Reference-Materials/e-gel-ladder-reference-card.pdf))
    a. Use the E-Gel™ **1 Kb Plus DNA Ladder** (size range: 100–15,000 bp) (ThermoFisher 10488090).
    b. Do NOT use the E-Gel™ **1 Kb Plus Express DNA Ladder** (size range: 100–5,000 bp) (ThermoFisher 10488091) as the ligation product exceeds the size range.

3. Prepare samples: load 50 ng of DNA (minimum) + 2 µL 1X E-gel loading buffer; make-up the total volume to 20 µL with water (ThermoFisher 10977023).

4. Undock the camera from the E-Gel™ Electrophoresis Device, place it aside. Open a gel packet and load the gel by clicking it into place in the electrophoresis device.

5. Load samples into the lanes. For each row of samples, ensure the ladder is loaded onto one lane (10 µL to be used; ThermoFisher 10488090).

6. Dock the camera onto the electrophoresis device, set the run (ensure the gel is chosen correctly on the interface – 1% Agarose gel with SYBR™ Safe).

7. The duration should display 26 min; however, capture images at 3-, 6-, 10-minute timepoints (via the user interface on the camera screen). NOTE: do not miss the early time points as the contrast is the sharpest here.

8. Export images as high-resolution TIFF files via a USB-A pen drive.

3-min timepoint | 6-min timepoint | 10-min timepoint

D – Digestion
L – Ligation

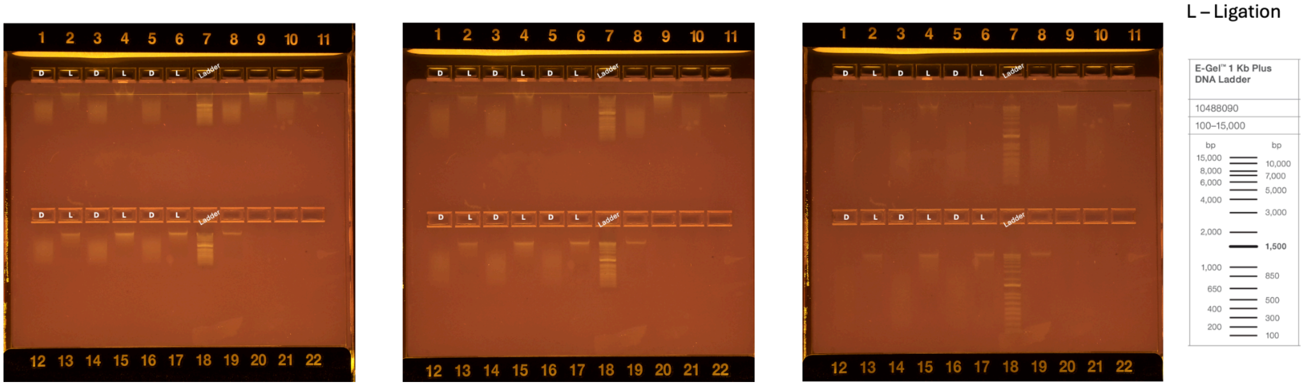
