## Supplementary Protocol 2 for "Whole-genome 3D architectural screen reveals modulators of brain DNA structure"

### Easy Dip-C Protocol

Updated Apr 15, 2026

#### Important Notes

Use **swinging-bucket centrifuge** to minimize cell loss.

When handling single cells, take every precaution to **avoid cross-contamination** between cells and between libraries.

Prepare the following reagents accordingly.

- Prepare **1.5 M sucrose** (recipe below for 40 mL):
  - 20.538 g sucrose (Sigma 84097; final: 1.5 M, 51.3% w/v)
  - Add water (ThermoFisher 10977023) so that the final volume is 40 mL (corresponding to ~27.1 mL water added)

- Prepare **Nuclei Isolation Medium** (recipe below for 45 mL; note that the commonly used Tris is replaced with HEPES):
  - 36.26 mL water (ThermoFisher 10977023)
  - 7.5 mL **1.5 M sucrose** (final: 250 mM, 8.56% w/v)
  - 562.5 µL 2 M KCl (ThermoFisher AM9640G; final: 25 mM)
  - 450 µL 1 M HEPES pH 7.5 (ThermoFisher 15630080; final: 10 mM)
  - 225 µL 1 M $MgCl_2$ (ThermoFisher AM9530G; final: 5 mM)
  - Vortex to mix. Store indefinitely at 4 C as shared lab stock. NOTE: Discard when it grows white stuff.

- Freshly prepare **1 mM DTT**:
  - 1 mL water (ThermoFisher 10977023)
  - 1 µL 1 M DTT (Sigma 646563; if unopened, each glass ampule can be stored indefinitely at room temperature; once opened, store indefinitely at –20 C)
  - Vortex to mix.

- Freshly prepare **Nuclei Isolation Buffer without Triton** (6 mL per sample; recipe below for 6 mL):
  - 6 mL **Nuclei Isolation Medium 1**
  - 6 µL 1 mM DTT (final: 1 µM)
  - Protease inhibitor (either 0.12 tablets of cOmplete EDTA-free (Roche 04693132001) (final: 1 tablet/50 mL), or 60 µL of 100 X Halt Protease Inhibitor Cocktail (ThermoFisher 78429) (final: 1 X)). NOTE (as of Nov 1, 2024): There is no evidence that this inhibitor makes any difference, while it is relatively expensive and often back-ordered. Therefore, no need to add it when practicing; only use it for important samples.
  - Vortex to mix. Chill on ice.

- Freshly prepare **Nuclei Isolation Buffer with Triton** (2 mL per sample; recipe below for 2 mL):
  - 2 mL **Nuclei Isolation Buffer without Triton**
  - 20 µL 10% Triton X-100 (Sigma 93443; final: 0.1%)

    - Vortex to mix. Chill on ice.

- Prepare **Iodixanol Medium** (recipe below for 45 mL):
    - 30.08 mL water (ThermoFisher 10977023)
    - 7.5 mL **1.5 M sucrose** (final: 250 mM)
    - 3.375 mL 2 M KCl (ThermoFisher AM9640G; final: 150 mM)
    - 2.7 mL 1 M Tris pH 8.0 (ThermoFisher AM9855G) (final: 60 mM)
    - 1.35 mL 1 M $MgCl_2$ (ThermoFisher AM9530G; final: 30 mM)
    - Vortex to mix. Store indefinitely at 4 C as shared lab stock.

- Prepare **50% Iodixanol in Medium** (recipe below for 45 mL):
    - 37.5 mL 60% Iodixanol (Sigma D1556-250ML; also known as OptiPrep) (final: 50%)
    - 7.5 mL **Iodixanol Medium**
    - Vortex to mix. Store indefinitely at 4 C as shared lab stock.

- Prepare **29% Iodixanol in Medium** (recipe below for 45 mL):
    - 21.75 mL 60% Iodixanol (Sigma D1556-250ML; also known as OptiPrep) (final: 50%)
    - 23.25 mL **Iodixanol Medium**
    - Vortex to mix. Store indefinitely at 4 C as shared lab stock.

- Freshly prepare **2% PFA in PBS** (recipe below for 12 mL):
    - 11.25 mL PBS (Invitrogen 10010023)
    - 750 µL 32% PFA (EMS 15714; if unopened, each glass ampule can be stored indefinitely at room temperature; **once opened, store at 4 C for 4 weeks**)
    - Gently invert to mix. Chill on ice.

- Freshly prepare **1% BSA in PBS** (recipe below for 10 mL):
    - 9 mL PBS (Invitrogen 10010023)
    - 1 mL 10% BSA in PBS (Miltenyi Biotec 130-091-376)
    - Vortex to mix. Chill on ice.

- Prepare **1% SDS in water** (recipe below for 10 mL):
    - 1 mL 10% SDS (Sigma 71736)
    - 9 mL water (ThermoFisher 10977023)

- Freshly prepare **Digestion Mix** (recipe below for 1 reaction):
    - 9 µL 10 X rCutSmart (NEB B9200)
    - 7 µL water (ThermoFisher 10977023)
    - 8 µL 5 U/uL MboI (NEB R0147L)
    - 4 µL 10 U/uL NlaIII (NEB R0125L)
    - Pipette to mix.

- Freshly prepare **Ligation Mix** (recipe below for 92 uL, 1 reaction):
    - 45.5 µL water (ThermoFisher 10977023)
    - 45 µL NEBNext Ultra II Ligation Master Mix (from NEB E7595L)
    - 1.5 µL NEBNext Ligation Enhancer (from NEB E7595L)
    - Pipette to mix.

- Freshly prepare Vazyme based **Ligation Mix** (recipe below for 8.2 mL):
    - 7 mL 2X Rapid ligation buffer (from Vazyme N103-01)
    - 1.2 mL T4 DNA ligase (from Vazyme N103-01)

- Prepare **Dip-C Lysis Buffer** (recipe below for 50 mL):
    - 46.45 mL water (ThermoFisher 10977023)
    - 1.25 mL 1 M DTT (Sigma 646563; if unopened, each glass ampule can be stored indefinitely at room temperature; once opened, store indefinitely at –20 C) (final: 25 mM)
    - 1 mL 1 M Tris pH 8.0 (ThermoFisher AM9855G) (final: 20 mM)
    - 750 µL 10% Triton X-100 (Sigma 93443) (final: 0.15%)
    - 250 µL **100 µM Carrier ssDNA,** TCAGGTTTTCCTGAA (final: 500 nM)
    - 200 µL 5 M NaCl (ThermoFisher AM9760G) (final: 20 mM)
    - 100 µL 0.5 M EDTA (ThermoFisher AM9260G) (final: 1 mM)
    - Vortex to mix. Aliquot into 2 mL per tube (25 tubes) and store indefinitely at –20 C as shared lab stock.

- Prepare **60 mg/mL Qiagen Protease (QP)**:
    - 1 vial (7.5 AU) of Qiagen Protease (Qiagen 19157)
    - 2.78 mL water (ThermoFisher 10977023)
    - Vortex to mix. Aliquot into ~500 µL per tube and store indefinitely at 4 C as shared lab stock.

- Freshly prepare **Dip-C Lysis Mix** (recipe below for 10 mL):
    - 10 mL **Dip-C Lysis Buffer** (can thaw at 37 C and cool to room temperature)
    - 2.5 µL **60 mg/mL Qiagen Protease** (final: 15 µg/mL)
    - Vortex to mix.

- Prepare **Transposition Buffer** (recipe below for 50 mL):
    - 39.06 mL water (ThermoFisher 10977023)
    - 10 mL 50% PEG 8000 (Hampton Research HR2-535) (final: 10%)
    - 625 µL 1 M TAPS pH 8.5 (Boston Bio Products BB-2375) (final: 12.5 mM)
    - 312.5 µL 1 M $MgCl_2$ (ThermoFisher AM9530G) (final: 6.25 mM)
    - Vortex vigorously (because PEG is viscous) to mix. Aliquot into 2 mL per tube, and store indefinitely at –20 C as shared lab stock, labeled “TB”.

- Prepare **Stop Buffer** (recipe below for 50 mL):
    - 42.45 mL water (ThermoFisher 10977023)
    - 4.5 mL 0.5 M EDTA (ThermoFisher AM9260G) (final: 45 mM)
    - 3 mL 5 M NaCl (ThermoFisher AM9760G) (final: 300 mM)
    - 50 µL 10% Triton X-100 (Sigma 93443) (final: 0.01%)
    - Vortex to mix. Aliquot into 2 mL per tube (25 tubes) and store indefinitely at –20 C as shared lab stock.

- Freshly prepare **Stop Mix** (recipe below for 1 mL):
    - 1 mL **Stop Mix** (can thaw at 37 C and cool to room temperature)
    - 8 µL **60 mg/mL Qiagen Protease** (final: 480 µg/mL, Qiagen 19157)
    - Vortex to mix.

- Freshly prepare **PCR Mix** (recipe below for 5mL) in a regular 15 mL tube:
  - 2.242 mL Q5 Reaction Buffer (NEB M0491S) (might have white precipitate when thawed for the first time; dissolve by heating with palm and vortexing)
  - 2.242 mL Q5 High GC Enhancer (NEB M0491S)
  - 268 μL 10 mM (each) dNTP mix (Vazyme P031-02 or NEB N0447S) (final: 538 μM (each))
  - 26.8 μL 1 M $MgCl_2$ (ThermoFisher AM9530G) (final: 5.38 mM)
  - 112 μL 20 mg/mL BSA (NEB B9000S) (final: 448 ug/mL)
  - 112 μL 2 U/μL Q5 DNA Polymerase (NEB M0491S) (final: 0.0448 U/μL)
  - Vortex to mix.

### Nuclei Isolation

1. Chill a Dounce homogenizer (Sigma D8938 for 2 mL, D9063 for 7 mL, D9938 for 15 mL, D9188 for 40 mL, D0189 for 100 mL) on ice.

2. Add 2 mL (please adjust volume according to the size of the sample) ice-cold **Nuclei Isolation Buffer with Triton** to the homogenizer.

3. Dounce the tissue with 5 strokes of the loose pestle (A), and 15 strokes of the tight pestle (B).

4. (Updated about discarding a portion if needed) Transfer the homogenate to a tube. Discard a portion if there is too much tissue (for example, 90% of a whole brain, 50% of a cerebellum).

5. Centrifuge at 100 g for 8 min at 4 C.

6. Carefully remove supernatant without disrupting the soft pellet. Resuspend in 2 mL **Nuclei Isolation Buffer without Triton**.

7. Centrifuge at 100 g for 8 min at 4 C.

8. Carefully remove supernatant without disrupting the soft pellet. Resuspend in 2 mL **Nuclei Isolation Buffer without Triton**. NOTE: If performing Optional: Gradient Centrifugation, resuspend in only ~500 μL (with a goal of obtaining 450 μL after filtering).

9. Filter. You can use either filters that fit on a 5 mL tube (such as Corning 352235 for 35 μm), or filters that fit on a 50 mL tube (such as Corning 352340 or Fisherbrand 22-363-547 for 40 μm). Please record the filter choice.

### Optional: Gradient Centrifugation

10. Freshly prepare **25% Solution** (recipe below for 900 uL):
    a. 450 μL filtered cells

    b. 450 µL **50% Iodixanol in Medium** (final: 25% iodixanol)
    c. Pipette to mix. NOTE: The 1:1 volume ratio must be exact.

11. In a new 2 mL tube, add 900 µL **29% Iodixanol in Medium**.

12. Carefully layer 900 µL **25% Solution** on top of the 900 uL **29% Iodixanol in Medium** by pipetting very slowly onto the side of the tube, without mixing the two layers.

13. Centrifuge at 13,500 g for 20 min at 4 C. NOTE: Use fixed-angle centrifuge for this step, because our swinging-bucket cannot achieve this speed.

14. Carefully remove the supernatant (including the floating white layer). Resuspend in 2 mL **Nuclei Isolation Buffer without Triton**.

### Fixation

15. To every 1 mL of cells, add 66.7 µL 32% PFA (EMS 15714; if unopened, each glass ampule can be stored indefinitely at room temperature; **once opened, store at 4 C for 4 weeks**) (final: 2%). NOTE: Tan typically does this in a 50 mL tube.

16. Rotate at RT for 10 min.

17. Freshly prepare **1% BSA in PBS** (recipe below for 10 mL):
    a. 9 mL PBS (Invitrogen 10010023)
    b. 1 mL 10% BSA in PBS (Miltenyi Biotec 130-091-376)
    c. Vortex to mix. Chill on ice.

18. To every 1 mL of cells, add 1 mL ice-cold **1% BSA in PBS**. Invert to mix.

19. Centrifuge at 1000 g for 5 min at 4 C.

20. Remove supernatant. Resuspend in 1 mL ice-cold **1% BSA in PBS**.

21. Count, and aliquot to ~1 million cells per tube. NOTE: No need to count if you know roughly the total cell number (e.g., 20–40 million per adult mouse cerebellum, 5–10 million per mouse cortex, 1 million per mouse's hippocampus).

22. Centrifuge at 1000 g for 5 min at 4 C.

23. Remove supernatant. PAUSE POINT: Fixed cells can be stored indefinitely at –80 C.

### Optional: Flow Sorting Fluorescent Cells

If needed, fluorescently labeled cell populations (e.g., vDip-C or other KASH tags, Sun1 tag, H2B tag, or NeuN+) can be isolated at this step.

24. Freshly prepare **300 uM DAPI in PBS**:
    a. 100 µL PBS (ThermoFisher 10010023) (final: 1X)
    b. 2.1 µL **14.3 mM (5 mg/mL) DAPI** (final: 300 uM)
    c. Vortex to mix.

25. Freshly prepare **300 nM DAPI in PBS** (recipe below for 1 mL):
    a. 1 mL PBS (ThermoFisher 10010023) (final: 1X)
    b. 1 uL **300 µM DAPI in PBS** (final: 300 nM)
    c. Vortex to mix.

26. Resuspend cells in 1 mL (volume depends on the number of cells; 5 mL per mouse cerebellum) **300 nM DAPI in PBS**. Transfer to a FACS tube. PAUSE POINT: Cells can be stored indefinitely at –80 C.

27. Among DAPI singlets, sort desired cells into **1% BSA in PBS**.

28. Centrifuge at 1000 g for 5 min at 4 C.

29. Remove supernatant. PAUSE POINT: Fixed cells can be stored indefinitely at –80 C.

### Chromatin Conformation Capture (3C/Hi-C)

#### Permeabilization

1. Thaw up to 1 million cells on ice.

2. Resuspend in 20 µL water (ThermoFisher 10977023).

3. Add 22 µL **1% SDS**. Pipette to mix.

4. Incubate at 62 C for 10 min.

5. Add 22 µL 10% Triton X-100 (Sigma 93443). Pipette to mix.

6. Incubate at 37 C for 15 min.

#### Digestion

7. Freshly prepare **Digestion Mix** (recipe below for 28 uL):
    a. 9 µL 10 X rCutSmart (NEB B9200)
    b. 7 µL water (ThermoFisher 10977023)
    c. 8 µL 5 U/uL MboI (NEB R0147L)
    d. 4 µL 10 U/uL NlaIII (NEB R0125L)
    e. Pipette to mix.

8. Add 28 µL **Digestion Mix**. Pipette to mix.

9. Incubate at 37 C for 1 h.

10. Incubate at 65 C for 20 min.

11. Take 10 µL as **Digestion QC** (can store at –20 C).

#### Ligation

12. Wait for the reaction to cool down to room temperature (~15 min after removal from 65 C).

13. Freshly prepare **Ligation Mix** (recipe below for 92 uL, 1 reaction):
    a. 45.5 µL water (ThermoFisher 10977023)
    b. 45 µL NEBNext Ultra II Ligation Master Mix (from NEB E7595L)
    c. 1.5 µL NEBNext Ligation Enhancer (from NEB E7595L)
    d. Pipette to mix.

14. Add 92 µL **Ligation Mix** to the reaction. Pipette to mix.

15. Incubate at room temperature for 15 min.

16. Take 10 µL as **Ligation QC** (can store at –20 C).

17. Freshly prepare **300 uM DAPI in PBS**:
    a. 100 µL PBS (ThermoFisher 10010023) (final: 1X)
    b. 2.1 µL **14.3 mM (5 mg/mL) DAPI** (final: 300 uM)
    c. Vortex to mix.

18. Freshly prepare **300 nM DAPI in PBS** (each reaction consumes 1 mL; recipe below for 1 mL):
    a. 1 mL PBS (ThermoFisher 10010023) (final: 1X)
    b. 1 µL **300 uM DAPI in PBS** (final: 300 nM)
    c. Vortex to mix.

19. Add 1 mL **300 nM DAPI in PBS** to the ligated nuclei. Pipette to mix. PAUSE POINT: Ligated nuclei can be stored indefinitely at –80 C.

#### Quality Control

20. Centrifuge **Digestion QC** and **Ligation QC** at 1000 g for 5 min at 4 C.

21. Remove supernatant. Resuspend each in 95 µL PBS (ThermoFisher 10010023).

22. Add 5 µL Proteinase K (NEB P8107S) to each. Vortex to mix.

23. Incubate at 65 C for 1 h.

24. Purify the DNA with a column (Zymo D4013), using 5 times volume (500 µL) of DNA Binding Buffer and eluting into 6 µL of TE (Invitrogen AM9849).

25. Use 1 µL for quantification using Qubit HS DNA kit and dilute the rest of volume with 1X TE to have a final concentration of (less than) 5 ng/µL and submit to bioanalyzer.

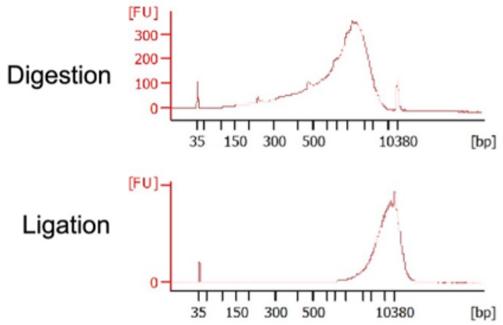


### Optional: Bulk Library Preparation

At this stage, bulk Dip-C libraries can be directly prepared from ligated cells, similar to how **Ligation QC** is purified.

26. Take a desired portion of ligated nuclei (for example, half). Centrifuge at 1000 g for 5 min at 4 C. NOTE: Speed can be increased to 3000 g if one is worried about losing the pellet when cell numbers are low.

27. Remove supernatant. Resuspend in 95 uL PBS (ThermoFisher 10010023).

28. Add 5 µL Proteinase K (NEB P8107S). Vortex to mix.

29. Incubate at 65 C for 1 h.

30. Purify the DNA with a column (Zymo D4013), using 5 times volume (500 µL) of DNA Binding Buffer and eluting into a desired volume (for example, 50 µL) of TE (Invitrogen AM9849).

31. Use 1 µL for quantification using Qubit HS DNA kit.

32. Use a desired amount (for example, 100 ng) for library preparation with a half reaction of Vazyme TruePrep Flexible (Vazyme TD504; a cheaper version of Illumina DNA Prep).

33. Remove shorter fragments with 2 times of 0.55 X size selection. NOTE: This is more stringent than size selection of single cells. Alternatively, only do 0.6 X left side size selection only once.

### Whole-genome Amplification

While this section can be easily done by hand, automated liquid handling can greatly accelerate the procedure.

### First Time: Determining Optimal Tn5 Concentration

When using a new tube of Tn5 transposome (Illumina TDE1, or TTE Mix V50 from Vazyme TD501-01) and/or a new type of sample, determine the optimal Tn5 concentration by performing whole-genome amplification (on cells or cell lysate) with varying amounts of Tn5, and identifying the concentration that leads to a flat Bioanalyzer profile.

This is also a good positive control (or practice reaction) for new users.

1. Thaw ligated cells on ice. If the tube contains ~1 million cells in 1 mL, the cell density will be ~1,000 cells/μL.

2. Pipette to mix. Transfer 1 μL (~1,000 cells) to a PCR tube.

3. Add 100 μL **Dip-C Lysis Mix** (see Flow Sorting and Lysis). Vortex to mix.

4. Lyse the cells into **Dip-C Lysate**. The lysate contains ~10 lysed cells/μL. PAUSE POINT: **Dip-C Lysate** can be stored indefinitely at –80 C.

5. Using 1 μL **Dip-C Lysate** per reaction, perform whole-genome amplification with varying amounts of Tn5 transposome (e.g., 30, 10, 3, 1, 0.3, 0.1, 0.03, 0.01 μL per 850 μL **Transposition Buffer**). Namely, following the sections Transposition, Stopping, Amplification, and Purification but eluting into 6 μL TE (Invitrogen AM9849) per reaction.

6. Measure DNA concentration with Qubit.

7. Measure DNA length distribution with Bioanalyzer HS DNA (if needed, dilute to less than 5 ng/μL).

8. Identify optimal Tn5 concentration as the one that leads to the **flattest** Bioanalyzer profile (a little higher on the long side is the most ideal). Perform a finer optimization if needed.

### Flow Sorting and Lysis

1. Thaw ligated cells on ice. Transfer to a 5 mL FACS tube and filter if necessary. If sorting for the first time, filter the cells with 35 μm (Corning 352235: the blue-capped FACS tubes) to avoid clumping the machine.

2. Prepare **Dip-C Lysis Buffer** (recipe below for 50 mL):
    a. 46.45 mL water (ThermoFisher 10977023)
    b. 1.25 mL 1 M DTT (Sigma 646563; if unopened, each glass ampule can be stored indefinitely at room temperature; once opened, store indefinitely at –20 C) (final: 25 mM)
    c. 1 mL 1 M Tris pH 8.0 (ThermoFisher AM9855G) (final: 20 mM)
    d. 750 μL 10% Triton X-100 (Sigma 93443) (final: 0.15%)
    e. 250 μL **100 μM Carrier ssDNA** (final: 500 nM)
    f. 200 μL 5 M NaCl (ThermoFisher AM9760G) (final: 20 mM)
    g. 100 μL 0.5 M EDTA (ThermoFisher AM9260G) (final: 1 mM)

    h. Vortex to mix. Aliquot into 2 mL per tube (25 tubes) and store indefinitely at –20 C as shared lab stock.

3. Prepare **60 mg/mL Qiagen Protease (QP)**:
    a. 1 vial (7.5 AU) of Qiagen Protease (Qiagen 19157)
    b. 2.78 mL water (ThermoFisher 10977023)
    c. Vortex to mix. Aliquot into ~500 uL per tube and store indefinitely at 4 C as shared lab stock.

4. Freshly prepare **Dip-C Lysis Mix** (recipe below for 1 mL):
    a. 1 mL **Dip-C Lysis Buffer** (can thaw at 37 C and cool to room temperature)
    b. 0.25 µL **60 mg/mL Qiagen Protease** (final: 15 µg/mL)
    c. Vortex to mix.

5. Add 1 µL **Dip-C Lysis Mix** per well to a 384-well plate.

6. Seal with film (the transparent 'B' type: Bio-Rad MSB1001; or the metallic 'A' type: Bio-Rad MSF1001) and roller (Bio-Rad MSR0001). PAUSE POINT: **Dip-C Lysis Mix** can be stored on ice for a few hours.

7. Sort using the 100 µm chip.

8. Sort a single cell per well based on the DAPI signal. Seal tightly with the transparent 'B' film (Bio-Rad MSB1001; NOT the 'F' film) to avoid evaporation. NOTE: This is the only step where the transparent 'B' film (Bio-Rad MSB1001) is required, because of the low volume (1 uL) and relatively high incubation temperature (50 C and 70 C). In all other steps, the metallic 'F' film (Bio-Rad MSF1001) can be used instead if you prefer its handling.

9. Immediately vortex at the FACS facility to spread out the lysis buffer to better capture the cells. Centrifuge at 1000 g for 1 minute (this can be done back in our lab). PAUSE POINT: Plates can be stored on ice overnight.

NOTE: Vortexing immediately and thoroughly at the FACS facility might be critical for 384-well plates.

10. Lyse the cells:
    a. 50 C for 1 h
    b. 70 C for 15 min
    c. 4 C forever. PAUSE POINT: Plates can be stored at 4 C overnight, or for a few months at –80 C. Store them in one of the six plate racks at the top of the –80 C. NOTE: The 'B' film might peel a little, or become difficult to thaw and peel off, when stored for a long time at –80 C. Feel free to swap it with the 'F' film (Bio-Rad MSF1001) before storing at –80 C.

### Transposition

11. Thaw plates of lysed cells. Wipe off condensation from the seal.

12. Spin down the plate (500 g at 4 C briefly).

13. Prepare **Transposition Buffer** (recipe below for 50 mL):

    a. 39.06 mL water (ThermoFisher 10977023)
    b. 10 mL 50% PEG 8000 (Hampton Research HR2-535) (final: 10%)
    c. 625 µL 1 M TAPS pH 8.5 (Boston Bio Products BB-2375) (final: 12.5 mM)
    d. 312.5 µL 1 M $MgCl_2$ (ThermoFisher AM9530G) (final: 6.25 mM)
    e. Vortex vigorously (because PEG is viscous) to mix. Aliquot into 2 mL per tube and store indefinitely at –20 C as shared lab stock, labeled “TB”.

14. Freshly prepare **Transposition Mix** (recipe below for 2 mL, suitable for a 384-well plate with 30% or 464 µL overhead):
    a. 2 mL **Transposition Buffer** (can thaw at 37 C and cool to room temperature)
    b. Tn5 transposome (1.4 µL Vazyme TTE Mix V50 from TD501)
    c. Pipette to mix. NOTE: Do not vortex!

15. Add 4 µL **Transposition Mix** per well.

16. Vortex the plate thoroughly (at 12 roughly evenly spaced points along the plate) and spin down the plate.

17. Transpose the genome:
    a. 55 C for 10 min
    b. 4C forever. PAUSE POINT: Plates can be stored at 4 C overnight.

### Stopping

18. Prepare **Stop Buffer** (recipe below for 50 mL):
    a. 42.45 mL water (ThermoFisher 10977023)
    b. 4.5 mL 0.5 M EDTA (ThermoFisher AM9260G) (final: 45 mM)
    c. 3 mL 5 M NaCl (ThermoFisher AM9760G) (final: 300 mM)
    d. 50 µL 10% Triton X-100 (Sigma 93443) (final: 0.01%)
    e. Vortex to mix. Aliquot into 2 mL per tube (25 tubes) and store indefinitely at –20 C as shared lab stock.

19. Freshly prepare **Stop Mix** (recipe below for 1 mL):
    a. 1 mL **Stop Mix** (can thaw at 37 C and cool to room temperature)
    b. 1.667 µL **60 mg/mL Qiagen Protease** (final: 100 µg/mL)
    c. Vortex to mix.

20. Add 1 µL **Stop Mix** per well.

21. Vortex the plate thoroughly (at 12 roughly evenly spaced points along the plate) and spin down the plate.

22. Stop transposition:
    a. 50 C for 40 min
    b. 70 C for 20 min
    c. 4 C forever. PAUSE POINT: Plates can be stored at 4 C overnight. NOTE: Do not store at –20 C, which will greatly reduce the yield!

#### Amplification

23. (Updated recipe to 3 mL, which is suitable for one 384-well plate or four 96-well plates, with 30% or 696 µL overhead) Freshly prepare **PCR Mix** (recipe below for 3 mL, suitable for a 384-well plate with 30% or 696 uL overhead) in a regular 15 mL tube:
    a. 1.345 mL Q5 Reaction Buffer (NEB M0491S) (might have white precipitate when thawed for the first time; dissolve by heating with palm and vortexing)
    b. 1.345 mL Q5 High GC Enhancer (NEB M0491S)
    c. 161 µL 10 mM (each) dNTP mix (Vazyme P031-02 or NEB N0447S) (final: 538 µM (each))
    d. 16.1 µL 1 M $MgCl_2$ (ThermoFisher AM9530G) (final: 5.38 mM)
    e. 67.2 µL 20 mg/mL BSA (NEB B9000S) (final: 448 ug/mL)
    f. 67.2 µL 2 U/µL Q5 DNA Polymerase (NEB M0491S) (final: 0.0448 U/µL)
    g. Vortex to mix.

24. Add 6 µL **PCR Mix** per well.

25. Use multi-channel pipette or Bravo (see Appendix B: Adding Nextera Primers with Bravo) to add 1 µL **6.25 µM (each) Nextera Primer Mix** (thaw at room temperature, and centrifuge at 3,000 g for 5 min) per well. NOTE: Set speed (both aspirating and dispensing speeds) of the motorized pipette (Eppendorf) to the slowest (1) to pipette such low volume (1 uL) reliably.

26. Vortex the plate thoroughly (at 12 roughly evenly spaced points along the plate) and spin down the plate.

27. Use 16 cycles for amplification:
    a. 4 C for 3 min
    b. 72 C for 3 min
    c. 98 C for 20 s
    d. 16 cycles of 98 C for 10 s, 62 C for 1 min, 72 C for 2 min
    e. 72 C for 5 min
    f. 4 C forever. PAUSE POINT: Plates can be stored at 4 C overnight, or for a few months at –80 C.

#### Purification

28. Pool reactions from the whole plate into 5 times volume of DNA Binding Buffer (Zymo D4004-1-L; needs to buy many bottles) (6 mL per 96-well plate, 24 mL per 384-well plate). You can either use a multi-channel pipette, or centrifuge (50 g for 5 min) the plate up-side-down into a V-bottom plate reservoir (ClickBio CBVBLOK200-1) pre-filled with DNA Binding Buffer. Vortex to mix.

NOTE: The V-bottom plate reservoir (ClickBio CBVBLOK200-1) does not fit our centrifuge, and will get stuck if put directly; to avoid this, put the plate reservoir on top of a stack of 3 empty 384-well PCR plates.

NOTE: You MUST visually examine the V-bottom reservoir for **cracks** before use, otherwise liquid can be entirely lost. PAUSE POINT: Pooled reaction mixed with Binding Buffer can be stored indefinitely at –20 C.

29. Purify a portion of the mixture with columns (Zymo D4013) and elute into 1X TE (Invitrogen AM9849). Tan typically purifies 7.2 mL of mixture by loading 3 columns (each can hold 800 uL) 3 times, and elute into 200 uL TE.

NOTE: Please **write the index** (such as “UDI G”) on every tube to avoid confusion.

30. Measure DNA concentration with Qubit, using the 1x dsDNA HS kit.

31. Measure DNA length distribution with Bioanalyzer, using a High Sensitivity DNA chip (dilute to less than 5 ng/uL if needed). NOTE: Adjust future Tn5 concentration if length is not optimal.

#### Size Selection

This section removes shorter fragments with 2 times of 0.6 X size selection.

32. Combine 100 µL of purified (un-sized) library with 60 µL of SPRIselect beads (Beckman B23318 or alternatives such as Vazyme). Vortex to mix. Incubate at room temperature for 5 min.

NOTE: Mix beads thoroughly before use. Beckman B23318 is stored at room temperature; but alternatives are usually stored at 4 C, which must be warmed to room temperature before use.

33. Spin down. Put the tube on a magnetic rack (for example, Thermo 12321D). Wait 5 min until all beads (brown) settle to the bottom.

34. Remove supernatant without disturbing the beads.

35. Carefully add 500 µL 80% EtOH (200 or 180 µL if using smaller tubes) without disturbing the beads.

NOTE: Do NOT disturb beads because they take a very long time to settle in EtOH, if at all.

36. Remove supernatant without disturbing the beads.

37. Carefully add 500 µL 80% EtOH (200 µL if using smaller tubes and magnetic racks) without disturbing the beads.

38. Remove supernatant without disturbing the beads. Dry the beads by waiting ~5 min. Use a small pipette to remove any residual EtOH.

39. Remove the tube from the magnetic rack. Resuspend beads in ~105 µL TE (adjust the exact volume based on how much volume will allow you to confidently transfer 100 µL in the end without transferring any beads). Vortex to mix. Incubate at room temperature for 2 min.

40. Spin down. Put the tube on the magnetic rack. Wait 5 min until all beads (brown) settle to the bottom.

41. Carefully transfer 100 µL supernatant to a new tube without transferring any beads.

NOTE: If brown color (beads) appears at the end of the pipette tip when transferring, transfer it back and try again more carefully and/or with less volume.

NOTE: The above steps finish the first round of size selection. Steps below perform another round, which **differs only by the elution volume** (~100 μL above, and ~25 μL below).

42. Combine the 100 μL supernatant with 60 μL of SPRIselect beads (Beckman B23318 or alternatives such as Vazyme). Vortex to mix. Incubate at room temperature for 5 min.

NOTE: Mix beads thoroughly before use. Beckman B23318 is stored at room temperature; but alternatives are usually stored at 4 C, which must be warmed to room temperature before use.

43. Spin down. Put the tube on a magnetic rack (for example, Thermo 12321D). Wait 5 min until all beads (brown) settle to the bottom.

44. Remove supernatant without disturbing the beads.

45. Carefully add 500 μL 80% EtOH (200 or 180 μL if using smaller tubes) without disturbing the beads.

NOTE: Do NOT disturb beads because they take a very long time to settle in EtOH, if at all.

46. Remove supernatant without disturbing the beads.

47. Carefully add 500 μL 80% EtOH (200 μL if using smaller tubes and magnetic racks) without disturbing the beads.

48. Remove supernatant without disturbing the beads. Dry the beads by waiting ~5 min. Use a small pipette to remove any residual EtOH.

49. Remove the tube from the magnetic rack. Resuspend beads in ~28 μL TE (adjust the exact volume based on how much volume will allow you to confidently transfer 25 μL in the end without transferring any beads). Vortex to mix. Incubate at room temperature for 2 min.

50. Spin down. Put the tube on the magnetic rack. Wait 5 min until all beads (brown) settle to the bottom.

51. Carefully transfer 25 μL supernatant to a new tube without transferring any beads.

NOTE: If brown color (beads) appears at the end of the pipette tip when transferring, transfer it back and try again more carefully and/or with less volume.

NOTE: This is the **final library**.

52. Measure DNA concentration with Qubit. Record the percentage yield after size selection: ~20% indicates optimal transposition (higher indicates insufficient transposition, lower indicates excessive transposition). As seen above, we typically size select 100 uL of DNA and elute into 25 uL TE for the final library.

### Appendix A: E-Gel Quality Control

If many samples need to be checked for quality control, one option is to run agarose gels. We have precast, prestained E-Gel™ Agarose Gels with SYBR™ Safe DNA Gel Stain (ThermoFisher A42100 which has 10 lanes, A42347/double-comb which has 11*2 = 22 lanes; ThermoFisher G820801 with 24*2 = 48 lanes; 96-lane gels and more).

1. Purify DNA as described in *Quality Control*.

2. Thaw 1X E-gel loading buffer (ThermoFisher 10482055) and the ladder of choice. (https://assets.thermofisher.com/TFS-Assets/BID/Reference-Materials/e-gel-ladder-reference-card.pdf)
    a. Use the E-Gel™ **1 Kb Plus DNA Ladder** (size range: 100–15,000 bp) (ThermoFisher 10488090).
    b. Do NOT use the E-Gel™ **1 Kb Plus Express DNA Ladder** (size range: 100–5,000 bp) (ThermoFisher 10488091) as the ligation product exceeds the size range.

3. Prepare samples: load 50 ng of DNA (minimum) + 2 µL 1X E-gel loading buffer; make-up the total volume to 20 µL with water (ThermoFisher 10977023).

4. Undock the camera from the E-Gel™ Electrophoresis Device, place it aside. Open a gel packet and load the gel by clicking it into place in the electrophoresis device.

5. Load samples into the lanes. For each row of samples, ensure the ladder is loaded onto one lane (10 µL to be used; ThermoFisher 10488090).

6. Dock the camera onto the electrophoresis device, set the run (ensure the gel is chosen correctly on the interface – 1% Agarose gel with SYBR™ Safe).

7. The duration should display 26 min; however, capture images at 3-, 6-, 10-minute timepoints (via the user interface on the camera screen). NOTE: do not miss the early time points as the contrast is the sharpest here.

8. Export images as high-resolution TIFF files via a USB-A pen drive.

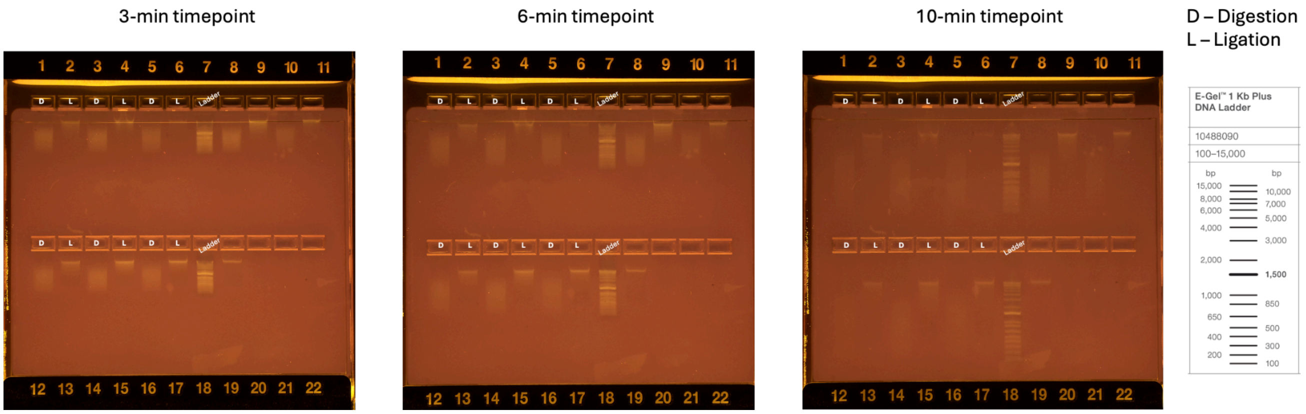
